# Identification and Pharmacological Targeting of Treatment-Resistant, Stem-like Breast Cancer Cells for Combination Therapy

**DOI:** 10.1101/2023.11.08.562798

**Authors:** Jeremy Worley, Heeju Noh, Daoqi You, Mikko M. Turunen, Hongxu Ding, Evan Paull, Aaron T. Griffin, Adina Grunn, Mingxuan Zhang, Kristina Guillan, Erin C. Bush, Samantha J. Brosius, Hanina Hibshoosh, Prabhjot S. Mundi, Peter Sims, Piero Dalerba, Filemon S. Dela Cruz, Andrew L. Kung, Andrea Califano

## Abstract

Tumors frequently harbor isogenic yet epigenetically distinct subpopulations of multi-potent cells with high tumor-initiating potential—often called Cancer Stem-Like Cells (CSLCs). These can display preferential resistance to standard-of-care chemotherapy. Single-cell analyses can help elucidate Master Regulator (MR) proteins responsible for governing the transcriptional state of these cells, thus revealing complementary dependencies that may be leveraged via combination therapy. Interrogation of single-cell RNA sequencing profiles from seven metastatic breast cancer patients, using perturbational profiles of clinically relevant drugs, identified drugs predicted to invert the activity of MR proteins governing the transcriptional state of chemoresistant CSLCs, which were then validated by CROP-seq assays. The top drug, the anthelmintic albendazole, depleted this subpopulation *in vivo* without noticeable cytotoxicity. Moreover, sequential cycles of albendazole and paclitaxel—a commonly used chemotherapeutic —displayed significant synergy in a patient-derived xenograft (PDX) from a TNBC patient, suggesting that network-based approaches can help develop mechanism-based combinatorial therapies targeting complementary subpopulations.

**Statement of significance:** Network-based approaches, as shown in a study on metastatic breast cancer, can develop effective combinatorial therapies targeting complementary subpopulations. By analyzing scRNA-seq data and using clinically relevant drugs, researchers identified and depleted chemoresistant Cancer Stem-Like Cells, enhancing the efficacy of standard chemotherapies.

## Introduction

Intratumor heterogeneity represents a major barrier in cancer treatment. Indeed, most tumors comprise co-existing, molecularly distinct subpopulations presenting non-overlapping drug sensitivities^1^. While some of the cells comprising them may represent genetically distinct subclones, a majority has emerged as representing the byproduct of pathophysiological epigenetic plasticity. In breast cancer (BRCA), for instance, there have been multiple reports of an isogenic Cancer Stem-like Cell (CSLC) subpopulation associated with differential expression of epigenetic regulators involved in controlling stemness programs, such as the BMI1, WNT, and NOTCH pathways^2–4^. CSLCs have been shown to display tumor-initiating capacity, expression of stem-cell markers, and resistance to common chemotherapeutics^5,6^, such as paclitaxel—a microtubule inhibitor and antimitotic widely used in the treatment of multiple malignancies, including breast cancer. Indeed, while frequently leading to initial tumor shrinkage, treatment with this drug is often followed by relapse and resistance. Indeed, it has been suggested that chemotherapy resistant breast CSLCs may regenerate the full heterogeneity of the tumor, as confirmed by limiting dilution assays^7,8^. Multiple non-mutually exclusive mechanisms of chemotherapy resistance have been proposed for CSLCs in breast and other tumors, including upregulation of multi-drug transporters, increased DNA damage repair, and better scavenging of ROS^9–11^. Taken together, these data suggest that breast CSLCs pose a fundamental challenge to achieving durable remissions in BRCA, especially in Triple Negative Breast Cancer (TNBC), where chemotherapy remains a cornerstone of treatment.

To gain insight into the molecular heterogeneity of breast cancer and to predict the sensitivity of individual subpopulations to clinically relevant drugs, we generated single-cell RNA sequencing (scRNA-seq) profiles of malignant cells isolated from biopsies of seven metastatic breast cancer patients. To enrich for cells with a stem-like phenotype—or CSLCs for simplicity—which may include only a very small fraction of tumor cells, we used fluorescence-activated cell sorting (FACS), with antibodies selected to purify malignant cells with a phenotype analogous to that of stem/progenitor cells in the normal mammary epithelium^12^. See **Fig. 1A** for an illustrative graphical workflow of this process.

**Figure 1.**
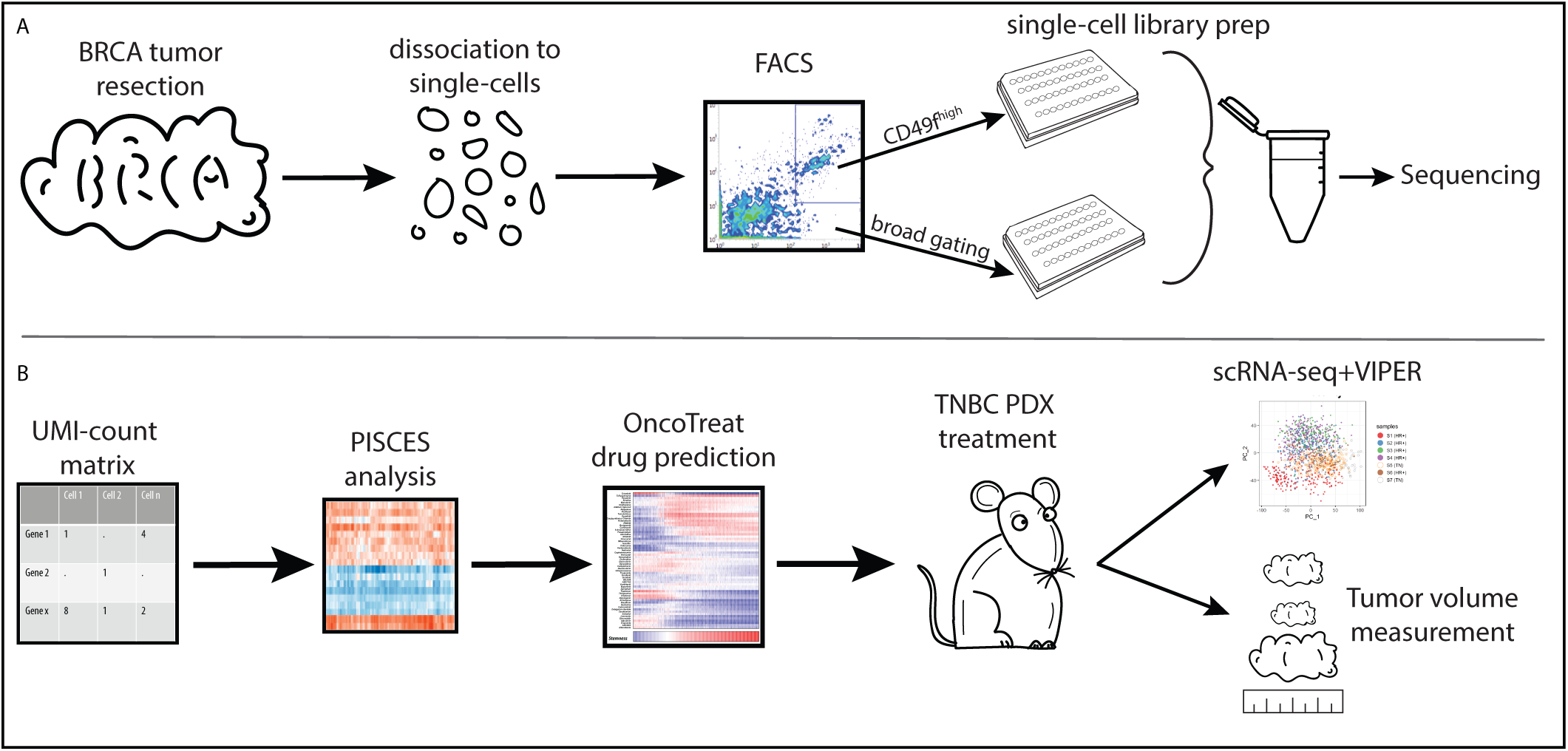
Overview of the workflow. **A.** The experimental workflow for generating scRNA-seq data from breast cancer cells from patient samples. FACS was used to enrich CSLCs. **B.** A systems biology approach to identifying a candidate drug targeting the CSLCs and subsequent experimental validations.

In previous studies, we have shown that highly sparse single scRNA-seq profiles, where >80% of the genes may produce no reads, can be transformed to fully populated protein activity profiles by the metaVIPER algorithm^13^—the single-cell adaptation of the extensively validated VIPER algorithm^14^. This is accomplished by measuring the activity of each regulatory and signaling protein based on the expression of its entire repertoire of transcriptional targets, akin to using a highly multiplexed, tissue-specific gene reporter assay. As a result, the most differentially active VIPER-inferred proteins are also enriched for Master Regulator (MR) proteins representing mechanistic determinants, via their target genes, of the associated transcriptional state.

MetaVIPER analysis of single cells isolated from the seven metastatic breast cancer patients accrued to the study—including five hormone receptor-positive (HR+) and two triple-negative (TNBC) tumors—effectively separated cells with a more stem-like vs. more differentiated transcriptional state, using a *stemness score* (*SS*) based on both established breast cancer stemness markers and CytoTRACE analysis^15^. Consistent with expectations, cells with the highest score (i.e., most stem-like) emerged as the most resistant to *in vivo* treatment with paclitaxel, while those with the lowest score (*i.e.*, most differentiated-like) were significantly depleted by the drug. This provided the molecular basis to identify and genetically/pharmacologically target candidate Master Regulators (MRs) of CSLC transcriptional state(s) identified by metaVIPER analysis.

We thus performed patient-by-patient analysis, using the VIPER algorithm to identify candidate MR proteins controlling the transcriptomic state of cells with the highest vs. lowest Stemness Score. Candidate MRs identified by the analysis were highly conserved across virtually all patients, independent of hormone receptor (HR) status, thus supporting the notion of a common CLSC MR signature. Indeed, >80% of the most significant VIPER-inferred activated and inactivated MRs were able to statistically significantly reprogram cells to a more differentiated or CSLC state, respectively, following their CRISPR-mediated silencing in a pooled CROP-seq^16^ assay in cell lines comprising both subtypes. We thus leveraged the OncoTreat algorithm^17^, which assesses the activity of MR proteins in drug vs. vehicle control-treated cells, to identify small molecule compounds capable of inverting the activity of the CLSC MR signature (MR-inverter drugs), thus potentially inducing differentiation or selective ablation. For this purpose, we leveraged gene expression profiles of BRCA cells—selected to faithfully recapitulate the CSLC MR signature—treated with a repertoire of 91 clinically relevant drugs, see **Fig. 1B** for an illustrative graphical workflow of these steps. Notably, OncoTreat-predicted drugs from either bulk^17–19^ or single-cell profiles^20,21^ have been extensively validated *in vivo* in prior studies.

Albendazole, a well-tolerated anthelmintic drug, emerged as the most statistically significant MR-inverter drug, yet at a concentration that was approximately ten-fold lower than its clinically tolerated dose; this was especially surprising since albendazole is not considered an anti-tumor drug. Based on these results, this drug was selected for experimental validation *in vivo*. Mice from a TNBC PDX model were treated with either albendazole or vehicle control for 14 days and compared to paclitaxel-treated animals. In contrast to paclitaxel, which caused highly significant increase of the CSLC to differentiated cell ratio, albendazole treatment induced equally dramatic yet opposite effects, suggesting that alternating treatment with the two drugs may abrogate the tumor-initiating potential of paclitaxel-resistant cells, while also preventing uncontrolled tumor growth. The strong rationale for combination-based, sequential therapy was confirmed by a preclinical study, where treatment with multiple cycles of albendazole and paclitaxel displayed superior anti-tumor activity compared to the corresponding monotherapies, resulting in a statistically significant synergistic effect.

## Results

### Intratumor heterogeneity in human breast carcinomas

Since patient-derived breast cancer tissues vary widely in size, cellularity, necrotic fraction, stromal infiltration, and overall quality, we used FACS to purify malignant cells using appropriate antibody combinations. Single cells isolated from these tumors were then processed to generate plate-based scRNA-seq profiles using an approach that combines elements of Smart-seq2^22^ and PLATE-seq^23^ (see STAR methods). This procedure, which allows sorting individual cancer cells into separate wells filled with lysis buffer for RNA-seq profiling, is especially effective in enriching for relatively rare subpopulations from fresh tumor tissue, since it effectively supports FACS-based cell isolation while removing debris and dead cells that may otherwise degrade the performance of other platforms. It was thus preferred at the time, despite its higher cost and complexity.

Fresh samples were obtained from two metastatic TNBC and five metastatic HR^+^ patients. To minimize post-resection transcriptional changes/drift, fresh samples were rapidly dissociated into a single-cell suspension (see STAR methods) and stained with DAPI, as well as ⍺-EpCAM, ⍺- CD49f, and Lin^-^ antibodies. EPCAM effectively distinguishes epithelial breast cancer cells from stromal subpopulations, whereas CD49f is known to be expressed at the highest levels in a subset of mammary epithelial cells acting as mammary repopulating units (MRUs) in transplantation assays^12^ and has been previously used to enrich for breast cancer cells with stem-like properties^24–26^. Starting from primary malignant tissues, we sorted live (DAPI^-^) epithelial (EpCAM^+^) cells into two distinct batches, including: (1) a first batch of unselected cancer cells (EPCAM^+^), representative of the full heterogeneity of the epithelial compartment, contributing ∼25% of the total cells in the analysis and (2) a second batch of epithelial cells with a phenotype characteristic of MRUs in the mouse mammary gland (EPCAM^+^, CD49f^high^), expected to be CSLCs-enriched^12,24–26^, contributing the remaining ∼75% of analyzed cells (**Fig. 1A, Suppl. Fig. S1**).

### Copy Number Variation Analysis

After NGS library generation and sequencing (see STAR methods), we performed several data pre-processing steps to ensure that subsequent analyses would be restricted to high-quality cancer cells. This included inference of somatic copy number alteration (CNA) assessment, using the Trinity CTAT Project inferCNV algorithm (https://github.com/broadinstitute/inferCNV) to exclude confounding effects from normal cells in the tumor microenvironment. Compared to cells representative of normal breast epithelium, most of the cells isolated from the seven patients presented clearly aberrant CNA structure, consistent with the high cellularity of metastatic samples (**Suppl. Fig. S2**). Interestingly, no intratumor CNA heterogeneity was detected by the analysis, suggesting that, at least from a copy number alteration perspective, the cells in these samples were clonally identical. However, as expected, the analysis showed significant inter-tumor CNA heterogeneity across the seven patients, especially between TNBC and HR+ samples.

### Protein activity-based analysis identifies a stem-like subpopulation

In addition to biological variation between tumors from different patients, substantial batch and biology-related effects may also challenge the analysis of single cells isolated from different samples. Batch effects can arise due to technical artifacts, such as changes in temperature or reagents between samples processed on different days, or liquid handling drift in multi-well plate assays. In addition, inter-patient CNA differences may also contribute to significant gene expression heterogeneity, which may confound the analysis. Indeed, while only a handful of genes in CNAs play a functional role in tumorigenesis, most of the genes in these amplicons may still produce substantial inter-patient bias at the gene expression level, even though the activity of their encoded proteins is ultimately buffered by the post-transcriptional autoregulatory logic of the cell. When combined with the high *gene dropout* rate of scRNA-seq profiles—where >75% of the genes may fail to be detected by even a single read—this limits the ability to perform detailed, quantitative analyses using traditional gene expression-based methodologies.

Various approaches to reduce noise and minimize gene dropout effects have been proposed^27,28^—such as metaCells^29^ and imputation-based^30^ methods—as well as normalization methods aimed at reducing batch effects^31,32^. These methodologies, however, may introduce artifacts that affect subsequent analyses. For instance, using metaCells may prevent identification of rare subpopulations, whose gene expression profile would be averaged with cells from molecularly distinct subpopulations, while normalization may reduce biologically relevant differences between samples. Most critically, generating a comprehensive repertoire of candidate molecular determinants of tumor cell state, potentially associated with differential expression of only a handful of genes, is quite challenging if the expression of most genes is undetectable in individual cells. Transcriptional regulators, which are critical in maintaining cell state/identity, are especially affected by such *gene dropout* issues because they can be functionally active even when expressed at very low levels.

To address these challenges, we leveraged the PISCES single-cell analysis pipeline^33^, which provides a systematic framework for protein activity-based analysis of single-cell data—from raw counts quality control to construction of gene regulatory networks, to the identification of MR proteins (see STAR methods). Specifically, PISCES leverages the metaVIPER^13^ algorithm to measure a protein’s differential activity based on the differential expression of its transcriptional targets, as inferred by the ARACNe^34^ algorithm. These algorithms have been extensively validated, showing low false positive rates (in the 20% – 30% range)^14,35,36^ and almost complete elimination of technical (i.e., non-biologically-relevant) batch effects. In particular, we have recently shown that metaVIPER protein activity measurements significantly outperform gene expression and even antibody-based measurements in single cells^20,37,38^, including based on large-scale CITE-seq assays^33^.

We used metaVIPER to infer protein activity of single cells isolated from breast cancer biopsies from the two TNBC and five HR+ patients described in the previous section. The relative tumor purity of metastases, combined with EPCAM-based flow cytometry sorting produced single cells that were virtually all tumor related, as shown by the inferCNV analysis (**Suppl. Fig. S2**). As a result, we used metaVIPER to integrate results from both a bulk-level ARACNe network— generated from the TCGA breast cancer cohort—as well as a network generated from the scRNA-seq profiles captured in this study (see STAR methods). This approach allows optimal dissection of tumor cell-specific interactions (from single-cell profiles), while still providing adequate coverage (from bulk profiles) of the transcriptional targets of regulatory proteins that are undetectable in single-cell profiles.

MetaVIPER computes the normalized enrichment score (NES) of a protein’s targets in genes differentially expressed between each individual cell and a reference state, typically the centroid of the entire single-cell population (see STAR methods). As a result, positive and negative NES scores indicate higher and lower protein activity compared to the average of the single-cell population, respectively. While VIPER is most effective in assessing the activity of regulatory proteins, we have shown that it can quantitate the differential activity of signaling proteins^14,39^ and surface markers^37^ with similar accuracy. As a result, we included 339 cell surface markers and 3,407 signaling proteins in the analysis (see STAR methods for selection criteria).

Due to the large-scale CNA differences detected by the analysis, inter-patient heterogeneity was highly dominant at the gene expression level, with almost each patient contributing to an independent cluster in a Principal Component Analysis (PCA) representation, using the 5,000 genes with the highest standard deviation (**Fig. 2A**). In contrast, since VIPER-inferred protein activity is robust to noise and resilient to technical artifacts that are inconsistent with the underlying regulatory network^13^ (**Fig. 2B**), protein activity-based PCA analysis virtually eliminated inter-patient variability, except when biologically relevant (**Fig. 2C**). For instance, differences linked to HR status were captured by the second principal PCA component (y-axis), which accounts for 15% of cross-cell variability. Yet, the most significant source of variance, accounting for 31% of cross-cell variability, was captured by the first PCA component (x-axis), which could be associated with high vs. low stemness (**Fig. 2D**).

**Figure 2.**
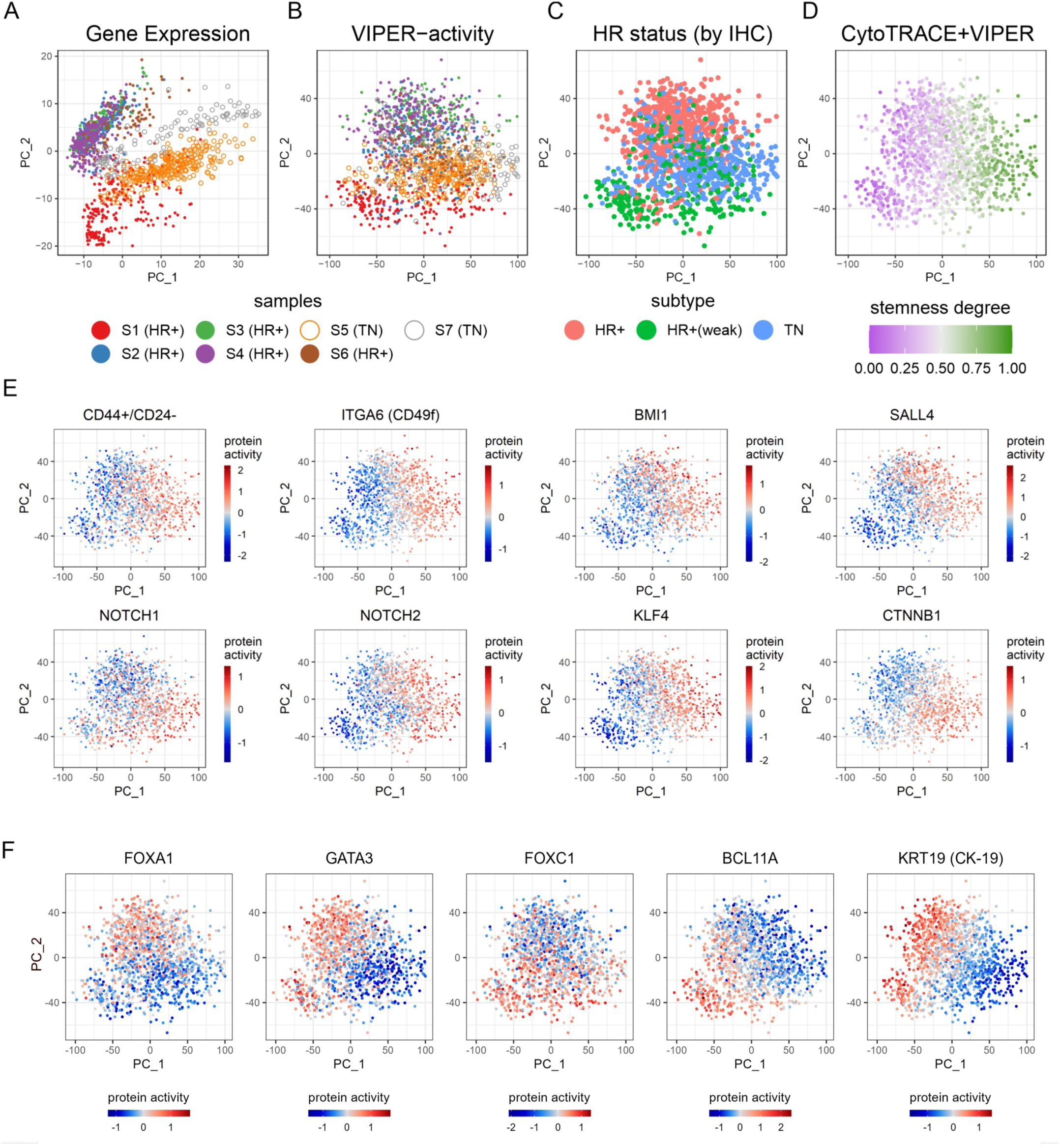
Analysis of scRNA-seq data for 7 breast cancer patient samples. Cells were clustered based on the first two principal components of the cell’s gene expression (**A**) and the protein activity inferred by VIPER (**B**-**F**). In **A** and **B,** cells are colored according to the patient they came from. **C.** The breast cancer subclasses (HR+, weakly HR+, and TN) are shown. **D**. The degree of each cell’s stemness is indicated using a green-grey-purple color gradient, corresponding to the degree of stemness from one (most stem- like) to zero (most differentiated). The stemness degree was estimated based on the combination of the CytoTRACE score and the protein activities of well-known stemness markers: CD44+/CD24-, ITGA6, BMI1, SALL4, NOTCH1, NOTCH2, KLF4, CTNNB1, ITGB3, ITGB1, PROM1, POU5F1, SOX2, and KIT (see Methods). This stemness degree score was re-scaled to the range between 0 and 1. **E**. The VIPER- inferred protein activity (centered) of individual breast CLSC markers. From the highest to the lowest, activity is shown with a red-white-blue color gradient (white = mean). **F**. The VIPER-inferred protein activity of HR+ markers (FOXA1 and GATA3), TNBC makers (FOXC1 and BCL11A), and a differentiated-cell

Cell stemness was assessed using two complementary metrics, including (a) the global activity of established breast CSLC markers and (b) CytoTRACE^15^, an experimentally validated algorithm designed to infer stemness based on gene count signature analysis (**Fig. 2D**, see STAR methods). CytoTRACE was previously validated within a hematopoietic lineage context and is based entirely on assessing expressed gene counts (a rough measure of cell entropy) rather than specific knowledge of stem cell biology. As a result, it has shown limitations, for instance, in differentiating quiescent stem cells from cycling progenitor cells^15^. To address this issue we complemented and compared the CytoTRACE analysis with biologically-relevant insights derived from the VIPER-measured activity of 14 previously reported CSLC markers, including CD44+/CD24-^40^, ITGA6 (CD49f)^26^, BMI1^4^, SALL4^41^, NOTCH1^42^, NOTCH2^42^, KLF4^43^, CTNNB1^44^, ITGB3 (CD61)^45,46^, ITGB1^47^, PROM1 (CD133)^48^, POU5F1 (OCT4)^49^, SOX2^50^, and KIT^51^, resulting in a *consensus Stemness Score*, ranging from *SS* = 0 (most differentiated) to *SS* = 1 (most CSLC), shown as a color gradient in **Fig. 2D** (see STAR methods). Supporting the use of such consensus metric, the CytoTRACE and CSLC marker-based scores were highly correlated despite being assessed by completely independent methodologies (Spearman’s *ρ* = 0.43, *p* ≤ 2.2×10^-16^) (**Suppl. Fig. S3**).

Despite the potential noisy nature of single-cell data, the PCA plot region comprising CD49f^high^ cells was strongly associated with high activity of other established markers of stem-like function in mammary epithelial cells, such as BMI1^4^ and NOTCH1/2^42^, among several others, critically in both TNBC and HR+ derived cells (**Fig. 2E, Suppl. Fig. S4**). Consistent with the literature^48,52–54^, activity of additional stemness markers such as PROM1, POU5F1, SOX2, and KIT was also more prominent in CD49f^high^ cells from TNBC patients (**Suppl. Fig. S4**). Differential activity of metabolic CSLC markers, such as ALDH1^55^, was not detectable, likely because these enzymes are less related to transcriptional regulation.

In sharp contrast to VIPER-based analyses—and fully consistent with prior studies, see^20,37,38^ for instance—the expression of genes encoding for these markers was mostly uninformative and failed to provide insight into CSLC characterization, because of the drastic gene dropout effect associated with scRNA-seq profiles (**Suppl. Fig. S5**). For instance, despite having a clear readout at the protein activity level, CD44, ITGB3, and SOX2 generated virtually no reads, thus preventing meaningful assessment of their differential expression, while expression of most other markers could not be associated to specific regions of the PCA plots.

Differential activity of subtype-specific markers was also evident for cells isolated from HR+ vs. TNBC patients, especially within the differentiated cell compartment. For instance, the activity of luminal markers, such as GATA3^56,57^, FOXA1^58^, the estrogen (ESR1) and progesterone (PGR) receptors, was markedly higher in differentiated HR+ derived cells (**Fig. 2F, Suppl. Fig. S6A**), while the activity of TNBC markers, such as FOXC1^59,60^ and BCL11A^61^, as well as basal cytokeratin (KRT17), and vimentin (VIM)^62,63^, was higher in differentiated TNBC derived cells (**Fig. 2F**, **Suppl. Fig. S6B**). To provide an objective baseline we leveraged KRT19, an established marker of luminal differentiation, whose NUMB-mediated interaction with WNT/NOTCH pathways is well documented^64,65^ and whose differential protein activity and differential gene expression could be effectively assessed in single cells. Indeed, differential expression of *KRT19* was highly consistent with metaVIPER-measured KRT19 activity (**Suppl. Fig. S7A-C**), confirming VIPER-based identification of luminal vs. basal cells. Compared to other cancers, such as colon cancer^65^, KRT19 holds special relevance in breast cancer, where its attenuated expression is strongly associated with poor prognosis and stemness^64,65^; consistent with these findings, KRT19 activity was also significantly lower in the PCA region associated with highest stemness (**Fig. 2F**).

### VIPER-inferred CSLCs are insensitive to paclitaxel

Rather than assessing self-renewal and multipotency as characteristics of *bona fide* CSLC state—still a rather controversial topic—we focused on the more pragmatic and objective assessment of the differential sensitivity to paclitaxel by cells identified as CSLC by our analysis, which presents critical relevance to patient treatment. For this purpose, we analyzed single cells dissociated from PDX models established by transplantation of a human primary TNBC in the mammary fat pad of immunodeficient NOD/SCID/IL2Rγ^-/-^ (NSG) mice, which were treated with either vehicle control or paclitaxel for 14 days after reaching a tumor volume of 100 mm^3^ (**Fig. 3A**; see STAR methods).

**Figure 3.**
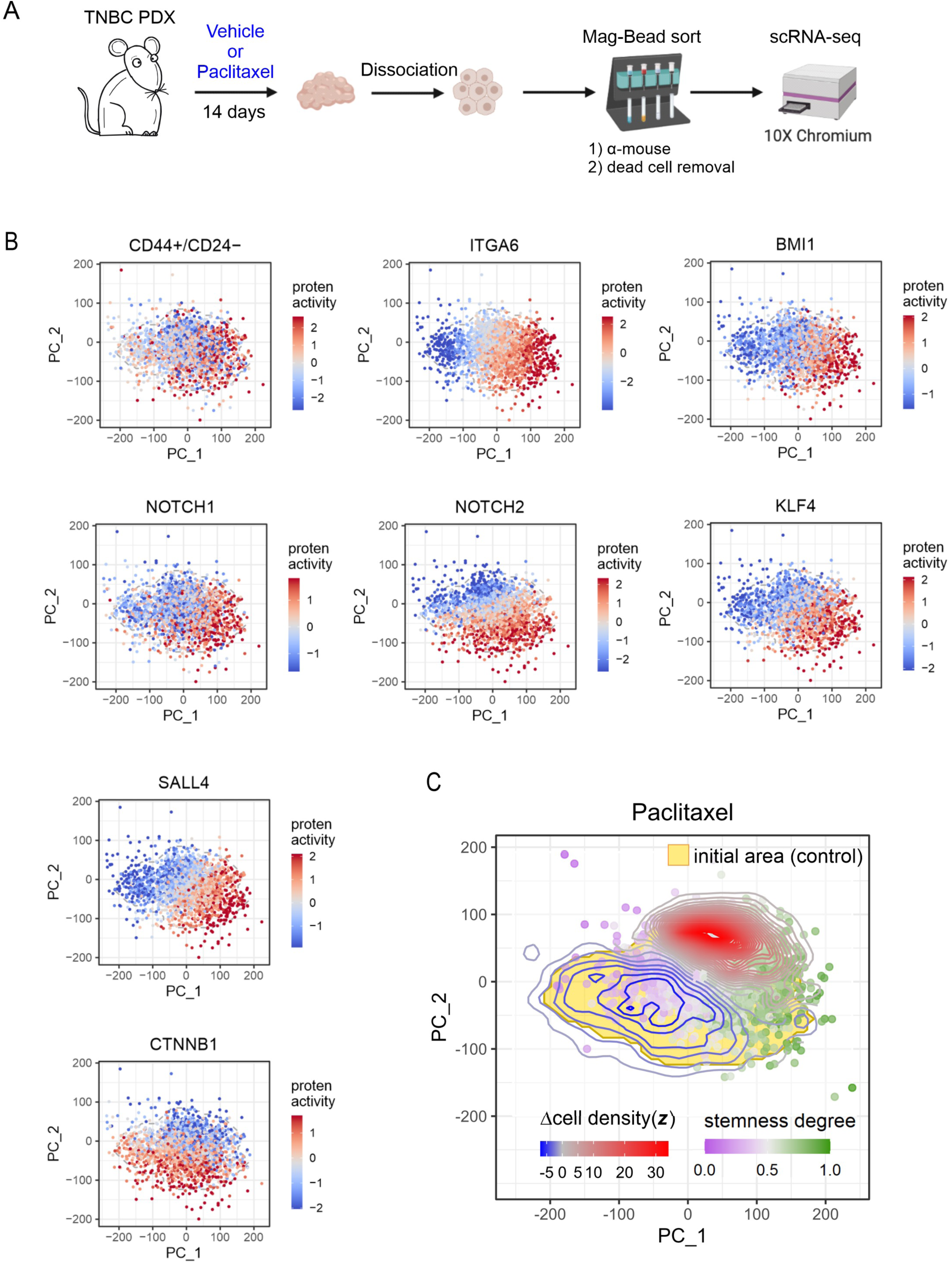
**A.** Workflow for scRNA-seq analysis of a TNBC PDX model. **B.** The VIPER-inferred activities of established breast CSLC markers in the vehicle control. **C**. Effect of Paclitaxel on the TNBC PDX cells. The cells were clustered based on the first two principal components of VIPER-inferred protein activity profiles under vehicle- and drug-treated conditions (see Methods). Based on the degree of stemness, cells were colored in a green-grey-purple color scheme (green: more stem-like cells, purple: more differentiated cells). The stemness degree was estimated by the combination of the CytoTRACE score and the protein activities of stemness markers (CD44+/ CD24-, ITGA6, BMI1, SALL4, NOTCH1, NOTCH2, KLF4, CTNNB1, ITGB3, ITGB1, PROM1, POU5F1, SOX2, and KIT) (see Methods). The estimated stemness degree score was rescaled to the range of 0-1. The area in yellow indicates a boundary of the cell cluster which 95% of cells in the control fall into. Cell density change (z-score) is shown with contour lines in the PDX sample treated with paclitaxel. The red and blue contour lines denote an increase or decrease, respectively, in cell densities under drug treatment compared to the control.

First, we assessed the fidelity of PDX-derived, single-cell subpopulations to those dissociated from human samples. Single-cell analysis of a vehicle control-treated mouse confirmed prior findings from patient-derived samples. Specifically, based on protein activity analysis with metaVIPER, the 1^st^ principal component (PC1) was again associated with cell differentiation and significantly correlated with both CytoTRACE score (Spearman’s ρ = 0.65, *p* ≤ 2.2×10^-16^, **Suppl. Fig. S8A-B**) and with overall activity of the 14 CSLC markers (ρ = 0.90, *p ≤* 2.2×10^-16^, **Suppl. Fig. S8C**). More importantly, there was a highly significant overlap of proteins differentially active in cells with the highest vs. lowest Stemness Score in PDX vs. human samples, as evaluated by GSEA analysis (OncoMatch algorithm^18^) (NES = 7.97, *p* = 1.6×10^-15^). Finally, based on GSEA analysis of MSigDB hallmarks^66^, genes encoding for proteins associated with the 1^st^ PC were highly enriched in hallmarks associated with cell developmental processes such as epithelial-mesenchymal transition and myogenesis (*p* = 3.4×10^-4^ and *p* = 1.9×10^-3^, respectively) as well as PI3K-AKT-mTOR^67^ (*p* = 9.8×10^-4^), KRAS^68^ (*p* = 2.0×10^-3^), and P53^69^ (*p* = 2.0×10^-3^) pathways (Suppl. Table 1).

Consistent with data from primary tumor tissues, differential expression of most CSLC markers in single cells isolated from PDX tissue was not informative or undetectable (**Suppl. Fig. S9**). However, at the protein activity level, the PCA regions with the highest activity of different CSLC markers—including CD49f, BMI1, CD44^+^/CD24, and NOTCH1/2—were largely overlapping in both human and mouse samples (**Fig. 3B**). Putative CSLCs from PDX samples (i.e., with highest Stemness Score) also presented high activity and expression of the established quiescent breast CSLC marker BIRC5^70^ (Spearman’s ρ = 0.53, *p* ≤ 2.2×10^-16^, **Suppl. Fig. S10**) and lower activity and expression of E2F family proteins (ρ = -0.69, *p* ≤ 2.2×10^-16^, **Suppl. Fig. S11**), which transactivate genes for G1/S transition^71^. These differences were likely more evident in PDX samples because of faster growth kinetics, as compared to primary human tumors. These data suggest that CSLC are more quiescent than differentiated cells, thus providing additional rationale for their paclitaxel resistance. Taken together, these data characterize the PDX as a high-fidelity model to study CSLC vs. differentiated cells^18^.

Changes in CSLC vs. differentiated cell density following drug treatment were then assessed by computing the normalized ratio between the number of cells with the highest (SS ≥ 0.8, most CSLC) and lowest (SS ≤ 0.2, most differentiated) Stemness Score in paclitaxel vs. vehicle control-treated samples, see STAR methods. Paclitaxel treatment induced striking depletion of differentiated cells vs. CSLCs (**Fig. 3C**) (*p* = 2.6×10^-4^, by Fisher’s exact test), thus confirming the expected paclitaxel resistance of CSLC compartment cells identified by VIPER analysis.

Since the PDX was derived from a TNBC tumor, the 2^nd^ PC could not be associated with HR status, as shown instead across the original 7 patient-derived samples. Rather, GSEA analysis revealed enrichment in two key categories, including cellular responses to DNA damage and oxidative stress, two hallmarks of paclitaxel mechanism of action (*p* = 1.9×10^-11^ and *p* =2.4×10^-12^, respectively, by GSEA) (**Suppl. Table 1**)^72–74^. Indeed, the cells that were least affected by the drug were those presenting both high stemness score and a low proliferative potential (upper right quadrant on the PCA plot). Yet, for any given value of the PC2 metagene, predicted CSLC were always less sensitive to treatment than their differentiated counterpart. Indeed, the density of cells with the highest stemness score was virtually unaffected by treatment.

### MR Analysis of human breast cancer cells

VIPER analysis has been effective in identifying candidate MR proteins representing mechanistic determinants of cell state^75,76^, as well as clinically validated biomarkers ^77–81^, see^82^ for a recent perspective. Critically, we have shown that VIPER-inferred MRs are highly enriched in tumor-essential genes^75,76,83^, such that their pharmacologic targeting can abrogate tumor viability *in vivo*^17–19^. Equally important, we have shown that genetic or pharmacologic targeting of MRs that are differentially active in molecularly distinct transcriptional states can effectively reprogram cells between these states^20,84,85^. This suggests that elucidating candidate MRs of breast CSLC state may help identify drugs that either selectively ablate paclitaxel-resistant cells or reprogram them to a paclitaxel-sensitive state, thus providing a rationale for combination therapy.

To discover the most conserved CSLC MRs across the available metastatic samples, we first leveraged metaVIPER to identify proteins whose transcriptional targets were most differentially expressed in the 20 cells with the highest vs. the 20 with the lowest Stemness Score in each individual patient, as well as in the PDX model, on an individual sample basis (see STAR methods). As discussed, the most differentially active proteins are also those expected to be most likely to mechanistically regulate the cell state of interest, via their transcriptional targets. As previously shown^18,19^, the PDX model was included in the analysis to help prioritize MR-inverter drugs that are conserved in a model that may be leveraged for drug validation *in vivo*.

As discussed, CytoTRACE was originally developed and validated only in a hematopoietic linage context^15^. As a result, for MR elucidation purposes, we decided to rely only on the differential activity of the 14 CSLC markers, including CD44+/CD24-, ITGA6, BMI1, SALL4, NOTCH1, NOTCH2, KLF4, CTNNB1, ITGB3, ITGB1, PROM1, POU5F1, SOX2, and KIT (see STAR methods). Indeed, while the enrichment of breast CSLC and stem-related markers in differentially active protein was still significant when CSLC were predicted by CytoTRACE analysis (NES = 2.57, *p* = 10^-2^), statistical significance increased substantially when relying only on the established CSLC markers (NES = 4.66, *p* = 3.2×10^-6^). Nevertheless, confirming that this choice has only minimal effects on MR analysis, statistically significantly MR proteins (*p* ≤ 10^-3^, Bonferroni corrected) were highly overlapping when CytoTRACE was included or excluded from the analysis (*p* ≤ 1.2×10^-44^, by hypergeometric test).

Surprisingly, independent analysis of each patient and of the PDX model produced highly consistent MR predictions, including across HR+ and TNBC samples (**Fig. 4A, Fig. S12A-B**), suggesting that CSLC MR proteins are conserved independent of tumor HR status. This provided the rationale for the generation of a consensus CSLC MR signature, obtained by ranking all proteins by integrating their metaVIPER NES across all samples, using the weighted Stouffer’s method (**Fig. 4B**, see STAR methods). Based on this analysis, in addition to the original 14 CSLC markers, other proteins broadly associated with stem cell processes—including ALDH family^86,87^, ABC family^87^, quiescent stem-cell markers (FGD5^88^ and HOXB5^89^), embryonic diapause^90^ and asymmetric cell division processes^91^ (**Suppl. Table 2**)—also emerged as significantly enriched among the most differentially active proteins (*p* = 2.0×10^-12^) (**Suppl. Fig. S13**).

**Figure 4.**
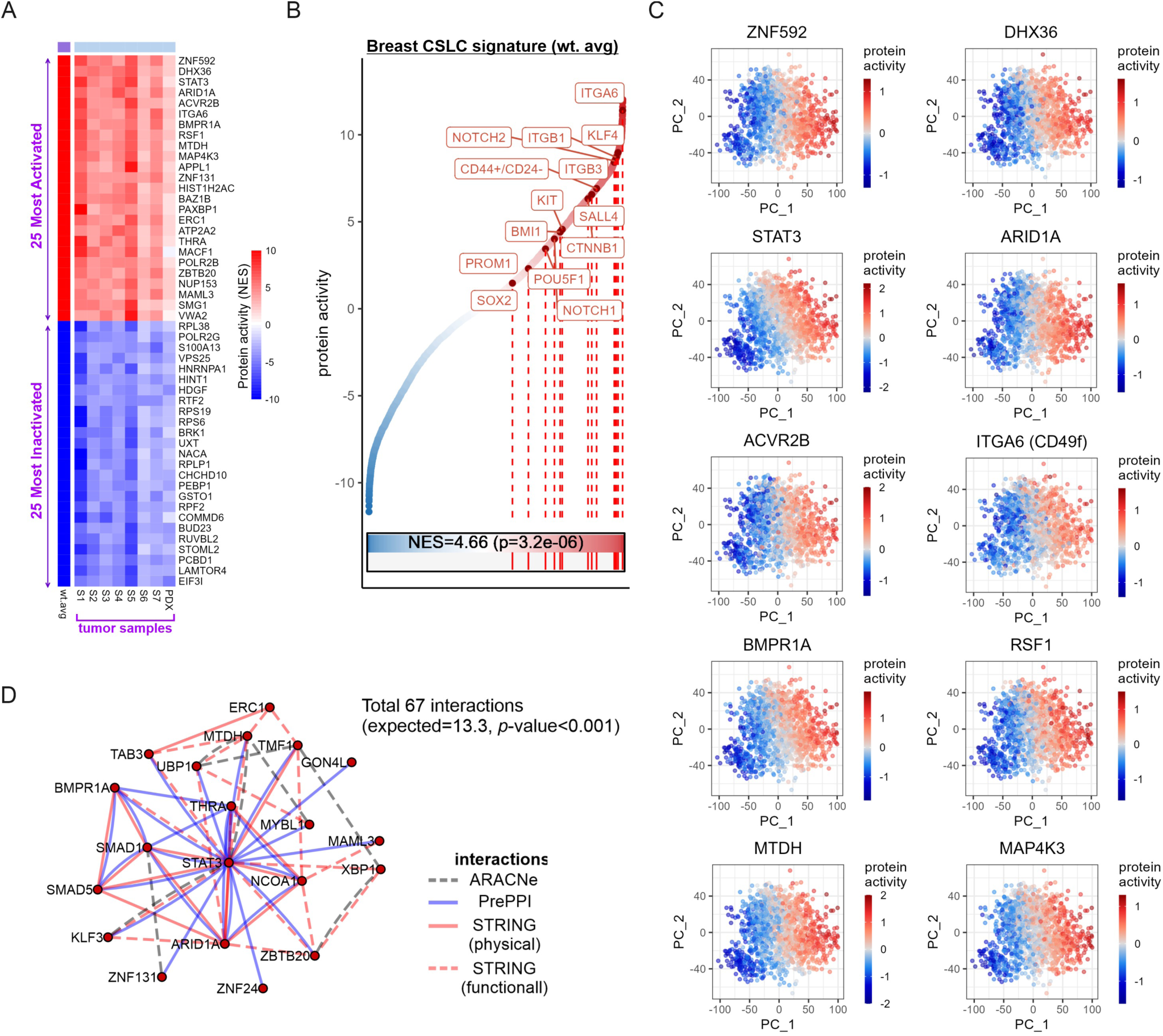
**A.** A heatmap showing the VIPER-inferred protein activity of the 25 most activated and the 25 most inactivated proteins in the breast CSLC signature and their activities in individual samples (7 patient samples and the PDX vehicle-treated sample). For each sample, differential protein activity from non-CSLCs to CSLC was computed using metaVIPER. The overall CSLC signature was obtained by the weighted average of the protein activities across samples. A larger positive (or negative) value in the signature means that the protein was more (or less) activated in CSLCs than in non-CSLCs. If there is little change in protein activity between non-CSLCs and CSLCs, the value approaches zero. Note that CSLCs and non-CSLCs were identified based on the average activity of the following CSLC markers in the sample: CD44+/CD24-, ITGA6, BMI1, SALL4, NOTCH1, NOTCH2, KLF4, CTNNB1, ITGB3, ITGB1, PROM1, POU5F1, SOX2, and KIT. **B.** A waterfall plot displaying the sorted protein activities in the breast CSLC signature, in which the signatures of individual samples were integrated using weighted Stouffer’s method. In this plot, the NES of the 14 breast CSLC markers is shown. **C.** Top 10 activated proteins in the identified signature and their protein activities in the patient data. **D.** Top 20 transcriptional regulators in the identified breast CSLC signature and their interactions identified by ARACNe, PrePPI, and STRING tools.

These results suggest that several of the most statistically significant differentially active proteins, not previously associated with breast CSLCs, may represent novel, *bona fide* MRs and potential biomarkers (**Fig. 4C and Fig. S14-18**, see also **Suppl. Table 3**), as later confirmed by CRISPR/Cas9-mediated KO (see next section). Among cell membrane-presented proteins, which may be leveraged for CSLC enrichment purposes, the analysis identified Integrin beta-8 (ITGB8) as the second most differentially active protein (after CD49f). ITGB8 was previously suggested as a marker of glioblastoma CSLCs^92^ and was identified as a prime receptor binding a latent complex of transforming growth factor beta 1 and beta 3 (TGF-β1/β3) in the extracellular matrix, responsible for activating TGF-β-associated signaling. Despite its role in tumor suppression in the early stages of tumorigenesis, TGF-β has been shown to prompt stem-like properties in advanced cancers and to increase chemotherapy resistance by promoting DNA damage response pathway activation^93–95^.

### MR Modularity Analysis

A key question in network-based analyses is whether—similar to what has been shown in other contexts^75,76,96^—candidate MRs may comprise hyper-connected, autoregulated modules providing coordinated, homeostatic cell state regulation. For this purpose, we assessed whether metaVIPER-inferred CSLC MRs were statistically significantly enriched in protein-protein and transcriptional interactions—as reported in PrePPI^97^, STRING^98^, and ARACNe-based networks—compared to an equivalent number of same-class proteins selected at random. The analysis revealed that the top 20 CSLC MRs form a highly hyperconnected module, with 67 MR-MR interactions, compared to only 13.2 detected on average in an equal size set of randomly selected proteins (*p* = 6.6×10^-7^). This supports the potential role of this module as a homeostatic On/Off switch controlling CSLC state (**Fig. 4D**), further suggesting that its inactivation may induce transition toward a more differentiated, paclitaxel-sensitive state.

### CSLC MR validation by pooled, CRISPR-KO-mediated CROP-seq analysis

To validate the CSLC MRs inferred by these analyses, CRISPR droplet sequencing (CROP-seq) was used to assess whether KO of the 25 most significant MR of CSLC state (MRCSLC, i.e., most active proteins in CSLC vs. differentiated cells) and 25 most significant MRs of differentiated state (MRDIFF, i.e., most active proteins in differentiated cells vs. CSLCs) would induce reprogramming towards a more or less differentiated cell state, respectively. To optimally assess reprogramming, we selected two breast cancer cell lines that most effectively recapitulate the CSLC state, also assuming that all cell lines comprise differentiated cells. For this purpose, we assessed the enrichment of proteins in the consensus CSLC MR signature in proteins differentially active in each CCLE breast cancer cell line (based on bulk RNA-seq analysis), and ranked them from the one with the highest NES (HCC1143)—*i.e.*, most likely to be enriched in CSLCs—to the one with the most negative NES (VP229)—*i.e.*, most likely to be enriched in differentiated cells—(**Suppl. Fig. S19**). We then selected two of the most CSLC-enriched cell lines for CROP-seq assays, including HCC1143 (ranked No. 1) and HCC38 (ranked No. 3), which were also supported by literature evidence on CSLC content^99,100^. Single-cell analyses confirmed that both cell lines had substantial CSLC representation, compared to two of the most differentiated cell lines (MCF7 and HCC2157), with HCC1143 presenting a greater fraction of differentiated cells compared to HCC38, potentially due to spontaneous differentiation in culture conditions (**Suppl. Fig. S20**).

The primary objective of CRISPR-Cas9-mediated gene knockout (CRISPR-KO) is to abrogate the function of the target protein. While it may reduce transcript copy number through mechanisms like nonsense-mediated decay, this effect is inconsistent and not generally detectable^101^. Therefore, we assessed KO efficiency based on VIPER-mediated analysis of the target protein in cells harboring the associated targeting guide RNAs (sgRNA) vs. intergenic control sgRNAs (see STAR methods). For each MR, we used 3 distinct sgRNAs and disregarded the effect of sgRNAs detected in < 10 cells. This allowed computing the effect of CRISPR/Cas9-mediated MR-KO on cell state, using the scRNA-seq profile of 10 or more cells containing the same targeting sgRNA, compared to cells harboring intergenic control sgRNAs. We then plotted the resulting effect on cell state reprogramming in HCC38 and HCC1143 cells by integrating across all positive and negative MRs of CSLC state (**Fig. 5A)**, as well as on an MR-by-MR basis (**Fig. 5B**). The expectation is that KO of positive and negative MRs will induce reprogramming towards a more or less differentiated state, respectively, as assessed by Stemness Score analysis. To avoid biasing the analysis, the MR directly targeted by a sgRNA in each cell was excluded from the Stemness Score assessment, such that only its downstream effectors were considered (see STAR methods).

**Figure 5.**
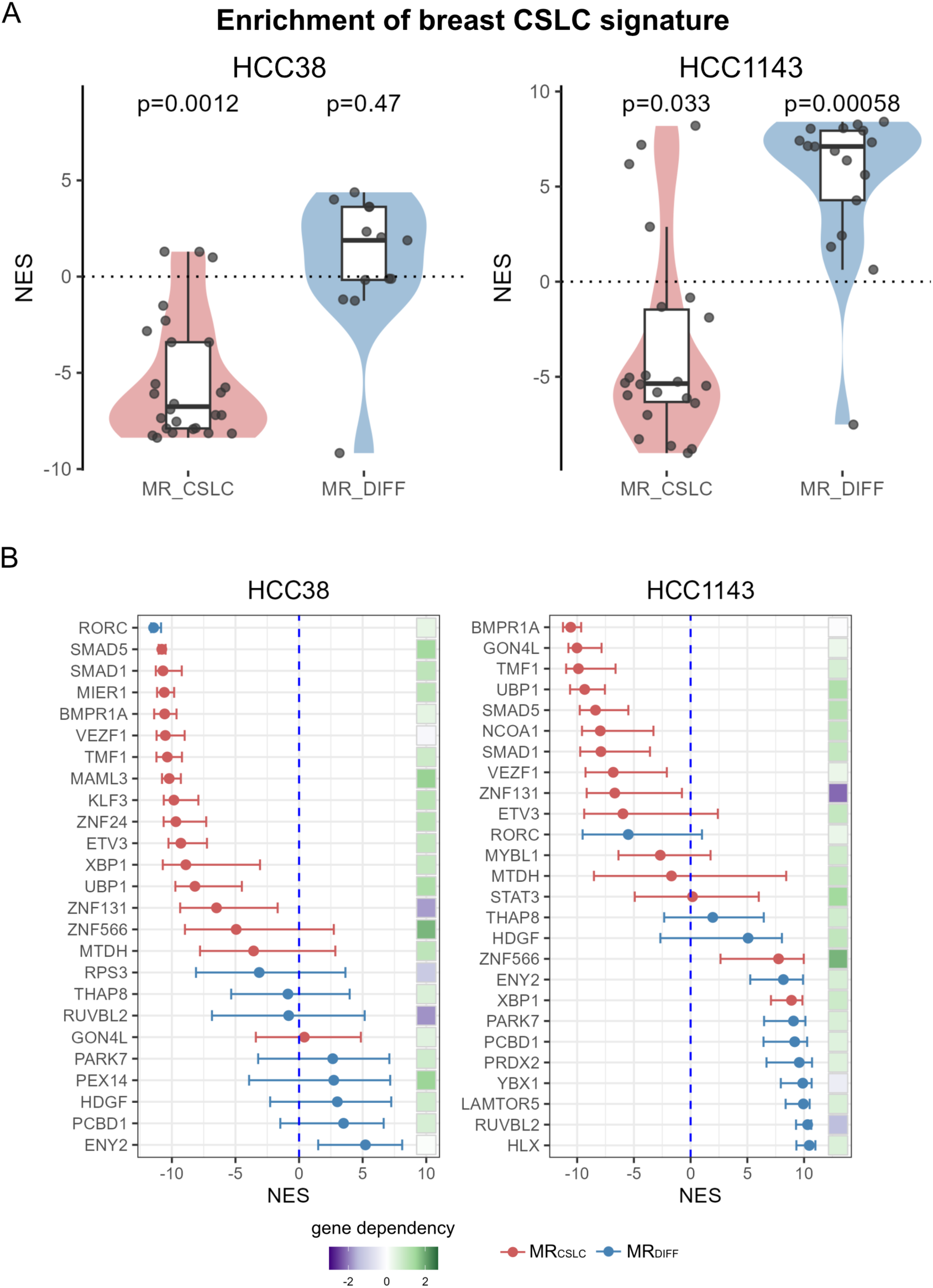
**A.** Cellular reprogramming after knocking out the top 25 most activated MRs (MR_CSLC_) and the 25 most inactivated MRs (MR_DIFF_) of breast CSLC signature, compared to the effects in the control sgRNA group for the breast cancer cell lines HCC38 and HCC1143. For each sgRNA, the VIPER-inferred protein activity profiles were generated from the pseudo-bulk expression of cells detected with the same sgRNA. The knockout (KO) efficiency was determined based on the threshold of one standard deviation below the target gene’s mean protein activity. The enrichment score of the 50-MR set (MR_CSLC_ and MR_DIFF_) was investigated in the protein activity profiles for each group of MR_CSLC_ and MR_DIFF_ (**A**) and for each sgRNA (**B**) to assess the effects of cellular reprogramming for both HCC38 and HCC1143 cell lines. To assess the reprogramming effects of each sgRNA, pseudo-bulk expressions were bootstrapped by resampling cells with the same sgRNA with replacement. Lower NES signifies greater differentiation. Error bars indicate the 1st and 3rd quartiles of NES for the reprogramming effects of multiple sgRNAs targeting the same MR in B. The effect of each MR on cell proliferation in the CROP-seq experiment is indicated by the gene dependency score, using a green-white-purple gradient where darker purple = greater dependence. White/green indicates no significant reduction in the proliferation rate when that MR is knocked out.

Based on Stemness Score analysis and fully consistent with predictions, MRCSLC KO induced significant shift of HCC38 cell state towards a differentiated state (*p* = 1.2×10^-3^, by Mann Whitney U Test). Given the small fraction of differentiated cells in this cell line (**Suppl. Fig. S20**), however, MRDIFF KO did not induce significant shift towards a CSLC state. In contrast, both MRDIFF KO and MRCSLC KO induced significant reprogramming towards a CSLC (*p* = 5.8×10^-4^) and differentiated state (*p* = 3.3×10^-2^), respectively, in HCC1143 cells, which comprise a more balanced ratio of CSLC and differentiated cells (**Suppl. Fig. S20**). When enrichment in genes associated with stem cell process-related genes (i.e., not breast cancer-specific) was considered (see **Suppl. Table 3**) the same statistically significant trends were observed (**Suppl. Fig. S21**).

In summary, CROP-seq analysis produced highly consistent results in both cell lines, confirming the predicted role of most VIPER-inferred MRs. Note that the statistical significance of this analysis is quite underestimated, because both cell lines include a mixture of CSLC (low MRDIFF and high MRCSLC) and differentiated cells (high MRDIFF and low MRCSLC), while MR KO-mediated effects can only be assessed in cells with high MR activity. As a result, the number of validated MRs is also likely to be underestimated.

The library-normalized differential abundance of sgRNA guides targeting positive MRs was not statistically significant compared to control sgRNAs (**Suppl. Fig. S22**), confirming that these MRs have no effect on cell viability or proliferation. In contrast, differential abundance of sgRNAs targeting negative MRs was significantly lower (**Suppl. Fig. S22**), suggesting that the latter— which includes cell proliferation and viability regulators—may include more essential proteins.

The contribution of each individual MR to cell state reprogramming was then analyzed and is shown in **Fig. 5B**. For the 25 MRCSLC and 25 MRDIFF tested in this assay, we only considered sgRNAs inducing effective MR KO, based on the above-described criteria. As a result, only 16 of 25 candidate MRSCLC (<u>BMPR1A</u>, <u>MTDH</u>, <u>ZNF131</u>, <u>MAML3</u>, GON4L, <u>ZNF24</u>, <u>SMAD5</u>, <u>KLF3</u>, <u>UBP1</u>, <u>SMAD1</u>, <u>TMF1</u>, <u>XBP1</u>, <u>MIER1</u>, <u>VEZF1</u>, <u>ETV3</u>, <u>ZNF566</u>, underlined are statistically significant at *p* ≤ 0.05, FDR corrected) and 9 of 25 candidate MRDIFF (<u>PCBD1</u>, RUVBL2, <u>HDGF</u>, RPS3, RORC, <u>ENY2</u>, PEX14, THAP8, <u>PARK7</u>) could be evaluated in HCC38. Similarly, in HCC1143 cells, only 15 of 25 MRSCLC (STAT3, <u>BMPR1A</u>, MTDH, <u>ZNF131</u>, <u>GON4L</u>, MYBL1, <u>SMAD5</u>, <u>UBP1</u>, <u>NCOA1</u>, <u>SMAD1</u>, <u>TMF1</u>, XBP1, <u>VEZF1</u>, <u>ETV3</u>, ZNF566) and 11 of 25 MRDIFF (<u>PCBD1</u>, <u>RUVBL2</u>, HDGF, <u>PRDX2</u>, <u>YBX1</u>, RORC, <u>LAMTOR5</u>, <u>ENY2</u>, THAP8, <u>HLX</u>, <u>PARK7</u>) could be evaluated.

In summary, of 16 and 15 MRCSLC tested one or both cell lines, 15 (94%) and 10 (67%) were validated in at least one or both cell lines (*p* ≤ 0.05, FDR corrected), respectively. Similarly of 9 and 11 MRDIFF tested one or both cell lines, 4 (44%) and 8 (73%) were validated in at least one or both cell lines (*p* ≤ 0.05, FDR corrected), respectively.

CRISPR-mediated KO of the 5 most activated candidate MRSCLS proteins, by VIPER analysis, identified 2 (BMPR1A and ZNF141) capable of inducing highly significant (p ≤8.0×10^-24^ and p ≤ 3.5×10^-7^, respectively for HCC38 and p ≤2.1×10^-24^ and p ≤4.2×10^-6^, respectively for HCC1143 after FDR correction) Stemness Score decrease in both cell lines, confirming their mechanistic role in CSLC state regulation. Among these, ZNF131 was the only one previously associated with essentiality in these cell lines (gene dependence score = -1.76 for HCC38 and –2.16 for HCC1143 by CERES ^102^, a copy-number correction method for computing gene essentiality). Indeed, ZNF131 KD-mediated centrosome fragmentation and cell viability decrease were previously reported in GBM^103^. This raises an important question related to the potential role of ZNF131 as a CSLC-specific essential gene in breast cancer. Similarly, CRISPR-mediated KO of the 5 most inactive candidate MRDIFF proteins, by VIPER analysis, identified PDBD1 capable of inducing statistically significant (p ≤ 0.022 for HCC38 and p ≤ 7.4×10^-13^ for HCC1143, FDR corrected) Stemness Score increase in both cell lines. Taken together, this confirms that VIPER-inferred MRs are highly enriched in mechanistic, causal determinants of CSLC state rather than pure gene/phenotype statistical associations.

### Identification of drugs able to invert stem-like MR programs

The high validation rate of VIPER-inferred MRs in the CROP-seq analysis suggests that MR-inverter drugs capable of inhibiting and activating the most positive and negative MRs, respectively, should induce CSLC differentiation, thus increasing their sensitivity to chemotherapy. Indeed, MR-mediated reprogramming of cell state has already been validated in multiple contexts, from de- differentiation^84^, to reprogramming^96,104^ and trans-differentiation^85,105^. For this purpose, we leveraged the OncoTreat algorithm, which has proven highly effective in discovering MR-inverter drugs that were extensively validated *in vivo*, based on MR proteins inferred by VIPER analysis of both bulk^17–19^ and single-cell profiles^20,21^.

OncoTreat relies on perturbational RNA-seq profiles representing the response of cells—selected based on their ability to phenocopy the MR activity signature of interest—to treatment with multiple drugs and vehicle control. Perturbational profile analysis, using VIPER, allows measuring the differential activity of each MR in drug vs. vehicle control-treated cells thus providing a quantitative assessment of the activity inversion across the entire MR-signature. For this purpose, we used previously generated perturbational profiles in the BT20 BRCA cell line, which strongly recapitulates the consensus CSLC MR signature (6^th^ most significant among 62 BRCA cell lines in CCLE, (NES = 7.3 by enrichment analysis), **Suppl. Fig. S22A-C**). Specifically, BT20 cells were profiled at 24h following treatment with 90 clinically relevant drugs, including FDA-approved, late-stage experimental oncology drugs (i.e., in Phase II and III clinical trials) and other selected drugs^23^ (**Fig. 6A; Suppl. Tables 4,5**). Transcriptional profiles were generated using PLATE-seq^23^—a fully automated 96- and 384-well, microfluidic-based technology that is highly efficient and cost-effective—at an average depth of 2M reads. To optimize elucidation of drug mechanism of action (MoA), rather than activation of stress or death pathways, drugs were titrated at 1/10^th^ of their EC50 concentration, based on 10-point dose response curves^17^.

**Figure 6.**
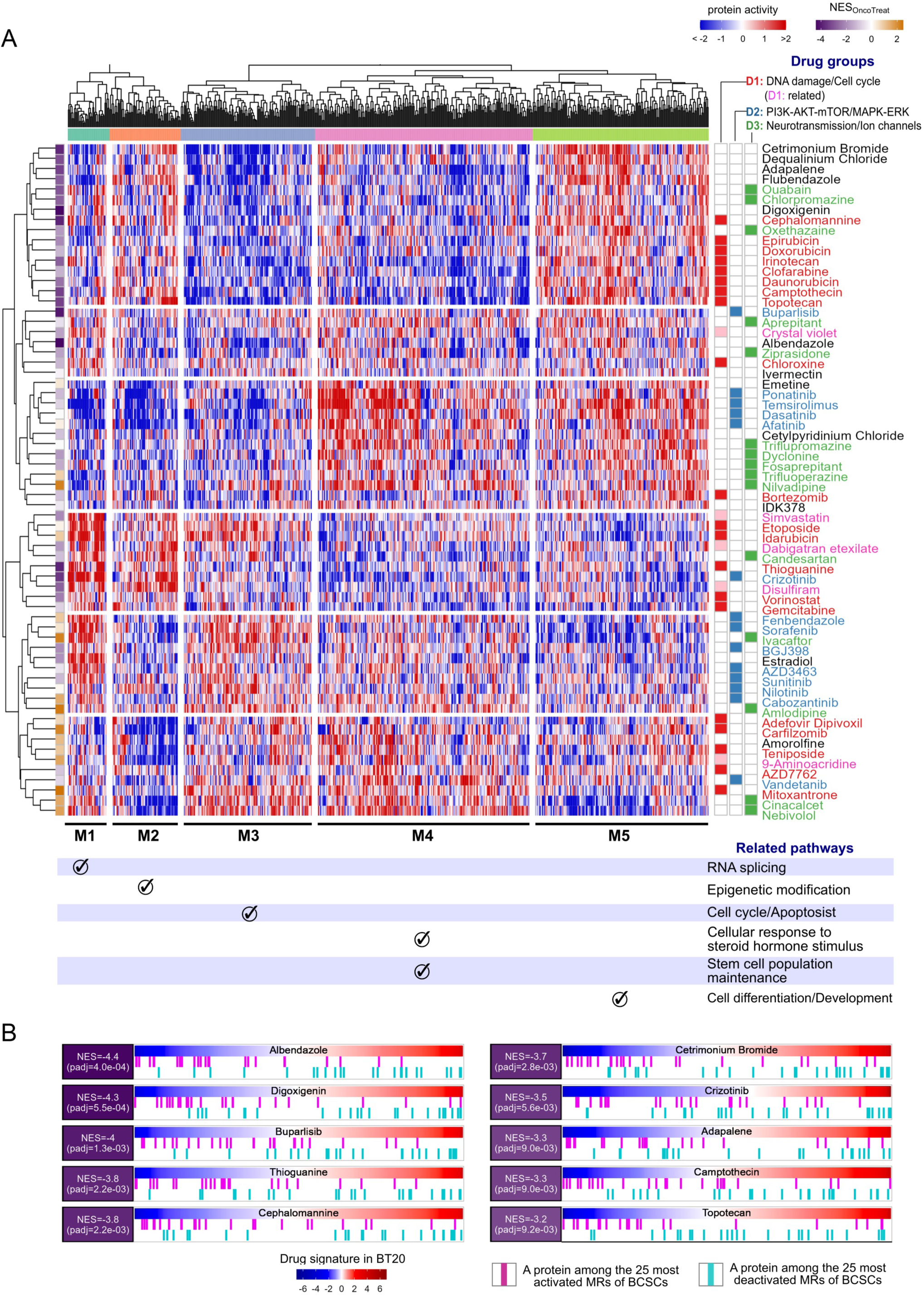
**A.** Bi-clustered drug perturbation profiles for the breast cancer cell line BT20. In the heatmap, the rows and columns are drug samples (24h, 1/10^th^ EC20) and master regulator proteins (FDR_BCSC_<1×10^-5^), respectively. The activated and inactivated proteins are shown in red and blue, and the protein activities with no change are shown in white. **B.** The enrichment plot of the top 10 drugs, predicted from the perturbation profiles with 24h treatment at 1/10^th^ the drug’s EC20 in BT20 using OncoTreat analysis. The magenta and turquoise bars denote the top 25 most activated proteins and the top 25 most inactivated proteins in the breast CSLC signature, respectively, which were derived from the 7 patient samples. In each plot, these 50

Analysis of proteins that were differentially active in drug vs. vehicle control-treated cells identified five protein clusters (M1 – M5) that were consistently activated or inactivated in response to different drug subsets. These were significantly enriched in five main Gene Ontology (GO) pathways, including RNA splicing/Ribosome biogenesis (M1), Epigenetic modification/DNA methylation (M2), Cell cycle/Apoptosis (M3), Cellular response to steroid hormone stimulus and Stem cell population maintenance (M4), and Cell differentiation/Development (M5), respectively (**Fig. 6A; Suppl. Table 4**). Notably, drugs inducing activation or inversion (*i.e.*, positive or negative NES) of breast CSLC MRs had opposite effects on the M4/M5 vs. M1/M2/M3 modules. Specifically, M5 proteins, which were associated with differentiation and developmental processes, were significantly activated by the drugs inducing strongest inversion of CSLC MR activity. In contrast, the drugs predicted to further activate the CSLC MR signature induced activation of M4 proteins, associated with stem cell population maintenance.

Among the 17 statistically significant MR-inverter drugs predicted by OncoTreat (p ≤ 0.05, FDR corrected), the anthelmintic drug albendazole emerged as the most significant one (*p* = 4.0×10^-4^) (**Fig. 6B; Suppl. Table 6**).

***Albendazole validation in vivo:*** To experimentally validate albendazole’s ability to deplete the CSLC compartment in breast cancer, we extended the protocol used to study paclitaxel in PDX models to assess the effect of 14-day treatment *in vivo* with albendazole vs. vehicle control treatment, at the single-cell level. For these *in vivo* studies, albendazole was used at 1/3^rd^ of its maximum tolerated dose in mice, consistent with assessment of MR-inversion potential at low concentration. Although albendazole is not an oncology drug, it has been shown to inhibit growth of some cancer cell lines and of a murine carcinoma, reportedly by inducing oxidative stress^106–108^. Consistently, albendazole clustered separately from chemotherapeutic drugs (**Fig. 6A**), and its activity was associated with activation of cell differentiation pathways (**Fig. 6A**).

Consistent with the paclitaxel analysis, depletion of CSLC vs. differentiated cell compartment was computed by measuring the ratio between the number of cells with the highest (*SS* ≥ 0.8) vs. lowest (*SS* ≤ 0.2) stemness score in albendazole vs. vehicle control-treated samples, normalized to the subpopulations size (see STAR methods). Whereby paclitaxel had induced dramatic increase in this ratio, indicating relative depletion of the differentiated tumor cell compartment (**Fig. 3C**), albendazole had the opposite effect (**Fig. 7A**), producing equally significant relative depletion of the breast CSLC compartment (*p* = 2.0×10^-4^, by Fisher’s exact test). When comparing albendazole to paclitaxel-treated tumors, relative changes in the density of the two compartments were even more statistically significant (*p* = 3.0×10^-12^, by Fisher’s exact test) (**Fig. 7B**), suggesting a highly complementary effect.

**Figure 7.**
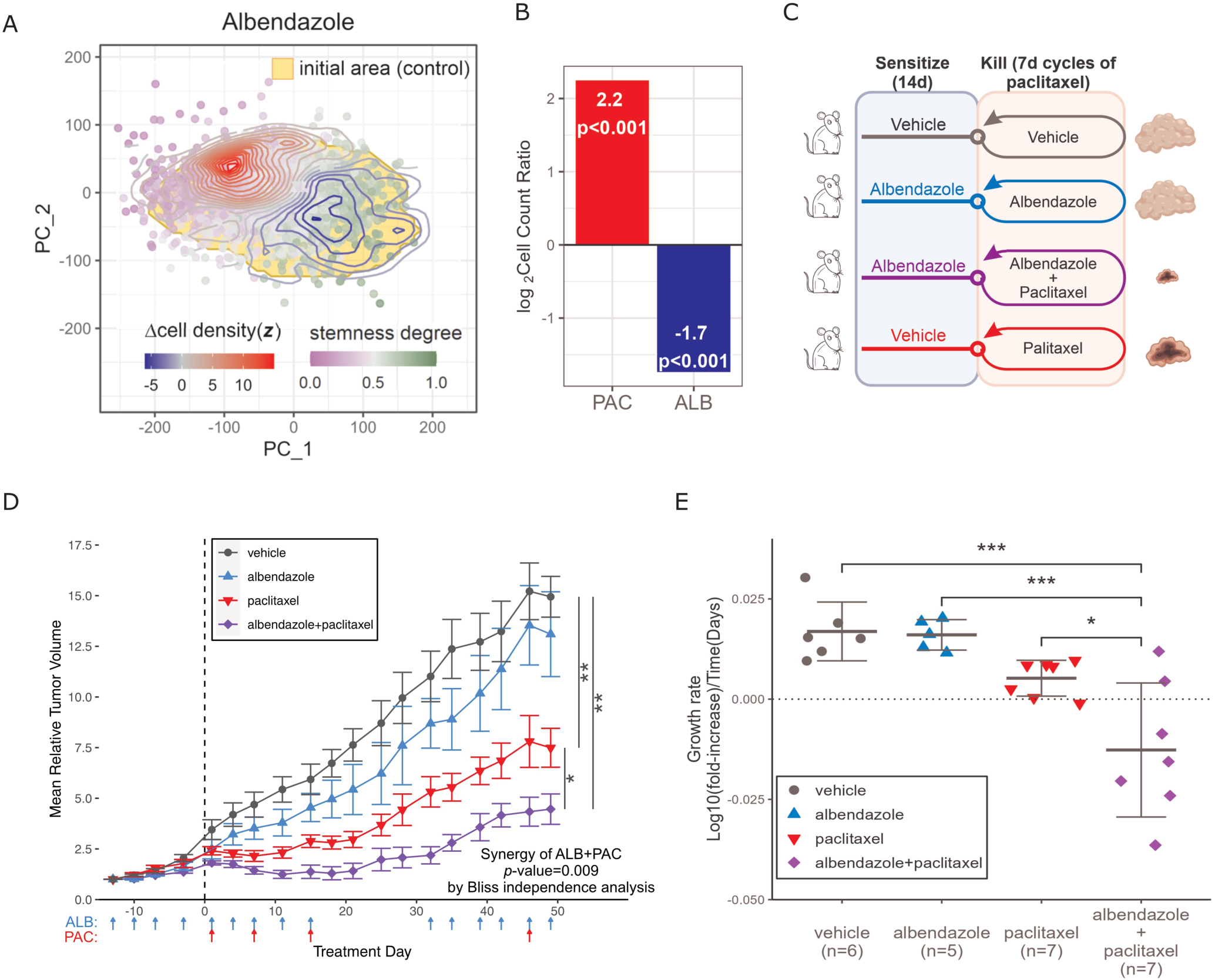
**A.** Analysis of scRNA-seq data showing the effect of albendazole on cells taken from a TNBC PDX model. The red and blue contour lines denote an increase or decrease, respectively, in cell densities under drug treatment compared to the vehicle control. The area in yellow indicates a boundary of the cell cluster into which 95% of cells in the control fall. **B.** The cell-count ratios between the stem-like (stemness score > 0.8) and differentiated (stemness score < 0.2) cells under the treatments of paclitaxel and albendazole compared to vehicle control. Based on Fisher’s exact tests, the differences between the treatments are statistically significant (*p*-value < 0.001) compared to the ratio in the control. **C.** Schematic view of combination therapies used in the preclinical tests. **D.** Mean relative tumor volumes over time under individual therapeutic strategies. Biological replicates were averaged, and the error bar indicates one standard error of the mean. Mice were treated with albendazole 3 times weekly for two weeks (Day -13 to Day 0) before the start date of the combined drug therapy with paclitaxel to sensitize the tumor cells. Mice with albendazole monotherapy were treated for the same amount of time as those in the combination therapy (Day 0 to Day 49). **E.** A comparison of the tumor growth rates during the 1^st^ cycle of drug treatments. Differences between mean growth rates were tested for statistical significance using Tukey’s honest significance test (*p<0.05, **p<0.01, ***p<0.001). Tumor growth rates were calculated assuming exponential kinetics from 1 to 18 days. The error bar indicates one standard deviation from the mean growth rates.

### Albendazole synergizes with paclitaxel in a TNBC PDX model

Since albendazole and paclitaxel deplete complementary metastatic breast cancer cell compartments, it is reasonable to hypothesize that combining or alternating their administration may outperform either drug used as monotherapy. To test this hypothesis, we evaluated whether CSLC compartment depletion by repeated administration of albendazole would enhance the *in vivo* anti-tumor activity of paclitaxel.

A PDX line, established from a human primary TNBC, was implanted in the mammary fat pad of NSG mice. When tumors reached a volume of 100 mm^3^, they were randomly enrolled to receive different treatments (paclitaxel monotherapy, albendazole monotherapy, albendazole + paclitaxel, and vehicle control) until six mice per arm were enrolled. Mice in the combination arms underwent two treatment cycles, separated by a 15-day drug holiday. Each cycle included albendazole-based sensitization for two weeks, starting at Day -13—defined as the day when a specific tumor reached a volume of 100 mm^3^—followed by three paclitaxel treatments (Day 1, 8 and 15) (**Fig. 7C**). For monotherapy treatment, mice were treated for the same amount of time and on the same schedule with albendazole, paclitaxel, and vehicle control, independently.

Paclitaxel monotherapy significantly reduced relative tumor volume (TV), compared to vehicle control (*p* = 0.0024), while albendazole was indistinguishable from vehicle control (*p* = 0.21) (**Fig. 7D; Suppl. Fig. S23**). TV change was assessed from initiation of albendazole therapy (Day -13) through Day 49; during this period, the majority of vehicle control-treated animals (n = 5 of 6) required euthanasia, due to attaining the maximal allowed humane TV endpoint (median TV = 1543 mm^3^). Additionally, compared to vehicle control, albendazole monotherapy showed no significant improvement in disease control (*p* = 0.83) or overall survival (*p* = 0.63) (**Suppl. Fig. S24**).

In sharp contrast, the albendazole + paclitaxel combination was associated with profound suppression of tumor growth, compared to both vehicle control (*p* = 1.7×10^-4^) and paclitaxel monotherapy (*p* = 0.015) (**Fig. 7E**). Drug synergy was further confirmed by Bliss independence analysis (*p* = 9.0×10^-3^) and translated into a statistically significant increase in overall survival (*p* = 0.02) (**Suppl. Fig. S24**).

## Discussion

Despite remarkable therapeutic advances, the prognosis for metastatic breast cancer patients remains dismal. Among the most critical obstacles to achieving a permanent eradication of the disease is the heterogeneity of tumor cell response to therapy. Indeed, while many chemotherapies and targeted therapies may be highly effective on subpopulations that contribute to the bulk of the malignant tissue, the presence of drug-resistant subpopulations within the same tumor mass inevitably leads to relapse and poor outcome. The cellular heterogeneity associated with pre-existing differential drug sensitivity can be of a genetic origin, for instance due to mutations in the active site of the target protein^109^ or to the presence of clonally distinct subpopulations with bypass or alternative mutations^110^. However, it is more often associated with the presence of epigenetically distinct transcriptional states with differential drug sensitivity— either pre-existing^1^ or induced by cell adaptation^111,112^—some of which can plastically regenerate the full heterogeneity of the tumor^113^. This is especially relevant in the metastatic context, where tumors have already reached a high degree of heterogeneity, due to paracrine interaction differences at distinct distal sites. Consistently, progression to metastatic breast cancer dramatically reduces the probability of achieving complete and durable responses. Indeed, most metastatic breast cancer patients rapidly progress through multiple lines of anti-tumor treatment, and eventually end up receiving conventional chemotherapy, which typically provides only short-term control of the disease.

A growing body of evidence suggests that less differentiated breast cancer cells may be chemotherapy resistant, while retaining the ability to further differentiate and reconstitute the full heterogeneity of the tumor. These cells may thus play a key role in relapse to drug-resistant disease. Tumor cells with stem-like properties (CSLCs) and tumor initiating potential were first discovered in leukemia^114,115^ and later reported also in solid tumors, such as gliomas^116,117^, breast^118^, and colon cancer^119^. As a result, the identification of novel therapeutic approaches to specifically target the CSLC compartment represents a potentially impactful area of investigation^120–122^ and may help identify drugs that synergize with chemotherapy. Network-based, single-cell analysis of cells dissociated from metastatic breast cancer patients identified a well-defined transcriptional state controlled by an exceedingly conserved repertoire of MR proteins— including transcription factors and co-factors previously associated with mammary repopulation units and breast cancer stem cells—whose sensitivity to chemotherapy is dramatically reduced compared to differentiated breast cancer cells. Indeed—based on a consensus Stemness Score that combines both the CytoTRACE metric and the activity of 14 established BRCA stemness marker proteins—there was highly significant association between cell stemness and chemotherapy resistance. This helped us identify a molecularly distinct subpopulation of chemotherapy resistant, poorly differentiated cells (CSLC for simplicity), based on the highly conserved repertoire of MR proteins that control their transcriptional state, across virtually all patients in the study. While this definition may encompass previously reported breast cancer stem cells, we use the term CSLC more broadly as it may also include an additional repertoire of undifferentiated, chemotherapy resistant progenitors. Thus, we make no claims that the CSLCs identified by our analysis represent *bona fide* tumor stem cells; rather, we show that they are chemotherapy resistant and would thus benefit from complementary therapeutic options. To enrich for CSLCs, we leveraged CD49f-based flow cytometry-based sorting of single cells dissociated from patient-derived samples. While CD49f is considered a marker of basal cells and is most highly expressed in a subset of cells from TNBC samples, previous results^24–26^ and our analysis confirmed that CD49f is also differentially expressed in CSLCs from HR+ patients. Indeed, its expression gradient was significantly correlated with the activity of 14 previously reported BRCA CLSC markers across all patients in the study, independent of HR status, thus justifying its use in our study. Confirming the value and accuracy of the proposed protein activity assessment methodology, CD49f was identified as significantly differentially active by metaVIPER in cells dissociated from human samples (**Fig. 2E**), even though its encoding gene, *ITGA6*, could not be identified as differentially expressed (**Suppl. Fig. S5**). This is fully consistent with the fact that these cells were FACS sorted with and without the associated antibody and highlights the limitations introduced by gene dropout effects in scRNA-seq profiles.

Targeting the CSLC compartment may be accomplished by developing drugs that either preferentially kill these cells or reprogram them toward treatment-sensitive states. The latter strategy is supported by recent results in fields ranging from hematopoiesis, cancer, and diabetes^84,85,105,123^ where genetic or pharmacologic targeting of MR proteins—as identified by network-based VIPER/metaVIPER analyses—effectively reprogrammed the cell’s transcriptional state towards a different target state, thus also confirming their nature as mechanistic determinants of cellular state transitions. An additional advantage of these approaches is that metaVIPER analysis effectively removes technical artifacts (batch effects) and non-functional gene expression differences, for instance due to inter-tumor CNA heterogeneity^13,38^, thus resulting in highly reproducible identification of MR proteins across samples from different patients.

To confirm mechanistic control of the CSLC state by metaVIPER-inferred MRs we performed pooled CRISPR/Cas9-mediated KO of candidate MRs in two cell lines, followed by scRNA-seq profiling, using the CROP-seq methodology. As shown, following CRISPR/Cas9-mediated KO, the vast majority of positive and negative CSLC MRs identified by metaVIPER analysis induced statistically significant reprogramming towards either a more differentiated or a more CSLC state, respectively, thus confirming the algorithm’s predictions. This includes four of the top five candidate MRs that had been previously nominated as potential players in CSLC biology but had not been experimentally validated, including STAT3^124,125^, MTDH^126^, ARID1A^127^, BMPR1A^128^, and ZNF131^103^, the first two of which had been proposed as key (co-)regulators of breast CSLCs, through the JAK/STAT3 and NF-kB pathways, respectively^124–126^. These two pathways are not only crucial in immune and inflammatory response but also pivotal for crosstalk between tumor and immune cells, especially in tumor microenvironment^129^. Moreover, the downstream effectors of these signaling pathways are often linked to cell survival and self-renewal as well as tumor proliferation, invasion, and metastasis^130^. Of these five metaVIPER-nominated MRs, only MTDH failed to induce statistically significant reprogramming in HCC38 and HCC1143 cells.

With the possible exception of ZNF131, CRISPR/Cas9-mediated KO of positive CSLC MRs had virtually no effect on cell viability, confirming that cells were reprogrammed to a chemotherapy sensitive state and not selectively ablated. This supports the identification of the MR-inverter drugs via the OncoTreat algorithm, leading to the selection and *in vivo* experimental validation of the anthelmintic albendazole as a highly efficient mediator of CSLC reprogramming. Consistent with these findings, combination therapy with albendazole and paclitaxel resulted in more profound and durable responses, as compared to either monotherapy, leading to a statistically significant increase in overall survival of preclinical models.

Remarkably, since metaVIPER identified a CSLC transcriptional state (and associated MR signature) that was virtually identical across all the tissues and models in this study, irrespective of hormone receptor status, we anticipate that the synergy between albendazole and paclitaxel in a PDX model from a metastatic TNBC patient may also be conserved in HR+ tumors, potentially in combination with hormonal blockade therapy, and may thus be relevant to a large fraction of metastatic breast cancer patients, especially since albendazole is well tolerated.

In parasites, albendazole’s mechanism of action is mediated by high-affinity binding to beta tubulin. While the binding is quite selective for parasite tubulin, the drug retains some tubulin- disrupting activity in cancer cells, even though no cytotoxicity is observed at clinically relevant concentrations. Consistently, there are a few tubulin-binding antineoplastic drugs in clinical trials—such as PTC596—that do not present the anti-mitotic cytotoxic effects of drugs such as paclitaxel, which induce harmful myelosuppression. Indeed, no cytotoxic effects of albendazole were detected in this study, either *in vitro* or *in vivo*. While It has been hypothesized that drugs like PTC596 may work by modulating trafficking of CSLC proteins, like BMI-1, and DNA repair proteins, which may provide a partial rationale for albendazole’s effect in CSLCs, and despite its highly reproducible effects *in vitro* and *in vivo*, the precise mechanism of action by which albendazole inverts the activity of CSLC MRs remains to be elucidated and will be the subject of future research. Notably, even though the study was limited to 90 drugs, it identified 17 as statistically significant candidates to reprogram CSLCs to a paclitaxel-sensitive state. As a result, we expect that extending this highly cost-effective approach to much larger drug/compound libraries may reveal even more potent agents.

Taken together, the data presented in this manuscript show that drugs targeting heterogeneous, drug-resistant subpopulations can be effectively identified by single-cell, network-based analyses and that non-oncology drugs may be effectively repurposed to enhance the therapeutic activity of anti-tumor agents, including chemotherapy.

## Acknowledgements

The results shown in this manuscript are in whole or part based upon data generated by the TCGA Research Network: https://www.cancer.gov/tcga. This study was supported by the NCI Outstanding Investigator Award R35CA197745, the NCI Office of Cancer Target Discovery and Development (CTD2) awards U01CA217858 and U01CA272610, as well as the NIH Shared instrumentation grants S10OD032433, S10OD012351 and S10OD021764, all to AC. This publication was also supported by the National Center for Advancing Translational Sciences, National Institutes of Health, through Grant Number UL1TR001873. These studies used the resources of the Herbert Irving Comprehensive Cancer Center Flow Cytometry Shared Resources funded in part through the NCI Center Grant P30CA013696. The content is solely the responsibility of the authors and does not necessarily represent the official views of the NIH.

## Methods

### Lead contact

Further information and requests for resources, reagents, and code should be directed to and will be fulfilled by the Lead Contact, Andrea Califano.

### Materials availability

This study did not generate new unique reagents.

**Patient-derived tumor dissociation and staining:** Fresh tumor fragments were acquired from tumor resections of 5 metastatic HR+ and 2 metastatic TNBC breast cancer patients, acquired under IRB AAAB2667. Fragments were quickly dissected to remove necrotic or calcified tissue and then minced, followed by single-cell dissociation using a Miltenyi gentleMACS Octo (Miltenyi, 130-096-427). The latter was performed at 37C using a human tumor dissociation kit (Miltenyi 130-095-929), as per the manufacturer’s instructions with the following revisions. When running the gentleMACS, samples were checked after 30 minutes and removed if dissociation was completed. Otherwise, they were re-checked every 15 minutes for a maximum of 60min. At all steps, it is critical to work quickly and, whenever possible, on ice to avoid substantial transcriptional changes.

On removal from the gentleMACS, single-cell suspensions were filtered through a 100uM strainer (Miltenyi, #130-098-463). Cells were then pelleted, supernatant was removed, and red blood cells were lysed using RBC lysis buffer (Invitrogen, 50-112-9751). RBC lysis buffer was diluted in DMEM (Gibco, #11965084), and the cells were washed prior to resuspension in FACS stain buffer (BD, # 554656). Cells were stained as follows:

| epitope | color | manufacurer | part # | ul per 100ul test |
| --- | --- | --- | --- | --- |
| CD49f | APC | biolegend | 313616 | 5 |
| CD326 (Ep-CAM) | FITC | biolegend | 369814 | 5 |
| CD3 | PE | BD | 561803 | 20 |
| CD45 | PE | BD | 103105 | 20 |
| CD64 | PE | BD | 561926 | 20 |
| CD16 | PE | BD | 561725 | 20 |
| CD31 | PE | biolegend | 303105 | 5 |
| CD140A | PE | biolegend | 323505 | 5 |

Cells were stained for 20min on ice, then washed 2X in cold stain buffer. Cells were resuspended in 500-1000ul stain buffer and filtered prior flow cytometry-based sorting (FACS) for CD49f^high^, CD326^+^, DAPI^-^, Lin^-^ . The number and purity of tumor cells obtained varies widely based on fragment size, cellularity, necrosis, calcification, RBC and adipocyte contamination, and other factors.

### Plate-based scRNA-seq of patient-derived samples

At the time this study started, in 2015, there was no peer-reviewed plate-based scRNA-seq method that met our cost-per-cell and quality requirements. Consequently, we created a UMI-enabled, 3’ scRNA-seq method that is described in detail at: *dx.doi.org/10.17504/protocols.io.s4hegt6*. In short, single cells are sorted directly into wells filled with hypotonic lysis buffer which contains RNAse inhibitor. Plates can be frozen at - 80C for at least two weeks. After reverse transcription, wells are pooled into a single tube for NGS library generation. The resulting libraries were sequenced on a NextSeq 550 with approximately 0.5M-1M reads per well. Note that subsequent to this study, this method was deprecated in favor of mcSCRB-seq^2^, which is more sensitive. Compared to droplet-based studies, this methodology provided much more effective elimination of dead cells and debris from necrotic tissue, which caused significant clogging and sequencing artifacts as well as higher-depth sequencing. Critically, the results obtained from the earlier technology were highly consistent with later technologies, including the 10X Genomics Chromium platform. Indeed, the initial results from human samples, including the Master Regulator proteins associated with BRCA stem-like progenitors, were fully recapitulated in single-cell profiles generated by Chromium 10X library generation from PDX models.

### Patient-derived xenograft (PDX) model

Patient-derived xenografts were generated by implanting cells obtained from a triple-negative breast carcinoma (TNBC) patient. Tumor cell suspensions were mixed (1:1) with Matrigel (Corning) and implanted into the mammary fat pad of 6 to 8-week-old female NOD (NOD.Cg-Prkdc^scid^Il2rg^tm1Wjl^/SzJ) SCID gamma (NSG) mice (Jackson Labs, Strain #005557) for scRNA-seq and therapeutic studies.

Preclinical studies in the PDX model

Paclitaxel was formulated in sterile saline and dosed at 25 mg/kg intravenously every week. Albendazole was formulated in 10% (v/v) DMSO (dimethylsulfoxide), 0.5% (w/v) carboxymethylcellulose, and 1% (v/v) Tween 80 in sterile water and dosed at 50 mg/kg IP 3 times weekly. The vehicle control mice were given 10% (v/v) DMSO (dimethylsulfoxide), 0.5% (w/v) carboxymethylcellulose, and 1% (v/v) Tween 80 in sterile water.

For therapeutic studies, tumor measurements were made biweekly using calipers. Treatment was initiated when tumor volume (TV) reached ∼100 mm3. TV is calculated using a modified ellipsoid formula: ½ (length X width2). Mice were treated with albendazole for two weeks prior to initiating three weekly paclitaxel cycles (n=7 mice/arm). Mice were treated for up to 2 cycles of albendazole + paclitaxel therapy with a 2-week treatment break between cycles as well as with individual monotherapies and vehicle control. Mean tumor volume and tumor growth rates were compared among treatment groups, using a Tukey’s honest significance test. Tumor growth rates were calculated assuming exponential kinetics3 during the first cycle (from 1 to 18 days). Under this assumption, the growth rate (b) of each tumor model can be calculated by solving the following linear regression:

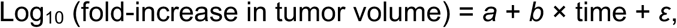

where a is the log-scaled initial tumor volume, and ε denotes the error.

To support collection of sufficient tumor material for the analysis, treatments for single cell drug response studies were initiated when TV reached ∼350-400 mm3. Animals (n=2/arm) were treated for 15 days and tumors harvested 2 hours after administration of the last dose of vehicle or drug. Harvested tissue was enzymatically dissociated into a single cell suspension for downstream scRNA-seq using the Human Tumor Dissociation Kit (Miltenyi, 130-095-929) according to manufacturer’s recommendations (other than the modifications discussed in the patient-derived sample preparation, above).

### scRNA-seq profiles from PDX samples

Tumors were resected at the Columbia University Irving Medical Center (CUIMC) and stored in cold DMEM media on ice for transport from surgery to the lab. After gross necrotic tissue removal using a scalpel, we used a gentleMACS Octo (Miltenyi, 130-096-427) to dissociate minced samples to single-cell suspensions using a human tumor dissociation kit (Miltenyi, 130-095-929). The procedure was performed at 37C, as per the manufacturer’s instructions, with the following change: samples were checked after 30min and the protocol was terminated if chunks of undissociated tissue were not visible. If a significant amount of undissociated sample remained, the program was run for 15min additional time. After dissociation, samples were filtered through a 100uM strainer, the cells pelleted, the supernatant removed, and red blood cells were lysed using RBC lysis buffer (Invitrogen, 50-112-9751). RBC lysis buffer was diluted in DMEM, and the cells were washed prior to mouse cell depletion using magnetic beads (Miltenyi, 130-104-694) following the manufacturer’s instruction. The resulting suspension of human tumor cells was adjusted in DMEM medium (Gibco, #11965084) to a concentration appropriate for loading into a 10X genomics Chromium controller (3’ gene expression kit, #120267) with an expected output of 7,000 cells/sample. Samples were run in separate wells and libraries constructed according to the manufacturer’s instructions. Libraries were sequenced on an Illumina Novaseq 6000, aiming for approximately 100,000 reads/cell. The raw PDX data were deposited in GSE226329.

### Generation of scRNA-seq profiles from cell lines

To generate single-cell profiles of breast cancer cell lines, MCF7, HCC2157, HCC38, and HCC1143 cells were seeded into 15cm circular plates and incubated for 72 hours to achieve < 50% average confluence, followed by dissociation and scRNA-seq profile generation. Specifically, the resulting human tumor cell suspensions— HCC38 and HCC1143 in batch 1 and MCF7 and HCC2157 in batch 2—were diluted to the appropriate concentration for loading into a 10X genomics Chromium controller (3’ gene expression kit, #120267) at an expected output of 5,000 cells per cell line. Within each batch, cells were pooled into the same well and libraries were constructed according to the manufacturer’s instructions. Libraries were sequenced on an Illumina Novaseq 6000, aiming for approximately 100,000 reads/cell. Pooled scRNA-seq samples—i.e., sample1, including pooled reads from HCC38 and HCC1143 cells and sample2, including pooled reads from HCC2157 and MCF7 cells—were deposited in GEO (accession number: GSE241115).

### Cell Culture Conditions and Media

HCC1143 - RPMI 10%FBS + pen/strep HCC38- RPMI 10%FBS + pen/strep MCF7 - EMEM 10%FBS + 0.01 mg/ml Insulin + pen/strep HCC2157 – DMEM 10%FBS + pen/strep All cell lines were routinely tested for mycoplasma contamination. Cell lines were kept in a 37 °C humidity-controlled incubator with 5.0% CO2.

### CROPSeq library design

For the CROPSeq screening4, we designed a target gene list that included the top 25 predicted MR proteins (MRCSLC) and bottom 25 predicted MR proteins (MRDIFF) of the CSLC vs. differentiated cell states, for a total of 50 candidate MR genes in total. Each gene was targeted with 3 sgRNAs designed using CRISPick5,6. 15 additional sgRNAs targeting intergenic genomic regions7 were selected as negative controls for the assay.

sgRNA oligo synthesis and library cloning

Oligo libraries (165 oligos) were ordered from Twist-biosciences in following format: CGATTTCTTGGCTTTATATATCTTGTGGAtttCGTCTCCCACCGNNNNNNNNNNNNNNNNNNNNGTTTAGAGACGAAAGAGCTAAGCTGGAAACAGCATAGCAAG

Twist oligo pool was amplified by using following protocol:

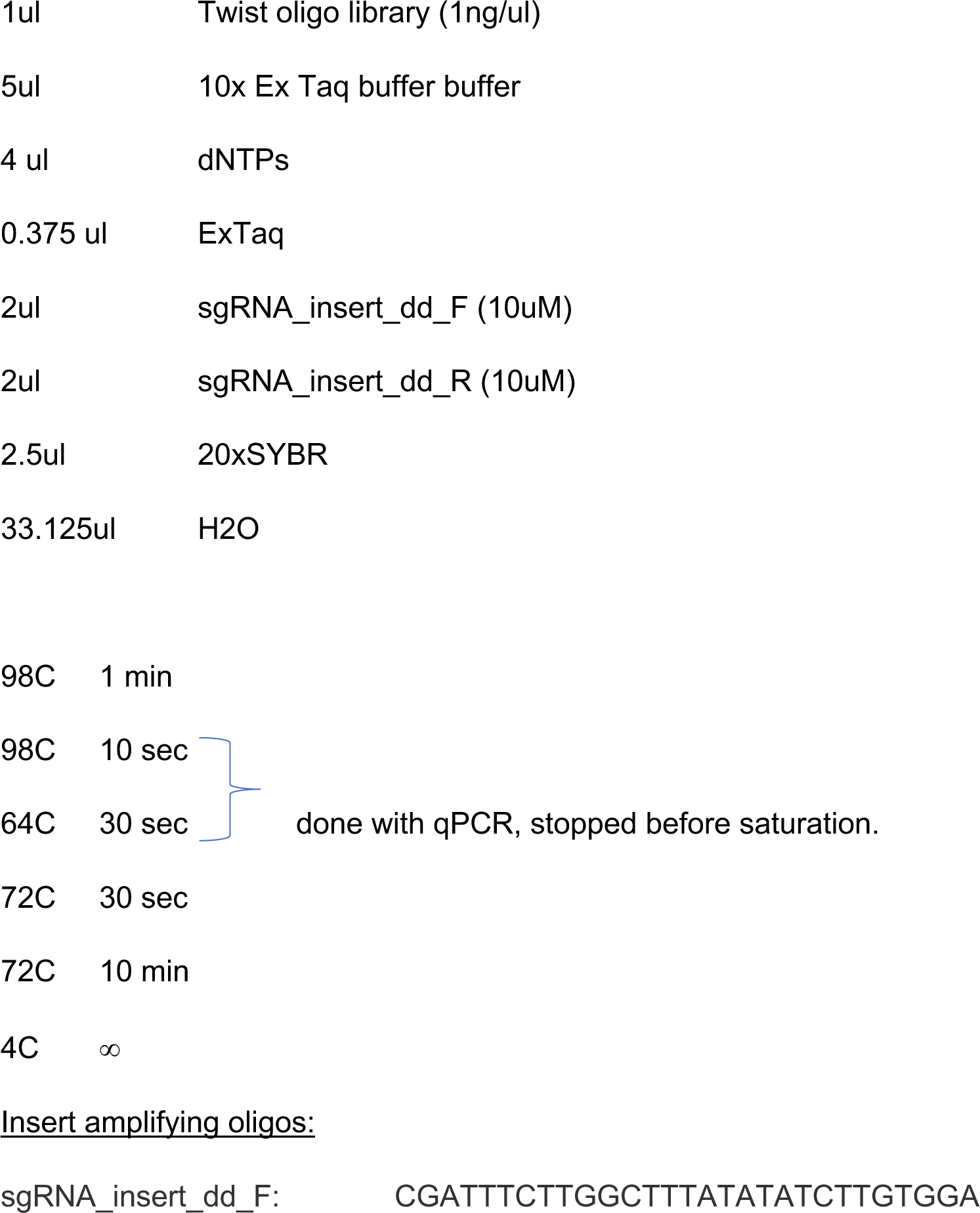
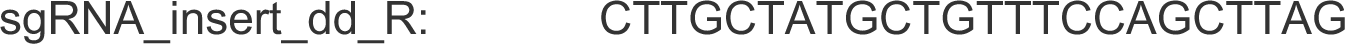

After PCR amplification, the insert was gel purified (GeneJet) and Golden-gate cloned into BsmBI-digested CROPseq-CaptureSeq-Guide-Puro-plasmid8. The Golden-gate cloned product was Isopropanol precipitated and large-scale electroporated into Lucigen Enduro competent cells. The bacterial colonies were scraped from 24,5cm x 24,5cm agar plates, so that the estimated library complexity was > 1000 colonies / sgRNA. The sgRNA library plasmid DNA was extracted by using NucleoBond Xtra Midi kit (Macherey-Nagel).

### Lentiviral library packaging

13 million 293T cells / plate were seeded in three 15cm dishes the night preceding the transfection assay. The following morning the viral transfections were conducted the following way:

- 22.1ug sgRNA-library containing CROPseq-CaptureSeq-Guide-Puro, or modified lenti-Cas9- sgHPRT1 (Addgene #196713)9. For this study the sgHPRT1 part was removed from the lenti- Cas9-sgHPRT1-vector.

- 16.6ug PsPAX2 (Addgene 12260)

- 5.5ug PMD2G (Addgene 8454).

- 1660ul of sterile H2O.

After mixing the plasmids, 110,6ul of Fugene HD (Promega) was added to the mix.

The transfection mixture was vortexed, and incubated 10 minutes in room temperature before adding dropwise to 293T cells. Altogether 3 x 15cm plates were transfected for sgRNA-library containing modified CROPseq-CaptureSeq-Guide-Puro and 1 x 15cm plates were transfected for modified lenti-Cas9-sgHPRT1.

The transfection mixture was removed the following day and virus was collected at 48h and 72h after initial transfections. To remove cellular debris, the virus containing supernatant was centrifuged 500 x g for 5min and filtered by using 0.45um PES filters (Millipore). The lentivirus was concentrated by using Lenti-X concentrator (Clontec), aliquoted and stored at -80C.

### Generation of Cas9 expressing cell lines

Cas9 expressing cell lines were generated as follows: Concentrated modified lenti-Cas9-sgHPRT1 -lentivirus was transduced into breast cancer cell lines HCC1143 and HCC38 (in presence of 8ug/ml polybrene) at an estimated MOI = 0.3. The virus was removed the following day and Blasticidin was added to the cells. Blasticidin selection was continued as long as the control cells (non-transduced) were viable.

### CROP-seq screening

sgRNA harboring lentiviruses were transduced into Cas9 expressing breast cancer cell lines HCC1143 and HCC38 (in 15cm plate-format) in presence of 8ug/ml polybrene, at an estimated MOI < 0.2. After 24h, the lentivirus containing media was removed, cells were washed with PBS, and puromycin-containing media (3ug/ml) was added to the cells for 48-96h until all control cells (not virus-infected) were dead. The cells were grown for 14 days post sgRNA transduction to allow potential reprogramming effects to take place, before the scRNA-seq. The CRISPR perturbed cell lines (HCC38, HCC1143) were adjusted to a concentration appropriate for loading into a 10X genomics (3’ gene expression kit, #120267) with an expected output of 10,000 cells/each cell line. The 10x libraries were prepared based on manufacturer’s instructions with feature barcode CRISPR screening protocol. Libraries were sequenced on an Illumina Novaseq 6000, aiming for approximately 100,000 reads/cell. The raw CROP-seq data were deposited in GSE241115.

### Bliss independence analysis

We tested for pharmacological synergy using a multiplicative null model based on Bliss Independence. Briefly, the efficacy of each drug as a monotherapy was evaluated based on the reduction in PDX tumor growth at 25 days. A conservative empirical null model was generated by fitting a distribution generated by using every possible triplet of PDX treated with each monotherapy and vehicle control. An empirical left-tailed p-value was then computed for each combination-treated PDX model using the empirical null model; these p-values were integrated across independent biological replicate PDX models using Fisher’s method to produce a single synergy p-value for each drug combination.

### Kaplan-Meier analysis

Kaplan-Meier analyses comparing disease control and overall survival across treatment groups was performed using the log-rank test. The endpoint for disease control is defined as the time to TV doubling relative to TV at the initiation of paclitaxel therapy (defined as Day 1). Overall survival is defined as time to mouse death, attainment of maximal allowed TV (humane endpoint), or development of moribund status requiring euthanasia. A Benjamini-Hochberg multiple testing correction was applied to relative TV comparisons and Kaplan-Meier analysis with adjusted p-values reported. Statistical significance was defined as p<0.05. All in vivo studies were performed in accordance with institutional guidelines and under protocol #16-08-011 approved by the Memorial Sloan Kettering Cancer Center (MSKCC) Institutional Animal Care and Use Committee.

### InferCNV

We applied inferCNV of the Trinity CTAT Project (https://github.com/broadinstitute/inferCNV) to the scRNA-seq data of patient tumor samples in order to estimate the copy number of variation of whole chromosomes and confirm the purity of tumor cells in a data set. For CNV reference data, we used the scRNA-seq data (GEO access number: GSE113196) of normal breast epithelial cells containing one basal and two liminal types in four different individuals from Nguyen et al., 201810. For inferCNV, we used the order of human genes based on transcription start sites in GRCh38.

### Protein activity analysis

Protein activity was computationally assessed using metaVIPER11. Briefly, metaVIPER is the single cell adaptation of the Virtual Inference of Protein-activity by Enriched Regulon analysis (VIPER)12. VIPER computes the normalized, rank-based enrichment score (NES) of a set of genes representing the transcriptional targets of each protein (regulon) in genes differentially expressed when comparing the cell state of interest to a reference cell state, generally the centroid of all the cells in the analysis. Thus, statistically significant positive and negative NES values identify either activated or inactivated proteins in the state of interest vs. reference state, while non-significant NES scores identify proteins that have not significantly changed activity. Unlike gene set enrichment analysis (GSEA), VIPER uses a probabilistic model to integrate the consensus between activated and inhibited targets, as well as targets with an unclear mode of regulation, and their differential expression. The use of context-specific networks, which can improve the fidelity of interactomes, is highly advantageous in the VIPER analysis with respect to the accurate inference of the protein activity. MetaVIPER, improves the analysis by allowing the use of multiple networks to define each protein’s regulon set, this is important in single cell analyses where the optimal network may be cell specific and also helps prevent overfitting to a single network.

For patient-derived tumor samples, we first removed low-quality cells where the total read count (nReads) was greater than 35,000, the number of detected genes (nGenes) was either below 1,000 or above 4,000, and the mitochondrial RNA percentage (MT%) was greater than 25%. For PDX data, we filtered cells for MT% ≤ 16%, 6,000 ≤ nReads ≤ 85,000, and nGenes ≥ 1,000.

To assess protein activity from scRNA-seq data, raw gene expression data was normalized to log2 count per million (CPM). The data were then centered at the median expression and divided by the median absolute deviation (MAD) of each gene. Note that MAD was calculated excluding the zero counts for patient tumor data due to sparsity of single cell gene expression profiles. Genes with zero counts were set to N/A and were thus ignored by the R implementation of the VIPER algorithm used in this study.

Finally, during the metaVIPER analysis, we utilized breast cancer specific networks generated by analyzing both bulk samples and single cell RNA-seq profiles using the Adaptive Partitioning version of the Algorithm for the Reconstruction of Accurate Cellular Networks13 (ARACNe-AP), as described in the next section.

### Network inference by ARACNe-AP

Gene regulatory networks for human breast cancer were created using the extensively validated reverse-engineering algorithm ARACNe-AP13. Briefly, ARACNe-AP infers protein-gene regulatory interactions by first measuring the statistical significance of the mutual information between the gene expressions of each regulator protein and each candidate target and then by removing statistically significant candidate targets that violate the Data Processing Inequality (DPI), see14 for additional details. After the analysis, each regulator protein is associated with a set of most directly regulated transcriptional targets (regulon). Since the accuracy of VIPER analyses does not further improve when regulons are larger than 40 genes12, we conservatively retained only the 50 most statistically significant targets in each regulon. This allows to remove potential bias in VIPER NES assessment, since NES measured from larger regulons would be more statistically significant.

In this study, we inferred the regulatory interactions not only for transcriptional (co-)regulators but also for proteins indirectly involved in gene transcriptional regulation such as cell surface receptors, intracellular signaling molecules, and DNA-binding molecules. This is strongly supported by recent studies confirming that ARACNe-inferred regulons for these additional protein classes are highly effective in assessing their differential activity using the VIPER algorithm15-17. The candidate regulator proteins for these analyses (n = 8,519) were thus identified based on the following Gene Ontology (GO) terms: transcription regulator activity (GO:0140110), transcription coregulator activity (GO:0003712), DNA-binding transcription activator activity (GO:0001216), regulation of gene expression (GO:0010468), signal transduction (GO:0007165) and cell surface (GO:0009986).

ARACNe-AP was then run using 200 bootstraps and a Bonferroni-corrected statistical significance threshold of p = 0.05 for protein-target interaction inference. The following datasets were used to generate complementary Breast Cancer networks: (a) RNA-seq profiles from samples in the TCGA-BRCA cohort18 and (b) scRNA-seq profiles from human breast cancer cells isolated from PDX-derived samples. We did not use the single cells from the human samples because single cell networks must be generated on a sample-by-sample basis, to avoid major artifacts associated with batch effects, and the number of cells generated by single cell profiling of the patient-derived sample was too small to generate accurate, sample-specific ARACNe-inferred networks. As a result, single-cell networks were generated using scRNA-seq profiles from malignant human cells isolated from the PDX model using the ARACNe-AP algorithm, as described in the next paragraph.

To reduce gene dropout effects, single cell network inference from PDX-derived scRNA-seq profiles was performed by first converting individual scRNA-seq profiles to MetaCell19,20 profiles. Additionally, to avoid batch effects, MetaCells were generated independently for each treatment condition (i.e., vehicle, paclitaxel and albendazole treatment). MetaCells are produced by combining the gene expression profiles of a “seed” cell, chosen at random, with those of k single cells with the closest gene expression profile (based on Spearman correlation). As discussed in the PISCES analysis pipeline21, the per-MetaCell UMI count for optimal network inference should be ≥ 10,000. Based on the average single-cell UMI count in this study, we thus used k = 9 resulting in MetaCell profiles (i.e., pseudo-bulk profiles) generated by merging the UMI counts of 10 individual cells in each MetaCell. Nearest neighbors were identified by gene expression-based Spearman’s correlation, after regressing out cell cycle contributions from the CPM-normalized and scaled data, using Seurat22. ARACNe-AP was then used to analyze the CPM-normalized expression produced by 150 MetaCells, with seed cells chosen at random from the entire population of each treatment conditions. Note that, to avoid mixing biological conditions, the nearest neighbor cells of each seed cell were selected within the same treatment condition.

For TCGA-based networks, we utilized the read count data normalized by Fragments Per Kilobase of transcript per Million mapped reads upper quartile (FPKM-UQ) in TCGA-BRCA project, which were downloaded using TCGAbiolinks23, an R package developed to retrieve datasets from the Genomic Data Commons24 (GDC). Subtype-specific networks were generated by sub-selecting samples from the following BRCA subtypes: Luminal A (ER+/PR+/HER2-), Luminal B (ER+/PR+/HER2+), HER2-enriched (ER-/PR-/HER2+), and basal-like (ER-/PR-/HER2-), as defined in the Clinical Supplement data in GDC. Finally, we applied these 4 BRCA subtype networks as well as the PDX-derived single-cell network for the metaVIPER-based assessment of protein activity from patient samples, or only the latter network for protein activity assessment in PDX samples.

### CytoTRACE Analysis

We utilized the CytoTRACE25 R package downloaded from its official website (https://cytotrace.stanford.edu/). Briefly, CytoTRACE is a computational tool to infer a cell differentiation order, using the gene count signature that is defined as the geometric mean of top 200 genes correlating with the number of expressed genes over the cells in a dataset. CytoTRACE generates a relative score between 0 (the most differentiated) and 1 (the least differentiated) for individual cells. For patient-derived samples, we performed CytoTRACE analysis on the individual cells of each sample independently to account for potential germline differences that may create batch effects. For PDX-derived samples, which had a common germline background, we performed CytoTRACE using the single cells from all samples, including those treated with vehicle control, paclitaxel, and albendazole. For both analyses, raw counts were used as an input.

### Marker-based Stemness Score

to generate a more biologically motivated, breast cancer specific assessment of cell stemness, we integrated the metaVIPER-based NES representing the differential activity of 14 previously established markers of CSLC state, including CD44+/CD24- 26, ITGA6 (CD49f)27, BMI128, SALL429, NOTCH130, NOTCH230, KLF431, CTNNB132, ITGB3 (CD61)33,34, ITGB135, PROM1 (CD133)36, POU5F1 (OCT4)37, SOX238, and KIT39. Note that the activity of the markers was inferred as described in Protein Activity Analysis above. In addition, the activity of CD44+/CD24- was calculated as the difference between the CD44 NES and the CD24 NES, such that greater NES values would indicate greater stemness. The integration of 14 maker’s activities was done by standardizing the activities for individual markers then by averaging them. Consequently, the marker-based Stemness Score (MSS) that integrates a set of the stemness markers (M) can be summarized using the following equation:

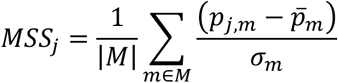

where *p*_*j,m*_ is the protein activity of marker m in cell j, *p̅*_*m*_ is the average protein activity for marker m, and *j* is a standard deviation of the protein activity of marker m.

### Integrated Stemness Score

As discussed in the main text, there was highly statistically significant correlation between the CytoTRACE (Scytotrace) and marker-based stemness (MSS) metrics, even though the methodology used to compute the two metrics were completely different and statistically independent. As a result, we proceeded to combine both metrics into a single integrated Stemness Score (ISS) by averaging the normalized score for each metric as follow.

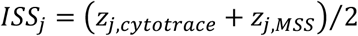

where zj,cytotrace=(Sj,cytotrace – *S̅*cytotrace)/σcytotrace and zj,MSS=(MSSj – *MSS̅*)/σMSS for cell j. Finally, ISS was linearly re-scaled to a score ranging from zero to one in Fig 2D and Fig3C, as follow.

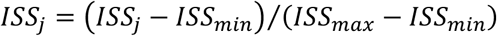

### Breast CSLC MR signature

To generate a consensus differential protein activity signature for breast CSLCs, we integrated the NES of each protein, as assessed by metaVIPER by comparing the 20 cells with the highest (most CSLC) and 20 cells with the lowest (most differentiated) marker-based stemness score across each patient-derived sample and the PDX-derived sample. The differential activity for each sample was computed using viperSignature() in VIPER11,12 R package, with the setting of the protein activity profiles for 20 cellsCSLC as a test and the protein activity profiles for 20 cellsDIFF as a referential state. Then, the z-score output by viperSignature() was averaged over cells for each sample. Justifying the generation of a consensus protein activity signature, we noted that the most differentially active proteins, as assessed in each sample, were exceedingly conserved across all samples (Suppl. Fig. S12B). Subsequently, the differential activity of each protein was integrated across the 7 patients and the control PDX model, using weighted z-test (zw):

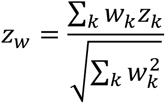

where zk is the differentially active protein for sample k and wk is a weight assessed as the inverse of the standard deviation of sample k. Thus, the transcriptional regulator proteins with the most positive (activated) and negative (inactivated) integrated NES in zw were selected as candidate MR proteins controlling the CSLC vs. differentiated states of breast cancer cells.

### Stemness Marker Analysis

in addition to the above-discussed established CSLC markers in breast cancer, we also assessed stemness based on an additional set of markers previously associated with cell stemness (non-breast-cancer-specific). These include genes in the ALDH family40,41 , ABC family41, quiescent stem-cell markers (FGD542 and HOXB543), embryonic diapause44 and asymmetric cell division processes45 (Suppl. Table 3). To identify genes involved in asymmetric cell division we used GO:0008356, while genes involved in embryonic diapause were selected from Rehman et al., 202144. To assess enrichment of these gene sets in the breast CSLC signature, we used analytic rank-based enrichment analysis (aREA) algorithm introduced in the VIPER manuscript12, which considers the sign (over or under-expressed). For this analysis the weight of each gene was set uniformly to one.

### Assessing CSLC enrichment in CCLE cell lines

We downloaded the TPM-normalized expression data (CCLE_expression.csv as of 06/30/2022) for all breast cancer cell lines in the Cancer Cell Line Encyclopedia46 from the Cancer Dependency Map (DepMap)47. Protein activity was assessed by metaVIPER, using the TCGA and single cell breast cancer-specific networks described above. To generate a signature for metaVIPER analysis, gene expression values were log2-scaled and normalized by subtracting the median value and dividing by the median absolute deviation (MAD) across all CCLE-BRCA samples. Note that MAD smaller than 0.01 was set to 0.01 to avoid denominator values approaching zero.

To assess the ability of each cell lines to recapitulate the CSLC MR signature differential activity, we used the OncoMatch algorithm48,49 analysis, designed to assess the overlap between two differential protein activity signatures. Specifically, we measured the normalized enrichment score (NES) of the 25 most activated and 25 most inactivated proteins in the breast CSLC signature in proteins differentially active in each CCLE-BRCA cell line (Fig. S19A). Thus, statistically significant NES scores identify cell lines that recapitulate the CSLC MR signature while statistically significant negative NES score identify cell lines that recapitulate the signature of the most differentiated BRCA cells. For this analysis, we only used the differential activity of transcriptional regulator proteins (n = 2088).

### Single-cell Stemness Analysis of HCC38, HCC1143, MCF7, and HCC2157 cell lines

To identify candidate breast cancer cell lines for the study, we assessed the stemness of 62 breast cancer cell lines in CCLE based on the enrichment (NES) of their most differentially activated proteins in the CSLC MR signature. As also supported by literature evidence50-52, we then selected HCC1143 (ranked 1) and HCC38 (ranked 3) as most enriched in the stem-like progenitor compartment and HCC2157 (ranked 49) and MCF7 (ranked 56) as basal and luminal cell lines most enriched in differentiated cells.

For efficiency purposes, we pooled HCC38 and HCC1143 cells and MCF7 and HCC2157 cells together in single-cell RNA sequencing. Cells were then demutiplexed using Demuxlet53. Briefly, this algorithm computes the statistical likelihood of a cell belonging to a specific sample based on the overlap of single-nucleotide polymorphisms (SNPs) between scRNA-seq reads and the referential Variant Call Format (VCF) data. For this analysis, we generated VCF files for each cell line from the CCLE raw data, as downloaded from the Sequence Read Archives54 (SRA), see below. After removing poor quality cells (MT% ≥ 7.5%, nReads ≥ 200,000, nGenes ≤ 5,000 for the HCC38/HCC1143 pool and MT% ≥ 12%, nReads ≤ 10,000 or nReads ≥ 90,000, and nGenes ≤ 3,000, for the MCF7/HCC2157 pool), we ensured that single cells from the 4 cell lines were clearly separated into distinct clusters in a UMAP projection, with each cluster associated with a specific cell lines by Demuxlet. The integrated stemness score of each single cell was then computed in the same manner as described above for the patient and PDX-derived samples.

Variant Discovery Analysis

For variant discovery analysis, used to deconvolute the pooled scRNA-seq profiles, we downloaded the CCLE raw data (FASTQ files) for the HCC38, HCC1143, MCF7, and HCC2157 cell lines from SRA (accession IDs: SRR8615458, SRR8615819, SRR8615758, and SRR8615891). We generated BAM files, with read sequences aligned and mapped to the human genome reference (GRCh38; Homo_sapiens.GRCh38.dna_sm.primary_assembly.fa as of 12/12/2022) from Ensembl55, using CellRanger56, and used the GATK57 (Genomic Analysis ToolKit) for variant calling analysis to generate the VCF files for each cell line. We applied SplitNCigarReads, a GATK subroutine, to split reads that contain Ns in their spliced alignment from the BAM files, before variant calling.

### CROP-seq Data Analysis

We used CellRanger to analyze the raw scRNA-seq data produced by the CROP-seq assay, for feature barcode analysis. The minimum UMI thresholds to call cells harboring a specific sgRNA was determined based on a gaussian mixture model by CellRanger’s CRISPR analysis for each sgRNA58. Cells with sgRNA UMI counts below the threshold were ignored as noise. Subsequently, poor quality cells were removed using the following criteria: For HCC38: MT% ≥ 9%, nReads ≥ 80,000, and nGenes ≤ 4,000; for HCC1143: MT% ≥ 10%, nReads ≥ 120,000, and nGenes ≤ 5,000). Cells with more than one sgRNA detected were excluded from the analysis.

To increase read depth and the number of detected genes in cells where a specific MR was knocked out, we used a pseudo-bulk59 approach, in which UMI counts were aggregated from the cells detected with the same sgRNA. This was critical to support effective differential protein activity analysis to assess the stemness score associated with knock-out of each MR compared to cells harboring intergenic control sgRNAs.

The pseudo-bulk expressions were CPM-normalized and transformed by log2-scaling. Then, the differential gene expression between a target sgRNA and the intergenic sgRNAs was computed by subtracting the median value and dividing by the median absolute deviation (MAD) across the expression of pseudo-bulks harboring intergenic sgRNAs (n=15). Note that MAD smaller than 0.01 was set to 0.01 to avoid denominator values approaching zero. Then, differential protein activity, as induced by CRISPR/Cas9-mediated, KO was assessed by metaVIPER analysis of the differential gene expression signature, using the TCGA-BRCA bulk and PDX-derived single-cell networks. MetaVIPER-assessed protein activity was converted into z-score, using the viperSignature() function in the VIPER12 R package. Specifically, viperSignature() was used to compute the z-score of the protein activity of the pseudo-bulk harboring a target MR sgRNA, compared to the protein activity of pseudo-bulks harboring intergenic sgRNAs (n=15), used as a reference control.

For each MR, effective KO was assessed by determining whether its activity in samples harboring the MR-targeting sgRNAs was lower than its average activity in all samples containing intergenic sgRNAs or sgRNAs targeting other MRs, by at least one standard deviation. Only MRs passing this test were further considered in the analysis. As a result, only 16 of 25 MRSCLC (BMPR1A, MTDH, ZNF131, MAML3, GON4L, ZNF24, SMAD5, KLF3, UBP1, SMAD1, TMF1, XBP1, MIER1, VEZF1, ETV3, ZNF566) and 9 of 25 MRDIFF (PCBD1, RUVBL2, HDGF, RPS3, RORC, ENY2, PEX14, THAP8, PARK7) were further analyzed in HCC38 cells, respectively. Conversely, only 15 of 25 MRSCLC (STAT3, BMPR1A, MTDH, ZNF131, GON4L, MYBL1, SMAD5, UBP1, NCOA1, SMAD1, TMF1, XBP1, VEZF1, ETV3, ZNF566) and 11 of 25 MRDIFF (PCBD1, RUVBL2, HDGF, PRDX2, YBX1, RORC, LAMTOR5, ENY2, THAP8, HLX, PARK7) were evaluated in HCC1143 cells, respectively.

The stemness NES of the cell state induced was computed by weighted enrichment analysis of the top 25 activated and 25 inactivated MRs following MR KO in proteins differentially active and inactive in the patient-derived stemness MR signature, using the aREA() function in the VIPER package. Specifically, the top 25 activated MRs were weighted positively (+1), whereas the top 25 inactivated MR were weighted negatively (-1) in the aREA analysis. To avoid biasing the analysis, each knocked-out MR was excluded from the corresponding aREA analysis, such that only its downstream effectors were considered. The statistical significance of the change in Stemness Score (NES) in each group (MRCSLC and MRDIFF) was evaluated by Mann Whitney U Test, compared to the Stemness Score of the control group.

Next, to assess the reprogramming potential of individual MRs, we generated bootstrapped pseudo-bulk expression profiles for each sgRNA. In short, cells containing sgRNAs targeting the same MR, as well as all cells containing the pool of intergenic control sgRNAs, were resampled 100 times with replacement to generate pseudo-bulk profiles. The latter were then normalized and used for protein activity inference, as discussed above. The Stemness Score (NES) of each bootstrapped sample was computed and the overall differential Stemness Score was assessed by Mann Whitney U Test, by comparing the NES of bootstrapped MR-KO samples vs. controls (intergenic sgRNAs).

### Cell fitness and gene dependency score

To assess cell fitness after CRISPR/Cas9-mediated MR KO, we first assessed the log2-scaled fold change (log2FC) of sgRNA abundance between the CROP-seq profiles and CRISPR library. Note that sgRNA counts were normalized by dividing the count of each sgRNA by the total sgRNA counts in the CROP-seq and CRISPR library, respectively. Then, we assessed MR essentiality by computing a gene dependency score, in which copy number-based bias was corrected in assessing cell fitness effects using the CERES60 R package. Briefly, we utilized the CCLE copy number data, the gene annotation data from the consensus coding sequence61 (CCDS), and bowtie62 indices for human genome (hg19), as provided in the CERES manuscript, to compute a gene dependency score from the log2FC values.

Note that we added the information of CCDS location for SMAD5 from Ensembl55 GRCh37 in the gene annotation data as it was omitted in the reference data provided in CERES.

### OncoTreat Analysis

To identify CSLC MR-inverter drugs, we used the extensively validated OncoTreat algorithm48,49,63,64. For each sample, the CSLC MR signature was assessed by metaVIPER analysis of the 20 cells with the highest vs. the 20 with the lowest stemness score in each of the 7 patient samples, using the viperSignature() function in the VIPER11,12 R package. . Stemness was assessed by integrating the 14 established breast cancer CSLC markers and the CytoTRACE scores. Finally, the z-scores representing the differential protein activities across the 7 patient samples were integrated using the weighted z-test, as described above.

For the OncoTreat analysis we leveraged drug perturbation profiles generated by PLATE-seq profiling65 of BT-20 cells at 6h and 24h following perturbation with 90 FDA-approved or late- investigational stage compounds, as described in64. Each compound was titrated at its 48h GI20 (the drug concentration at which 20% of the max inhibition of growth is achieved), as determined by 10-point dose response curves. To assess MR-inversion potential of a drug, we assessed the enrichment of the 25 positive CSLC MRs and 25 negative CSLC MRs in proteins that were inactivated and activated in drug vs. vehicle control treated cells. Drugs were then ranked based on their NES, with the top candidate MR-inverter drug having the most negative NES value. Consistent with previous OncoTreat publications48,49,63,64, we only considered direct transcriptional regulators (n = 2,088) as candidate MRs.

Bi-clustering of BT-20 perturbation profiles

To identify drugs inducing similar protein activity-level effects in BT20 cells we performed bi-clustering of the perturbational profiles generated at 24h and 1/10 GI20 concentration, after gene expression was converted to differential protein activity by metaVIPER analysis, by comparing each treated sample to the pool of vehicle control treated samples. For this analysis, only statistically significant differentially expressed regulatory proteins (TFs/co-TFs) (p < 10-5, n = 851) were considered. Bi-clustering was performed using complete hierarchical clustering method based on perturbational protein activity profiles Spearman’s correlation.

### GO enrichment analysis

Following bi-cluster analysis, 5 MR clusters were identified using the cutree() R function. These represent distinct subsets of MR proteins that are differentially activated or inactivated by specific drug subsets. To identify relevant biological pathways associated with each cluster, we performed enrichment analysis of biological pathway Gene Ontology (GO) terms, using the enrichGO() function in the clusterProfiler66 R package (see Supplementary Table 4).

### Drug Mechanism of actions in BT-20 data

Information on drug mechanism of actions (MoAs) of individual drugs in the analysis was obtained by manual analysis of the ChEMBL67 and DrugCentral68 databases (see Supplementary Table 5).

### Visualizing the differential cell density following drug treatment in vivo

To visualize cell density changes in drug vs. vehicle control-treated PDX samples, across the entire cell state space, we computed it over a principal component (PC) projection of their protein activity profiles. Single cells treated with either vehicle control, paclitaxel, or albendazole were visualized in independent plots. To avoid bias due to differential cell counts across different treatments, we selected 2,500 cells from each sample at random. Statistical significance of the differential cell densities in drug vs. vehicle-treated samples was assessed by computing the mean and standard deviation of the 2-D kernel density of each sample, by estimating optimal kernel density from 30% of the cells selected 100 times at random from each sample. The 2-D kernel density was computed, using the kde2d() function in the MASS69 R package, with bandwidth h = 0.01, n=50 grid points, and default bivariate normal kernel settings. The kernel density of each drug-treated sample was then converted to a z-score, using the statistical values from the control sample (i.e., vehicle treatment) as the null hypothesis. Therefore, a positive (or negative) z-score indicated an increased (or decreased) cell density in the drug-treated sample, compared to the control.

### Hallmark enrichment analysis of the PDX data

To assess the enrichment of 50 MSigDB hallmarks70 in the direction of the 1st and 2nd PCs of the PDX protein activity profiles, we first ranked the proteins based on the Spearman’s correlation between their activity and both the 1st and 2nd PCs, reflecting the greatest variance axes in the data. As a result, proteins were ranked based on the correlation between the ordering of cells along each PC and protein activity. Gene set enrichment analysis of the 50 cancer hallmark sets in each PC-specific ranked protein list was assessed, using the aREA() function in the VIPER R package. Cancer hallmark gene sets were obtained from the MsigDB database70.

### Assessing the Effect of Drug Treatment on the Ratio Between CSLCs and Differentiated Cells

To assess the effect of each drug treatment on cell stemness, we computed the ratio of most stem-like cells (stemness score > 0.8) to most differentiated cells (stemness score < 0.2) in drug vs. vehicle control-treated samples. Statistical significance was computed using the hypergeometric function (i.e., Fisher’s Exact Test).

### MR Modularity Assessment

For each of the top 20 candidate CSLC MRs, we collected its putative protein-protein interactions (PPIs) from the STRING71 and PrePPI72 databases and its putative regulatory interactions from ARACNe-AP13 analysis. The analysis identified n = 67 molecular interactions in the candidate module comprising these MRs. To determine whether the modularity was significant, we assessed the average connectivity of 1,000 sets of 20 regulatory proteins selected at random, using the same approach to determine PPIs and regulatory interactions. The degree distribution of these random modules (size = 20 proteins) was then fit to a negative-binomial (NB) distribution model, using the fitdistr() function from the fitdistrplus73 R package. The resulting parameters were mu = 13.2, which corresponds to the average expected number of interactions, and size = 5.80, corresponding to a dispersion of the NB distribution with p-value = 1, based on a Chi-squared goodness fit test. This null model was then used to assess the statistical significance of 20-MR modularity.

## ADDITIONAL RESOURCES

scPLATE-seq protocol: dx.doi.org/10.17504/protocols.io.s4hegt6

---

**Figure S1.**
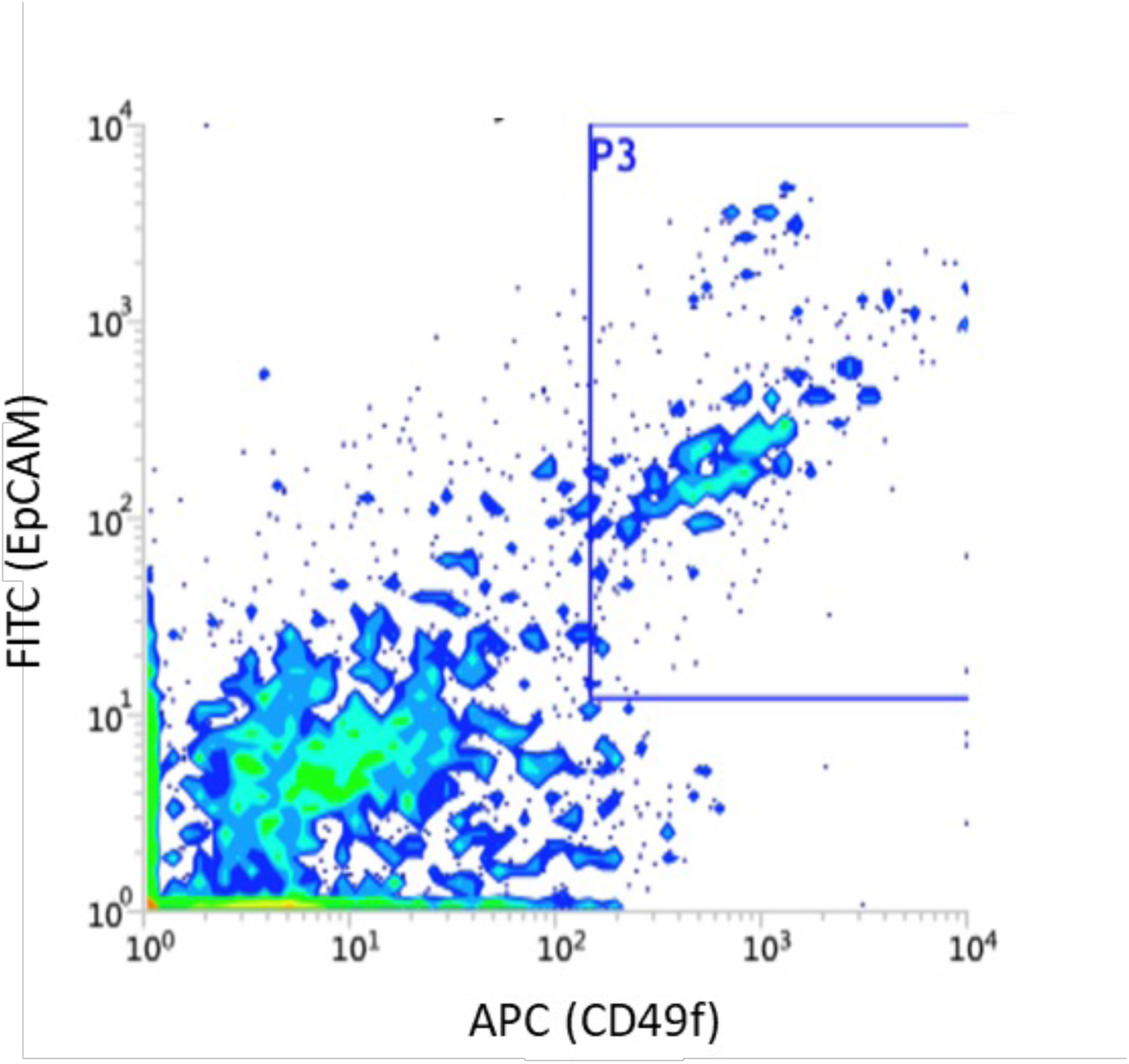
FACS of a human metastatic breast cancer tissue (patient S1). Malignant epithelial cells were isolated based on EPCAM^+^ and CD49f^high^ and subjected to scRNA-seq.

**Figure S2.**
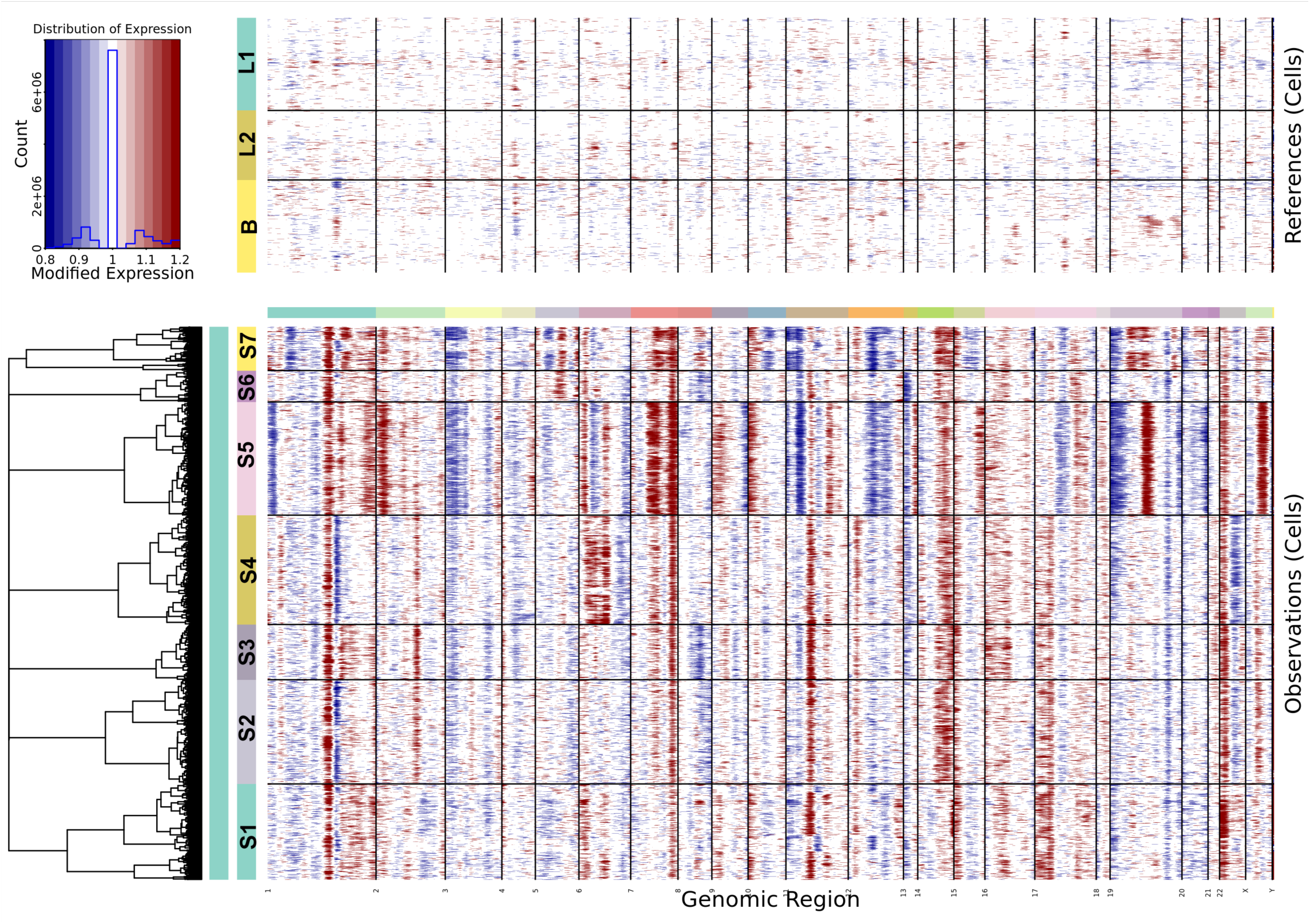
Inferred copy number variations in breast cancer patient samples (S1-S7) by inferCNV. Normal breast epithelial cells containing one basal (B) and two luminal (L1 and L2) subtypes from 4 individuals (GSE113196) were used as reference. In the patient samples, S1 and S2 were weakly HR+, S2, S3, and S4 were strongly HR+, and S5 and S7 were TNBCs, according to immunohistochemistry (IHC).

**Figure S3.**
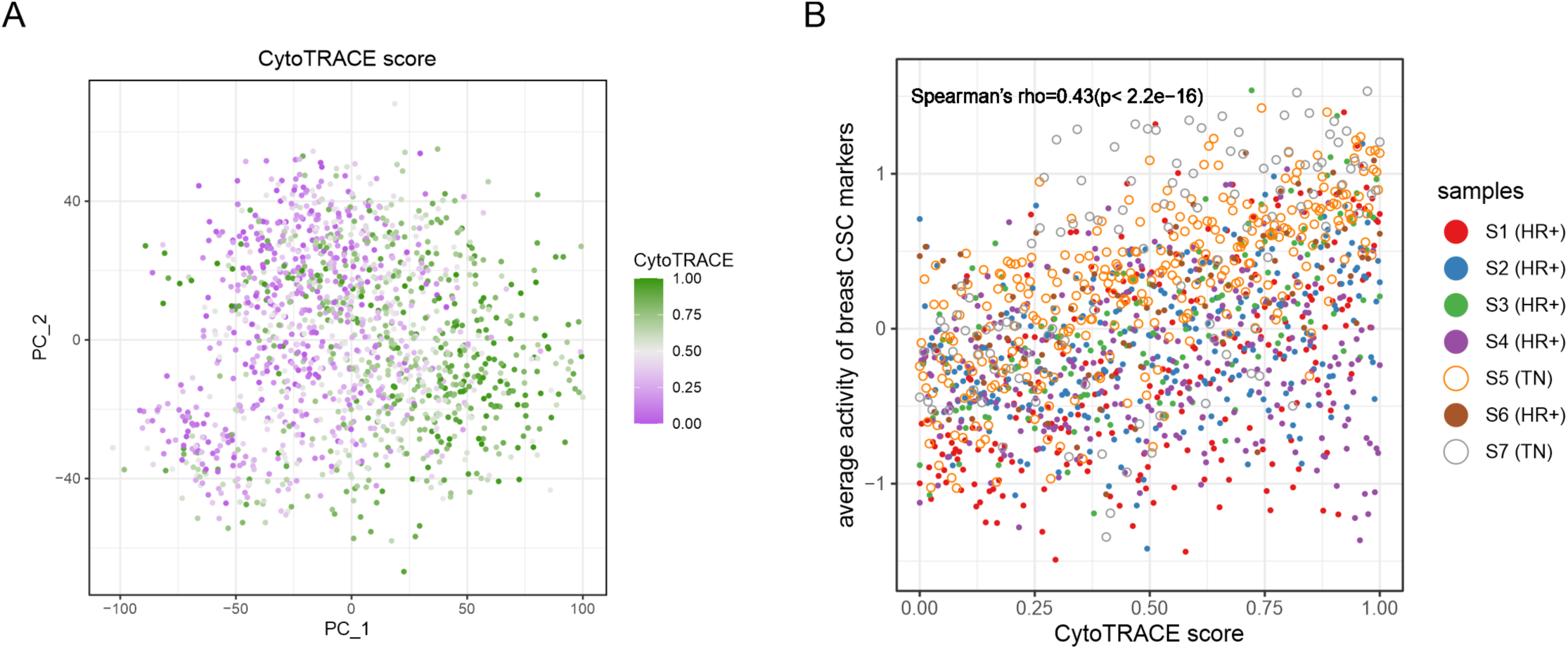
**A.** The CytoTRACE prediction in the patient scRNA-seq data. Cells were colored with a green-grey-purple gradient corresponding to the CytoTRACE score from one (least differentiated) to zero (most differentiated). **B.** The correlation between the CytoTRACE score and the average activity of breast CSC markers in the patient samples. The following proteins were considered as breast CSC markers in the patient data: CD44+/CD24-, ITGA6, BMI1, SALL4, NOTCH1, NOTCH2, KLF4, CTNNB1, ITGB3, ITGB1, PROM1, POU5F1, SOX2, and KIT. The correlation between CytoTRACE and the maker’s activity was statistically significant (Spearman’s rho=0.43, *p*-value<0.001).

**Figure S4.**
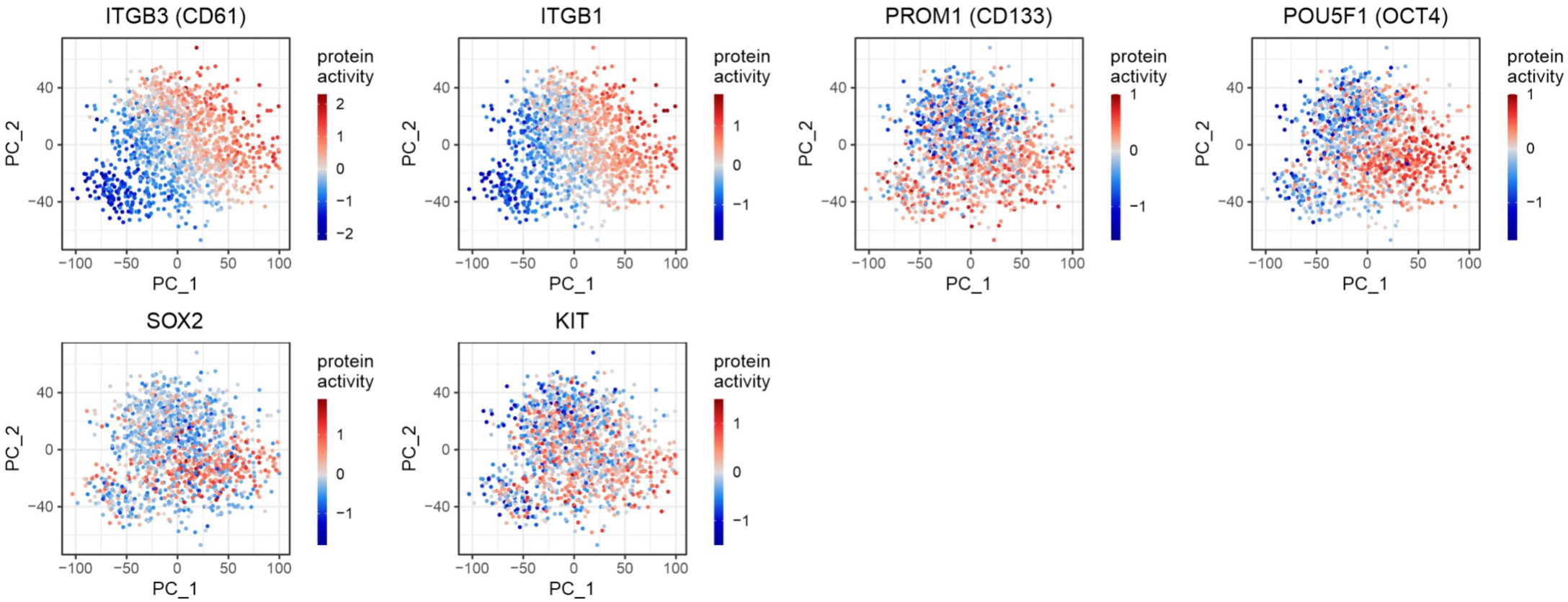
The inferred activity of several breast CSC markers, namely ITGB3, ITGB1, PROM1, POU5F1, SOX2, and KIT. Note that the VIPER-inferred activities were centered in this visualization.

**Figure S5.**
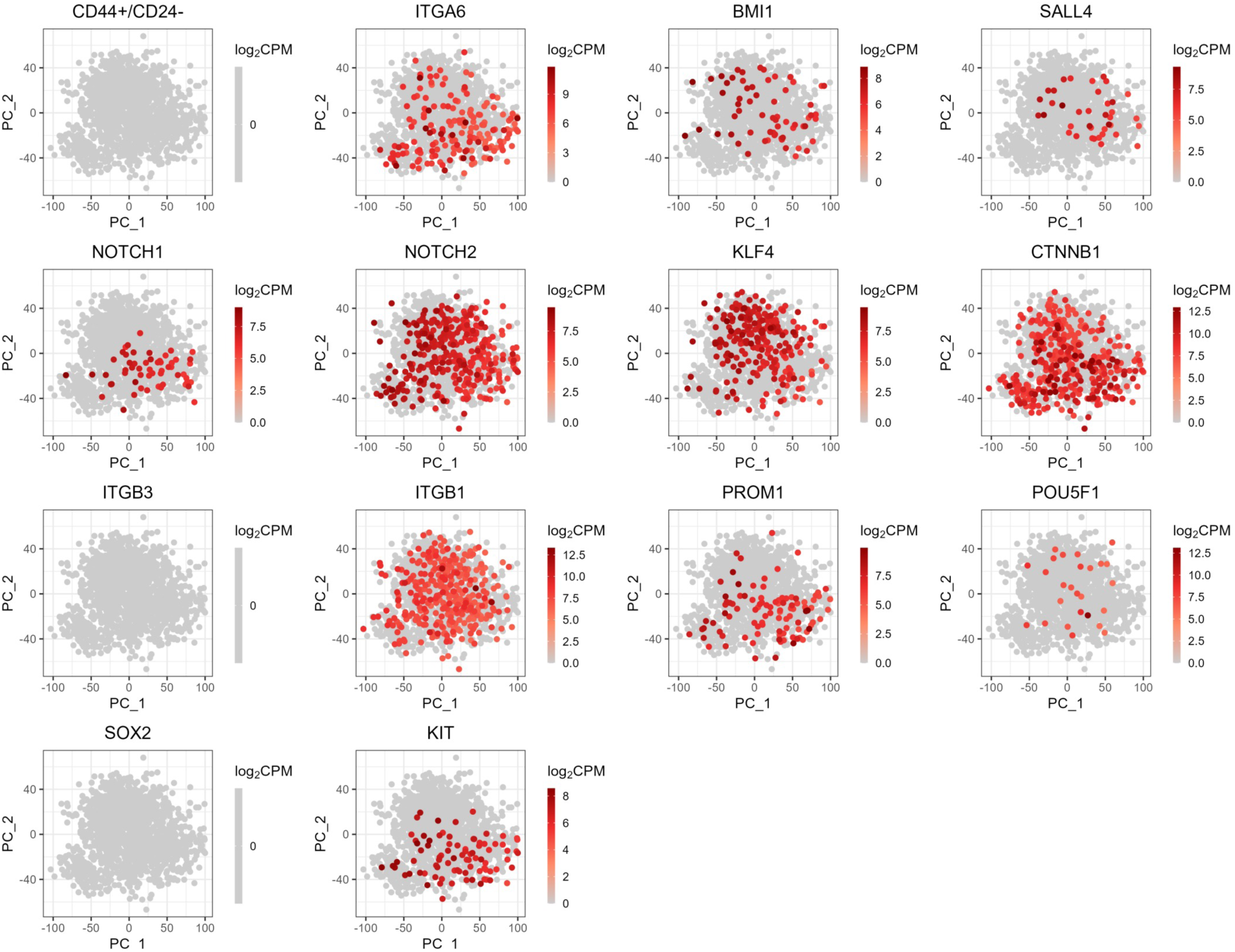
Gene expression (log_2_CPM) of known breast CSLC markers. The same PCA mapping based on the protein activity was used for the comparison. Note that only the gene expression greater than the upper quantile is shown for visualization purposes.

**Figure S6.**
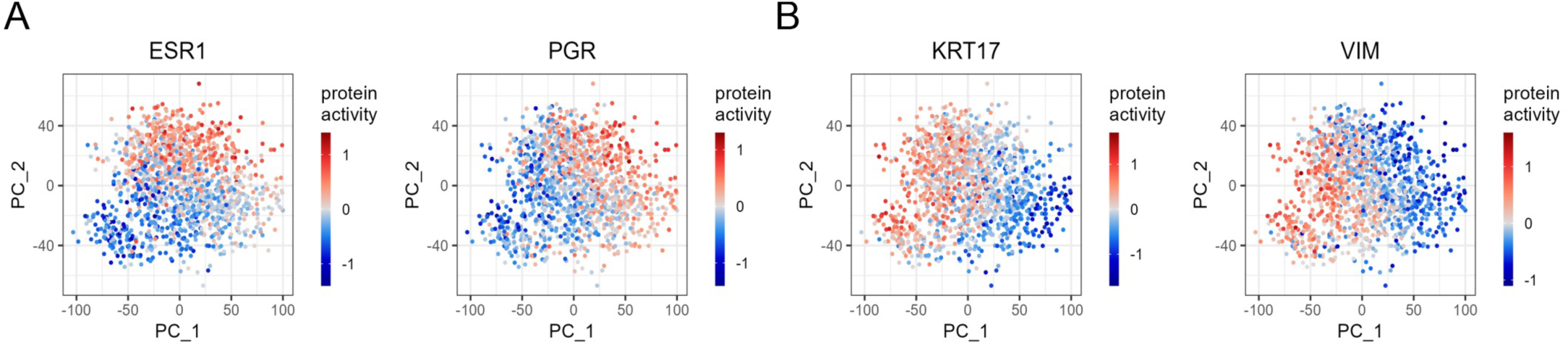
The inferred activity of breast cancer subtype markers. **A.** Two HRs (ESR1 and PGR). **B.** A basal-type marker (KRT17) and non-HR+ marker (VIM). While ESR1 and PGR activities are greater in HR+ patient samples, KRT17 and VIM activities are greater in TNBC samples. Note that the VIPER-inferred activities are centered in this visualization.

**Figure S7.**
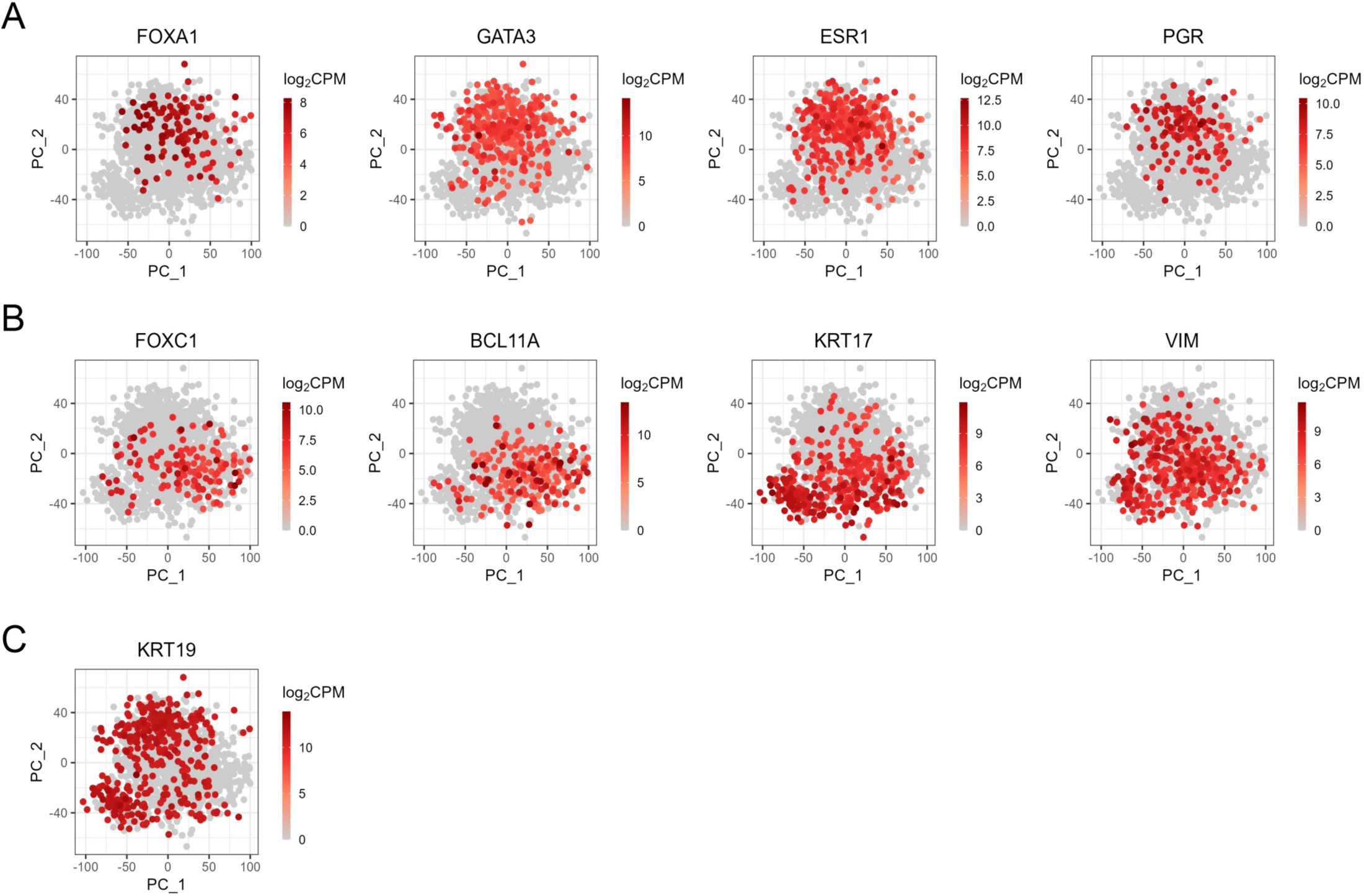
Gene expression (log_2_CPM) of known breast cancer subtype markers in the patient samples: HR+ markers (**A**), TNBC-specific markers (**B**), and a differentiated cell marker (**C**). The same PCA-mapping based on the protein activity was used for the comparison. Note that only the gene expression greater than the upper quantile is shown for visualization purposes.

**Figure S8.**
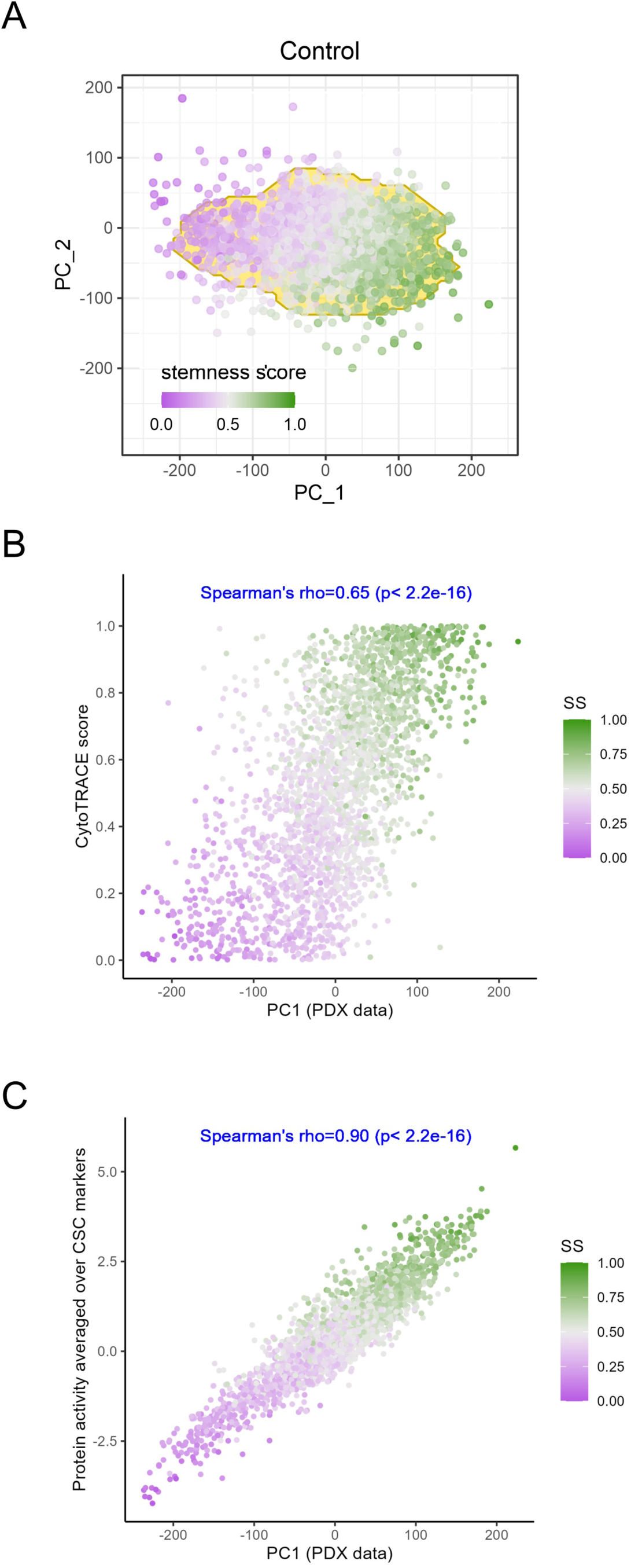
Protein activity-based cell clustering in the PDX control sample (i.e. vehicle treatment). The color of cells indicates the stemness score (SS) as described in Methods. **A.** The first two principal components (PCs) of the protein activity of cells. **B.** The correlation between the 1^st^ PC and the CytoTRACE. **C.** The correlation between the 1^st^ PC and the averaged activity of the 14 CSC

**Figure S9.**
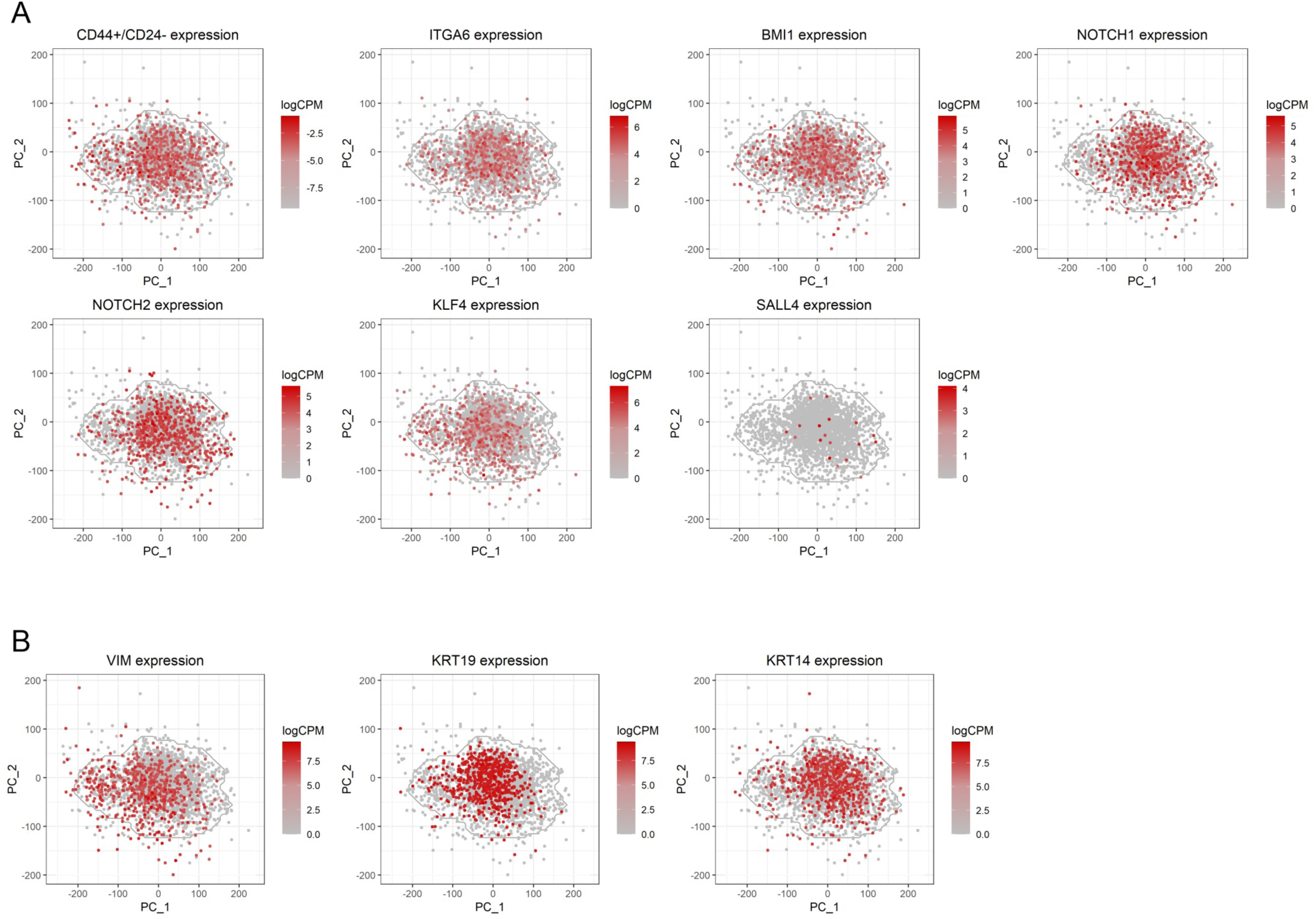
Gene expression (log_2_CPM) of known breast CSC markers (**A**) and differentiated markers (**B**) in the PDX control sample. Protein activity-based PCA-mapping was used for the comparison. Note that only the gene expression greater than the upper quantile was shown for a visualization purpose.

**Figure S10.**
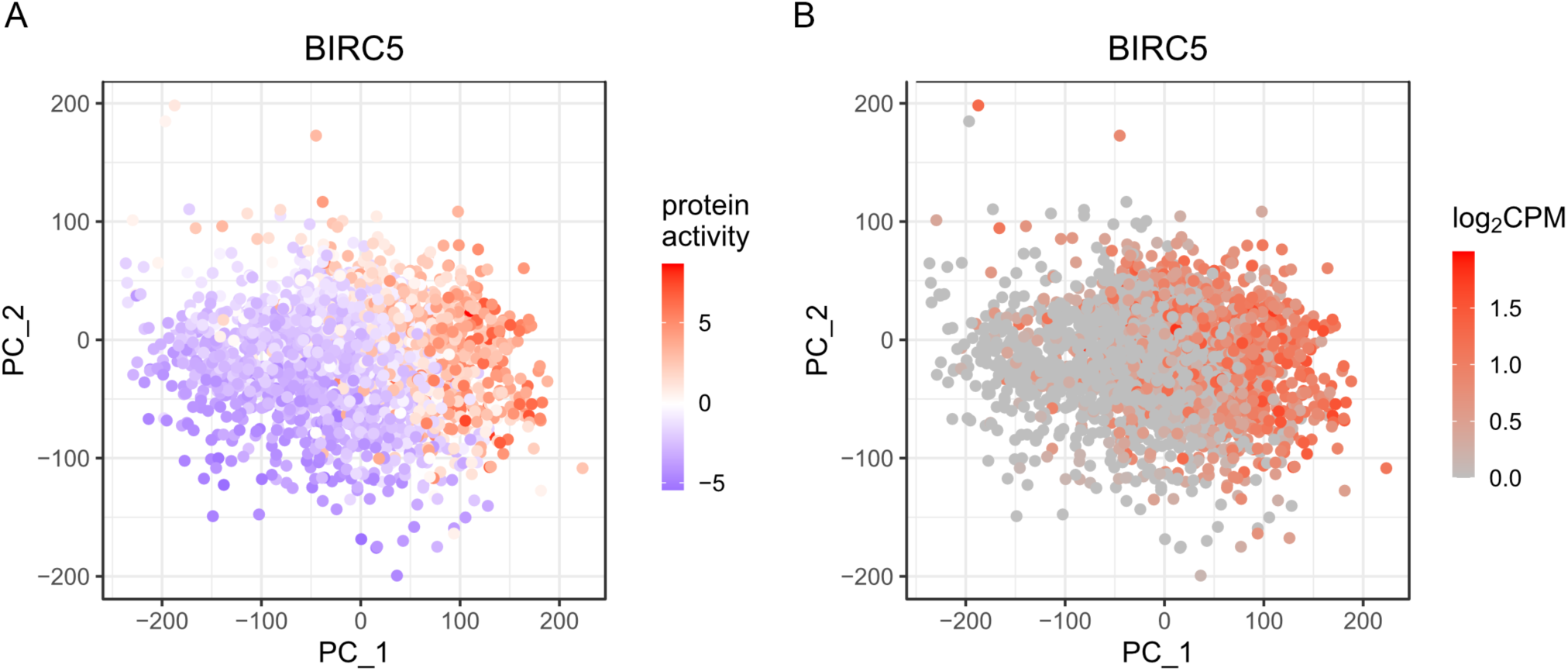
Protein activity (**A**) and gene expression (log_2_CPM) (**B**) of BIRC5, a marker of quiescent-breast CSCs.

**Figure S11.**
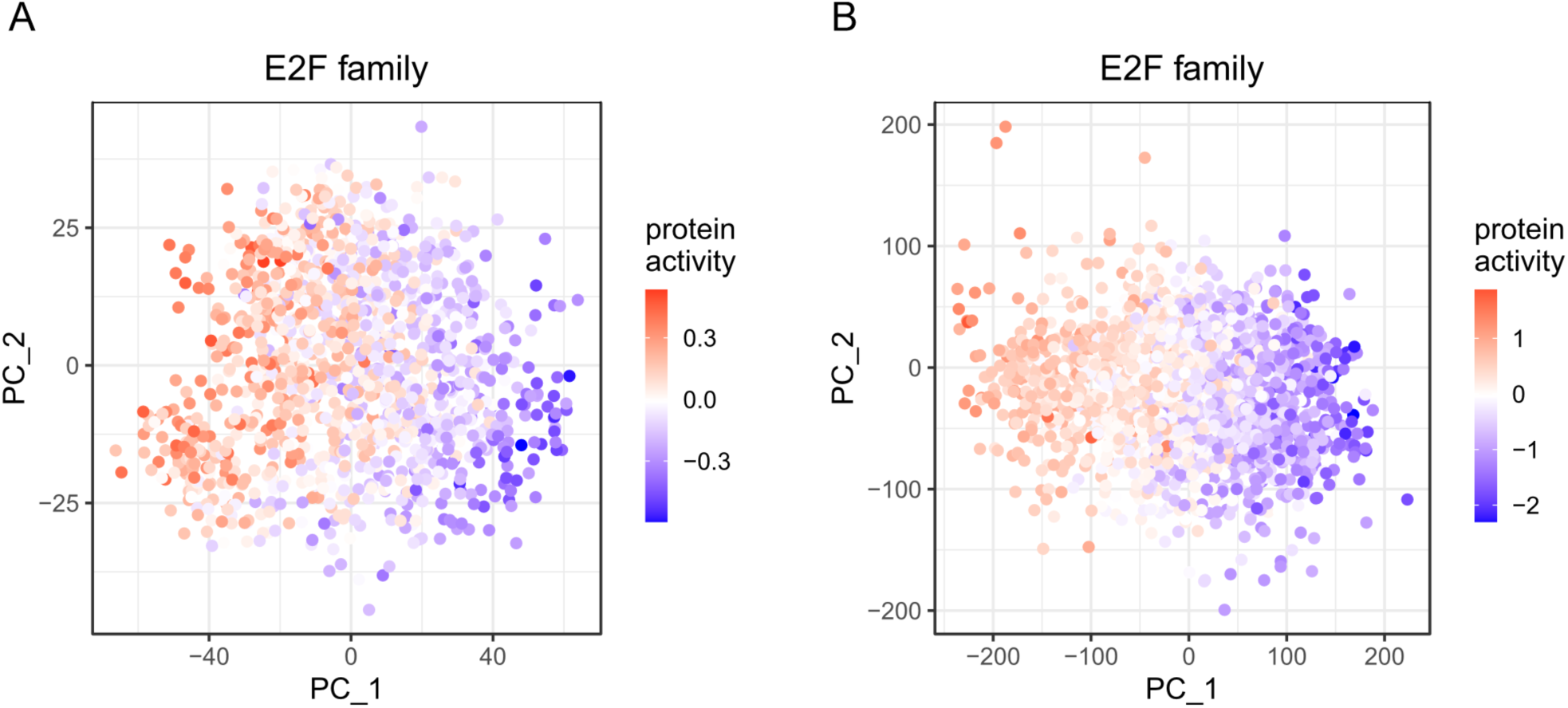
Protein activity of E2F family, a proliferative cell marker in the patient (**A**) and PDX control (**B**) data.

**Figure S12.**
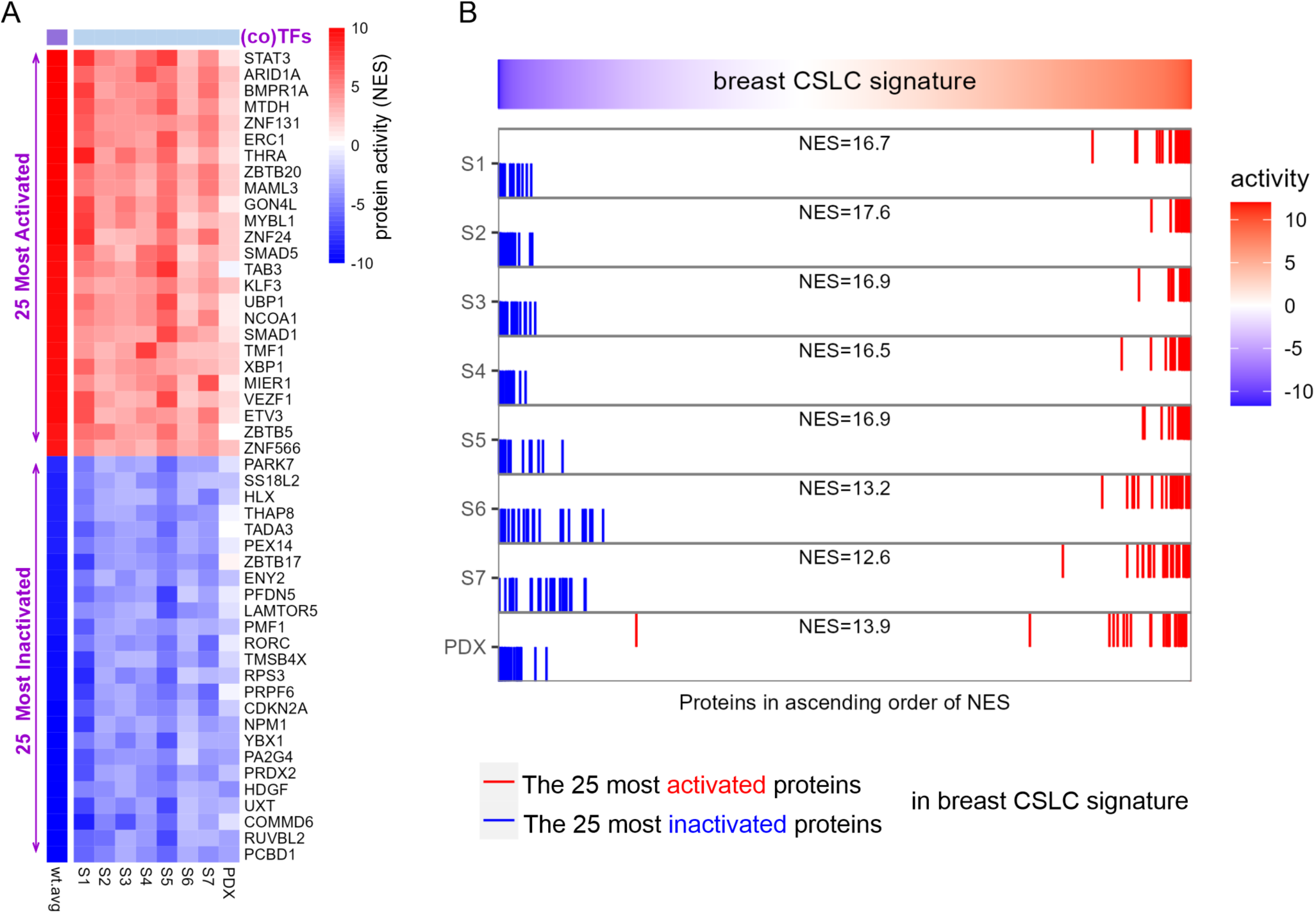
**A.** A heatmap exhibiting the activity of the 25 most activated and 25 most deactivated TFs/coTFs in the breast CSLC signature and their activities in individual samples (7 patient samples and the PDX vehicle-treated sample). **B.** An enrichment analysis plot of the 25 most activated and 25 most inactivated proteins of the CSLC signature in each individual sample. The consistency of the activity of these 50 proteins is observed in all samples, showing a significant enrichment (p-value< 1.0×10^-16^).

**Figure S13.**
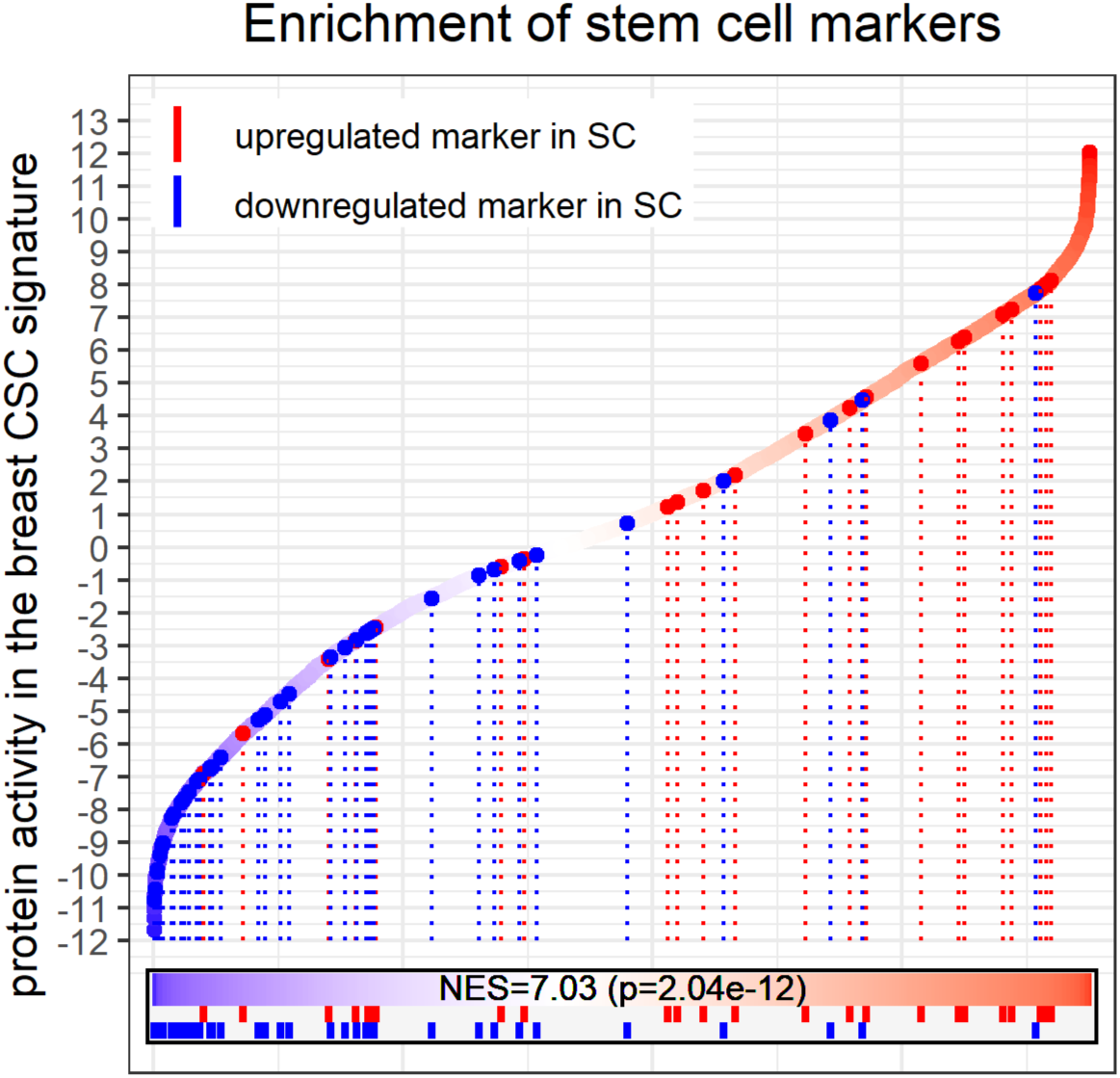
An waterfall plot of sorted protein activities in the breast CSLC signature, identified using both the patient and PDX control data. The NES of genes involved in general stem cell processes (supplementary data) is shown on the CSC signature.

**Figure S14.**
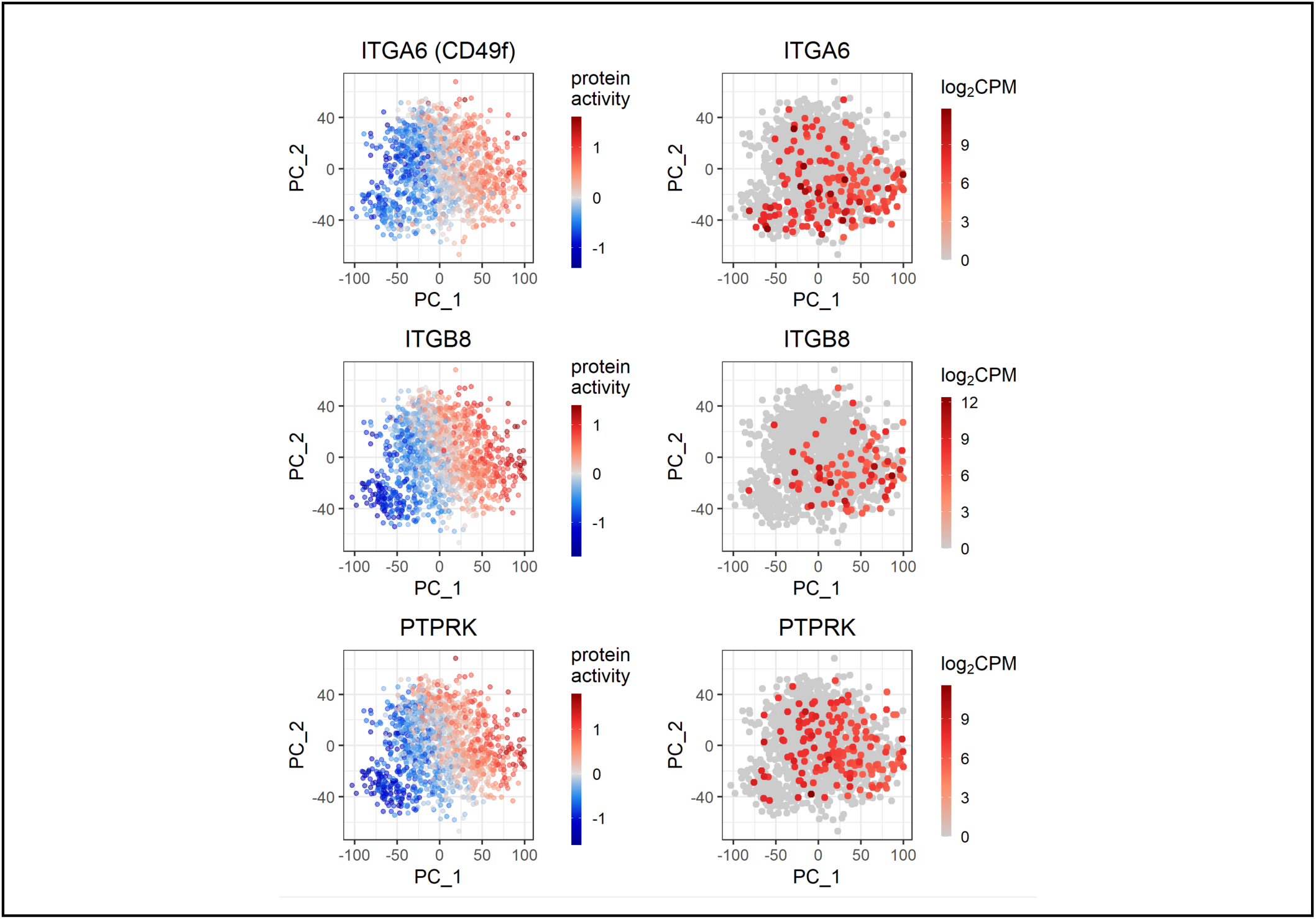
The VIPER-inferred activity (centered) and the gene expression (log_2_CPM) of cell surface proteins among the top 50 most activated proteins in the breast CSCs. For gene expression, only cells with expression greater than the upper quantile are shown for visualization purposes.

**Figure S15.**
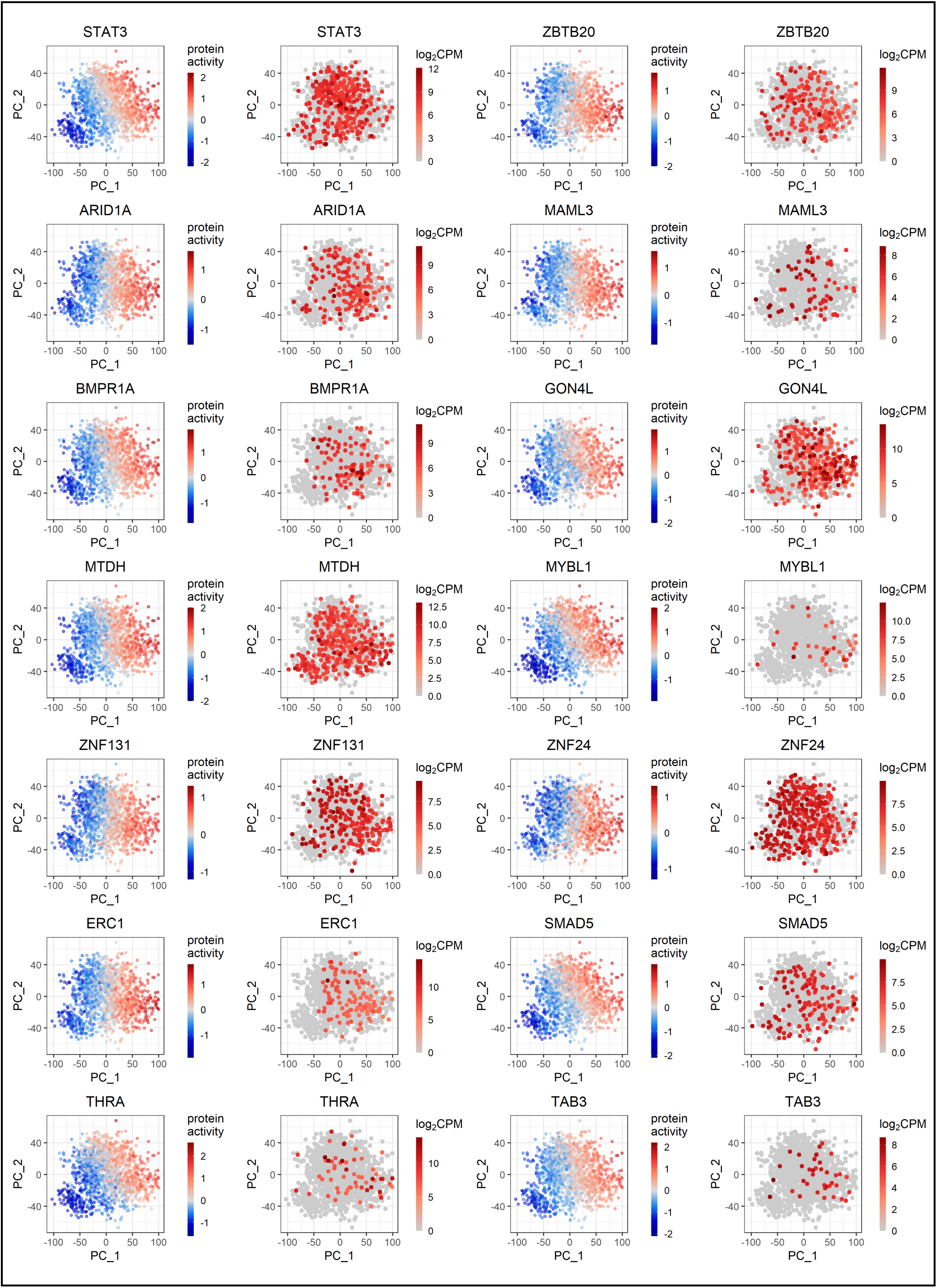
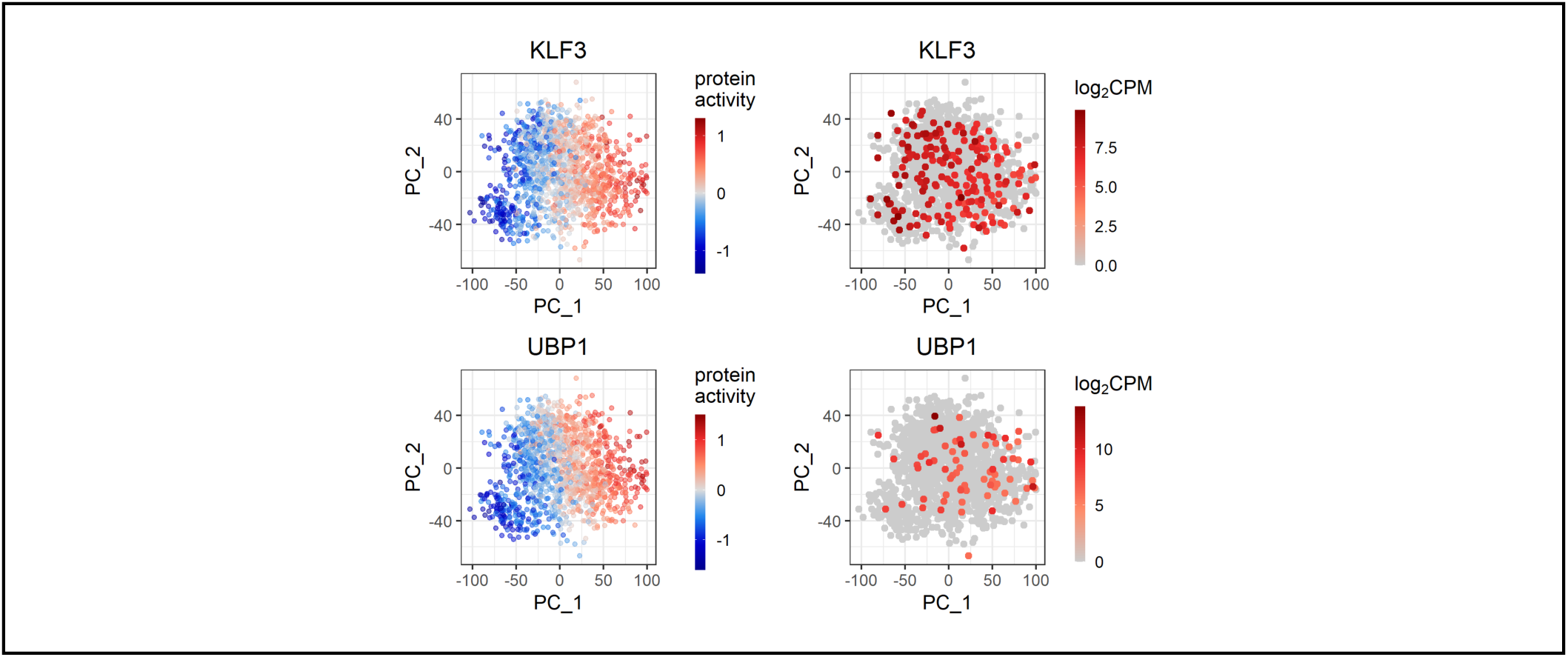
A. The VIPER-inferred activity (centered) and the gene expression (log_2_CPM) of transcription (co-)factors among the top 50 most activated proteins in the breast CSCs. For gene expression, only cells with expression greater than the upper quantile were shown for visualization purposes.

**Figure S16.**
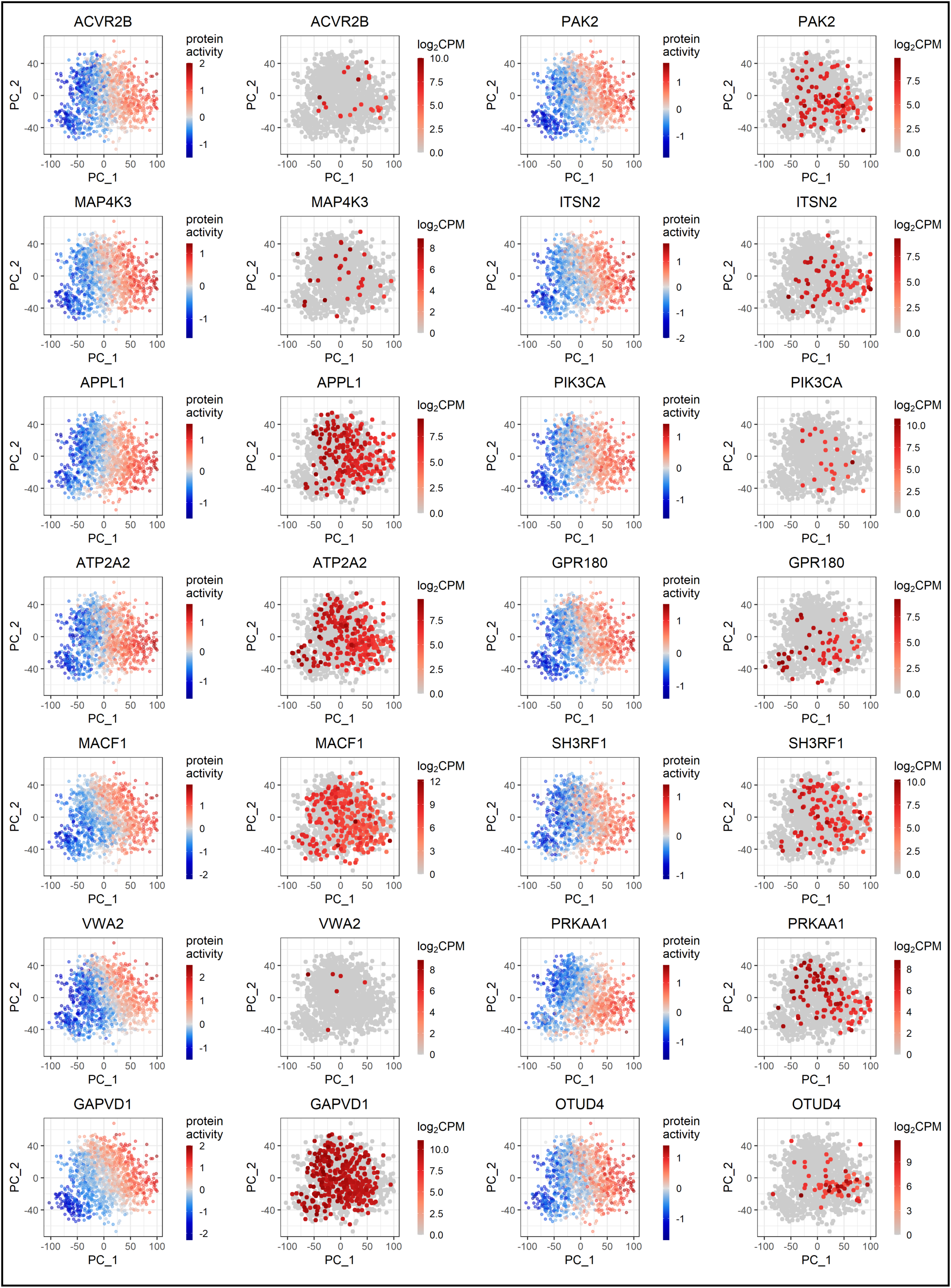
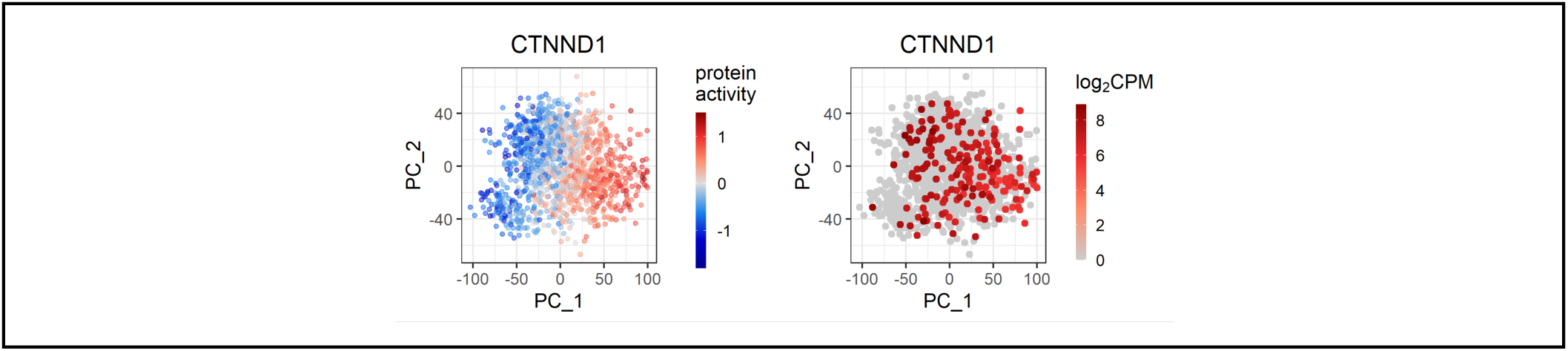
A. The VIPER-inferred activity (centered) and the gene expression (log_2_CPM) of signaling molecules among the top 50 most activated proteins in the breast CSCs. For gene expression, only cells with expression greater than the upper quantile were shown for visualization purposes.

**Figure S17.**
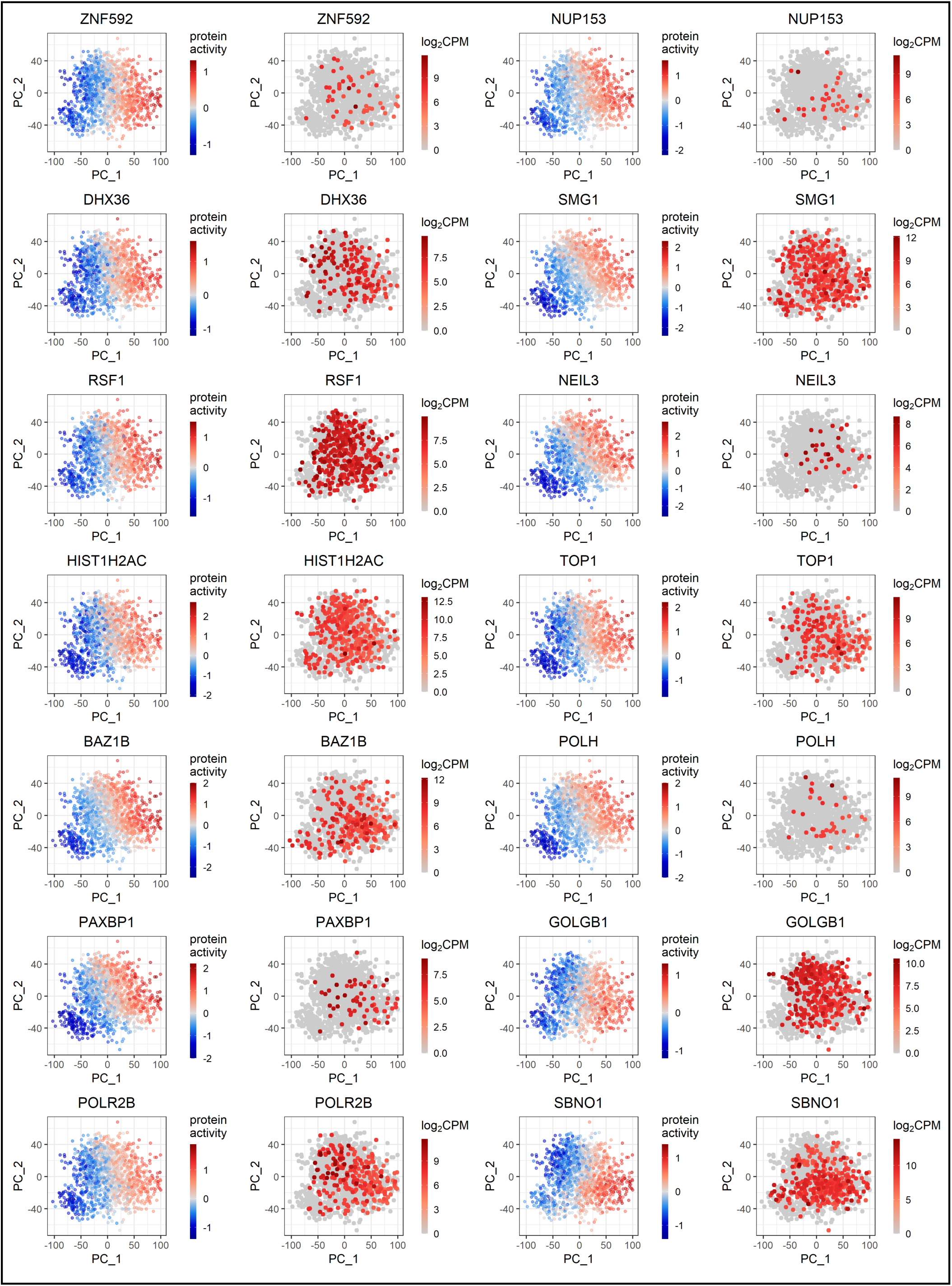
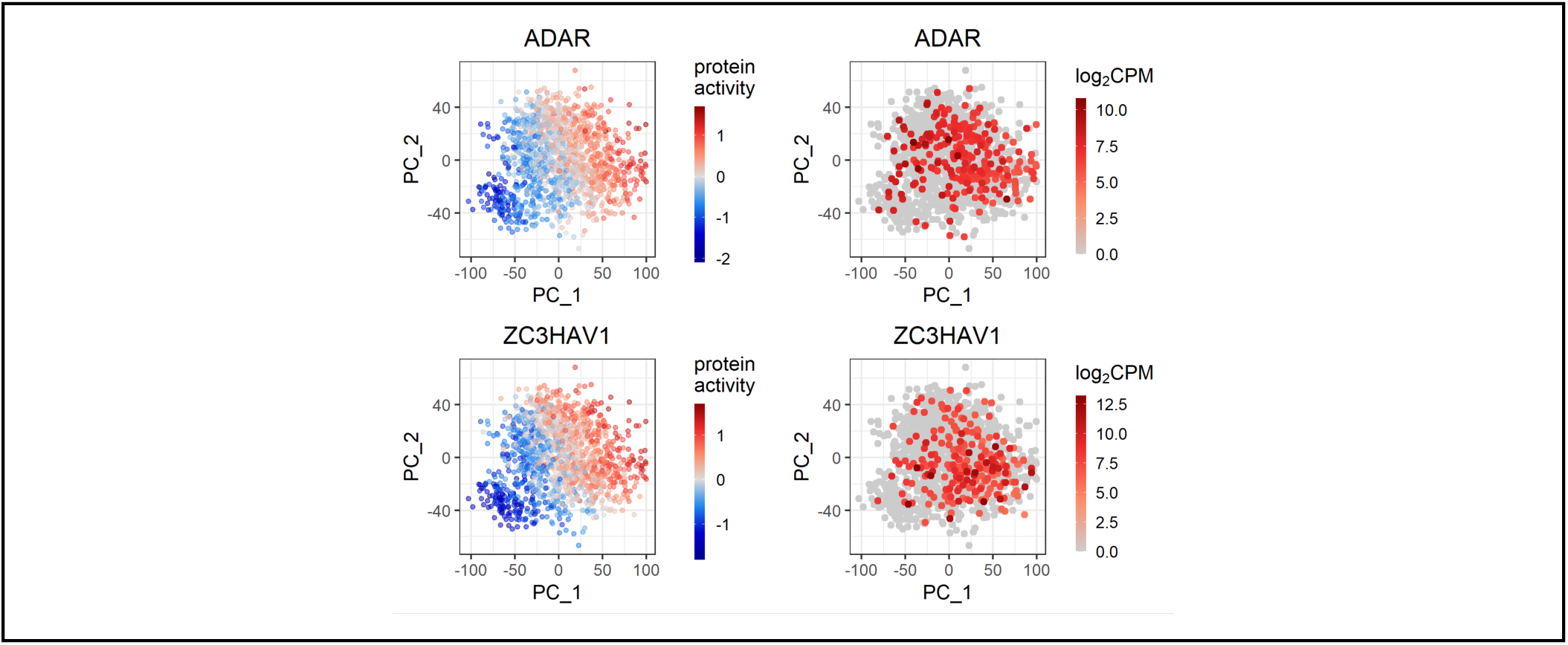
A. The VIPER-inferred activity (centered) and the gene expression (log_2_CPM) of other proteins involved in transcriptional programs among the top 50 most activated proteins in the breast CSCs. For gene expression, only cells with expression greater than the upper quantile were shown for visualization purposes.

**Figure S18.**
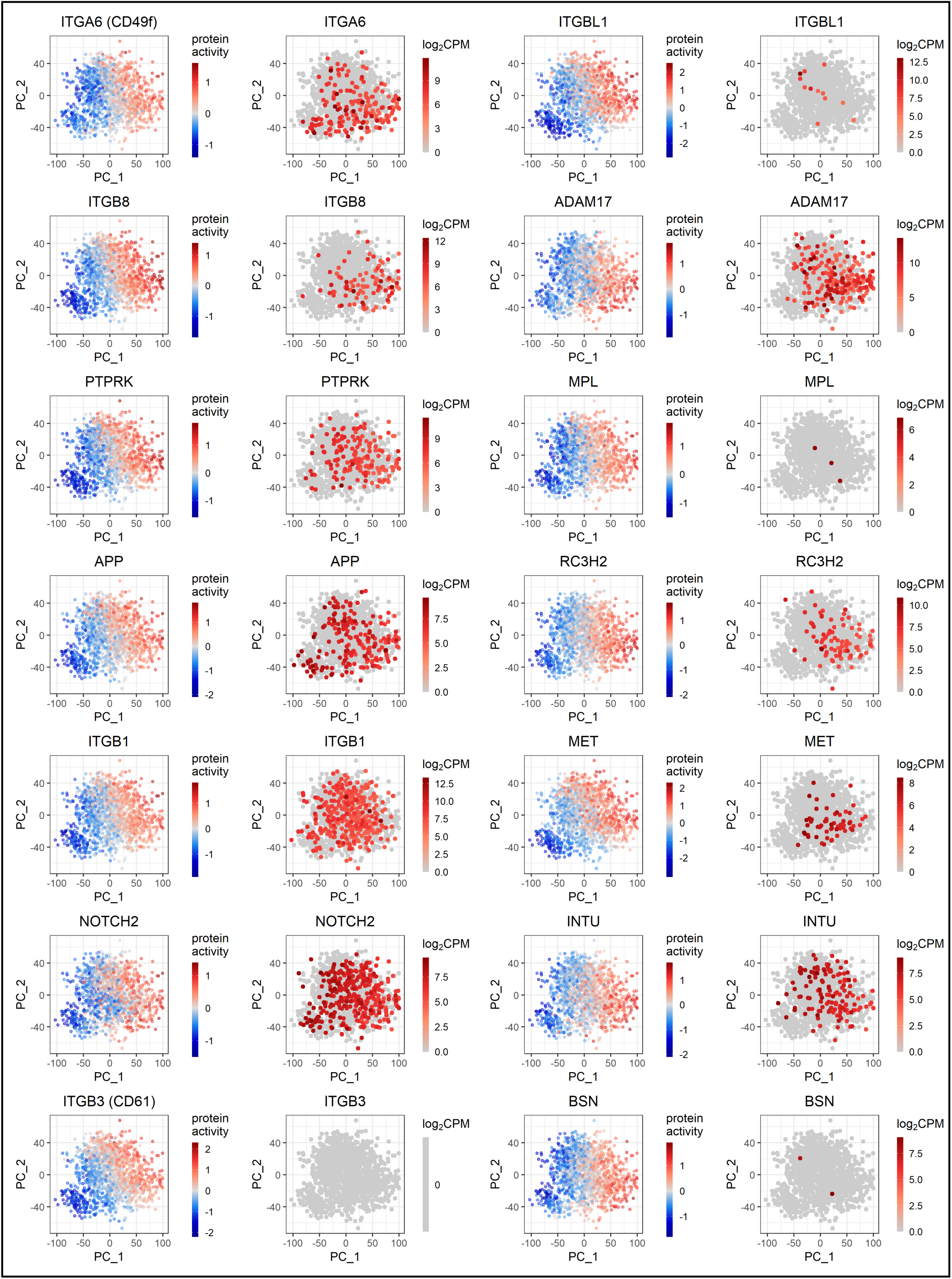
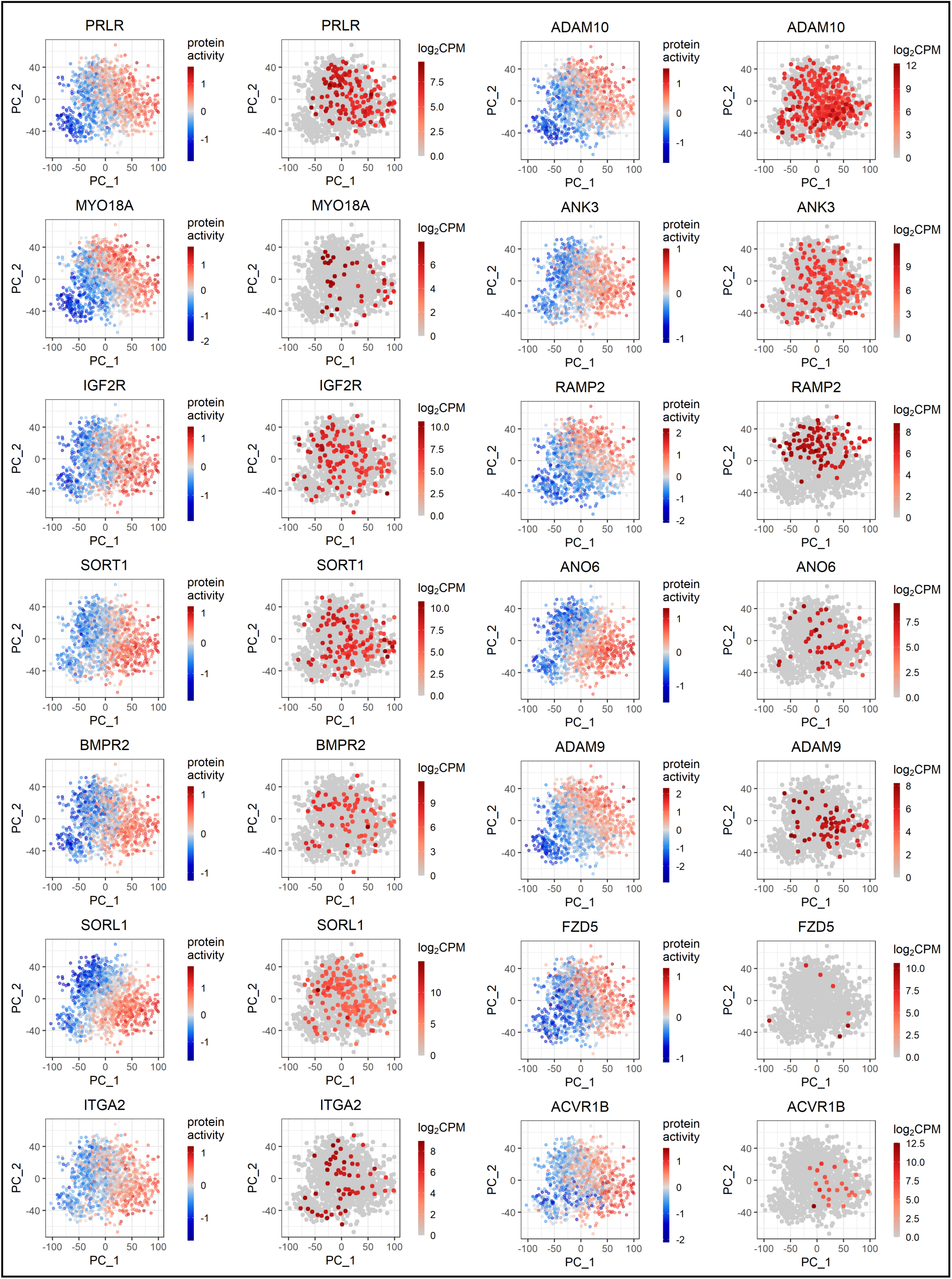
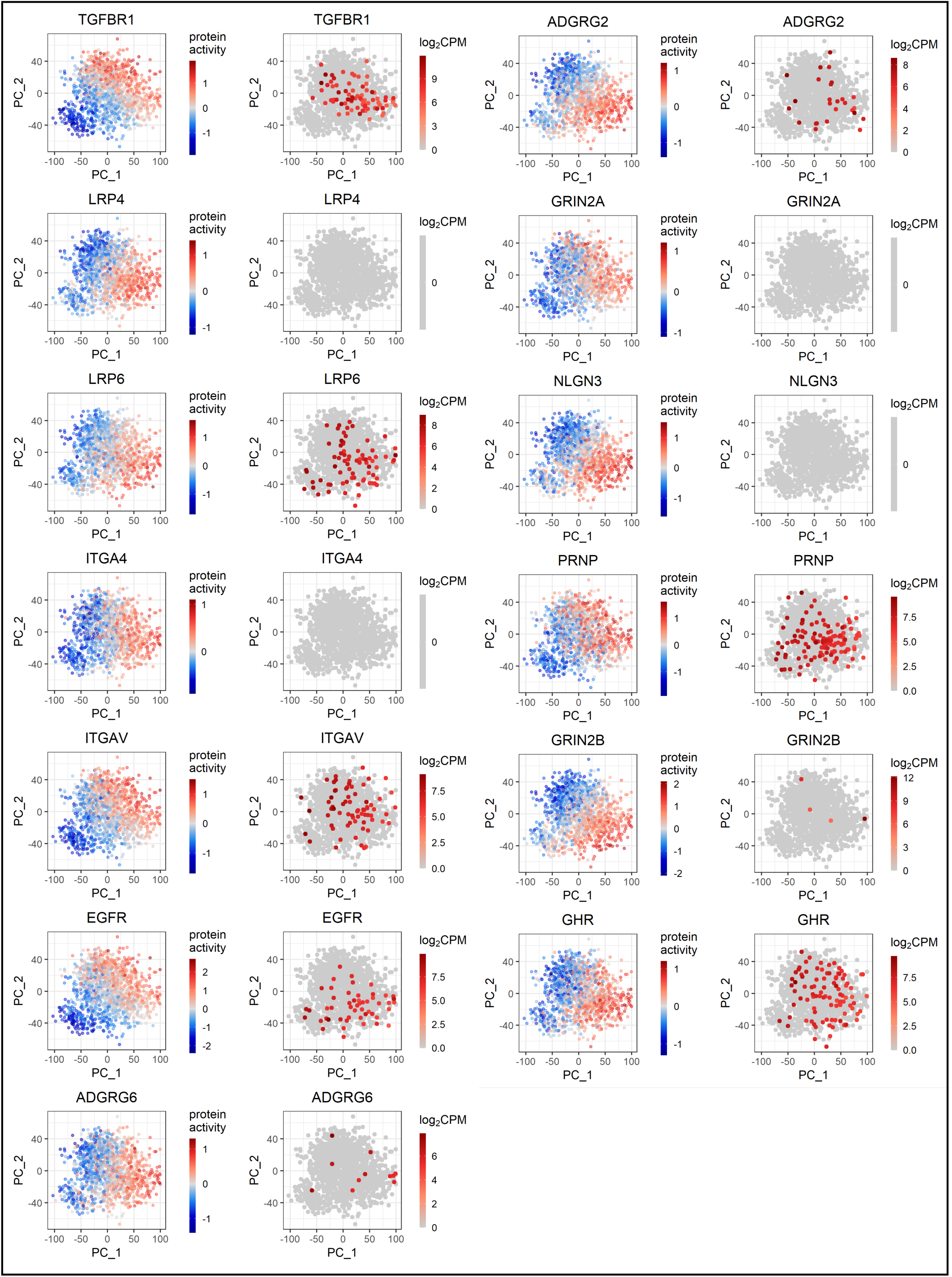
A. The VIPER-inferred activity (centered) and the gene expression (log_2_CPM) of cell-surface proteins among significantly activated proteins in the breast CSCs (Bonferroni adjusted *p*- value < 0.001). For gene expression, only cells with expression greater than the upper quantile were shown for visualization purposes.

**Figure S19.**
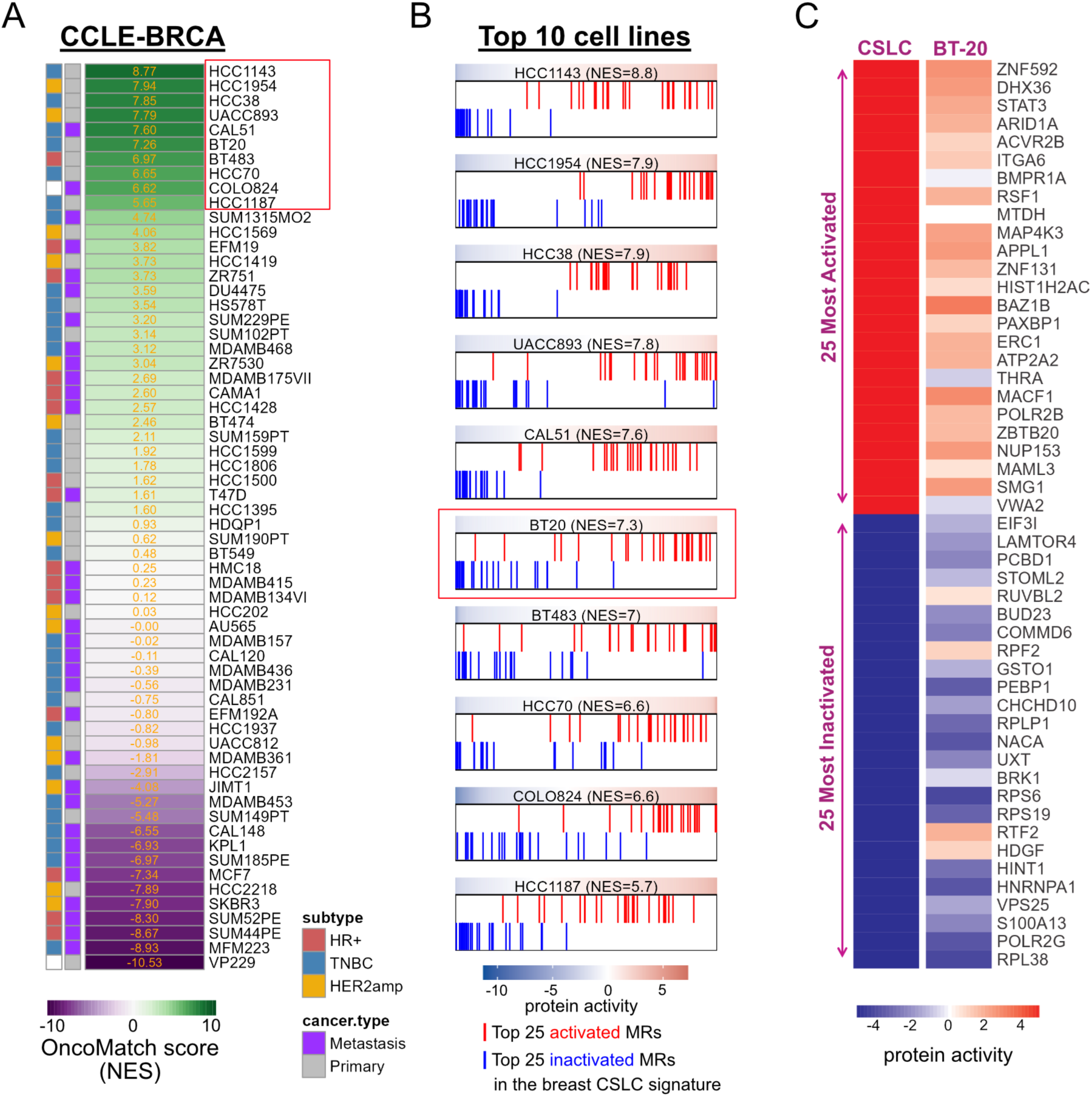
**A.** A heatmap illustrating the ranked list of breast cancer cell lines in CCLE, in a decreasing order of their OncoMatch score, which indicates the similarity score of the protein activity profiles between a cell line and the CSLC signature. Thus, the greater the OncoMatch score, the more stem-like cell properties. **B.** Enrichment analysis plots of the 25 most activated and the 25 most inactivated proteins in the breast CSLC signature for top 10 breast cancer cell lines by OncoMatch. The plot demonstrated a statistically significant enrichment (p-value< 1×10^-16^), between BT20 and the CSLC signature. **C.** A heatmap of the activity of the 25 most activated and the 25 most inactivated proteins in the breast CSLC signature and their activities in BT20.

**Figure S20.**
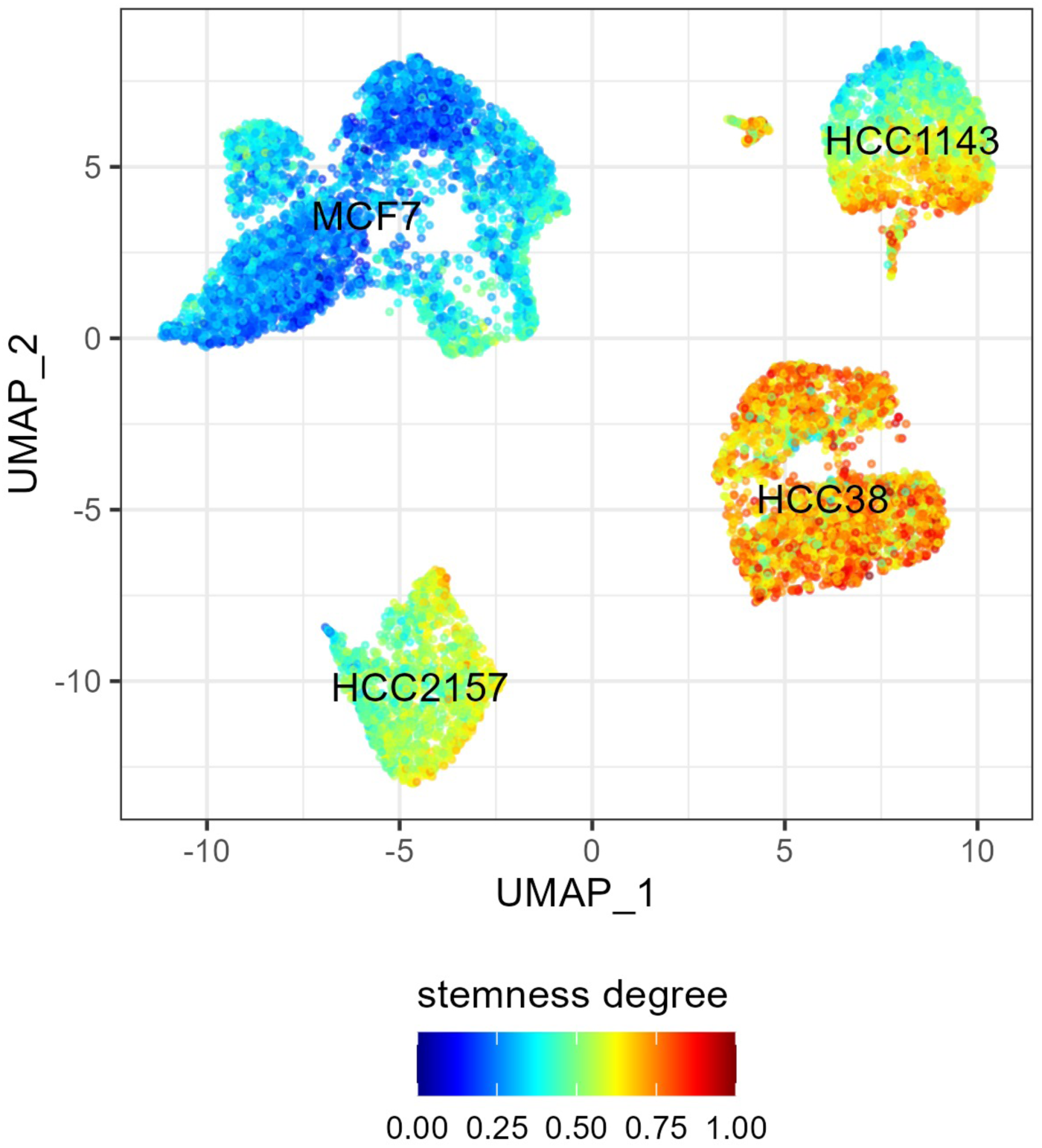
The UMAP projection of protein activity profiles of single cells for four breast cancer cell lines. HCC38 and HCC1143 are chosen for representing CSLC-rich cell lines, while HCC2157 and MCF7 are selected as well-differentiated cell lines (negative controls), based on our OncoMatch prediction as well as literature. The color of cells indicates the stemness degree calculated in the same manner as the previous (i.e. the weighted average of stemness marker activities and the CytoTRACE score). Thus, the greater score, the higher stemness in cells.

**Figure S21.**
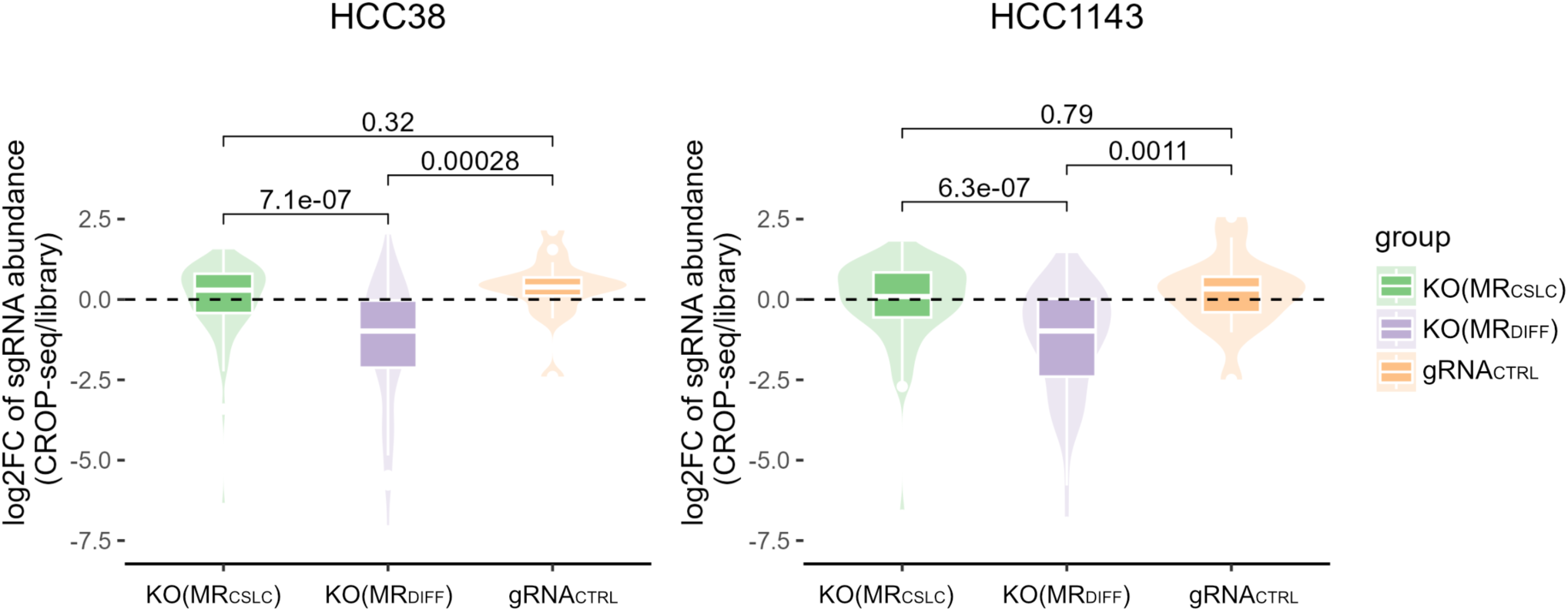
The log2 Fold Change (log2FC) of sgRNA abundance between CROP-seq and CRISPR library. sgRNA counts were normalized by dividing them by total sgRNA counts in each CROP-seq and CRISPR library. Unlike MR_CSLC_, log2FC after MR_DIFF_ KOs is significantly diminished, implying many of MR_DIFF_ are responsible for cell fitness/proliferation.

**Figure S22.**
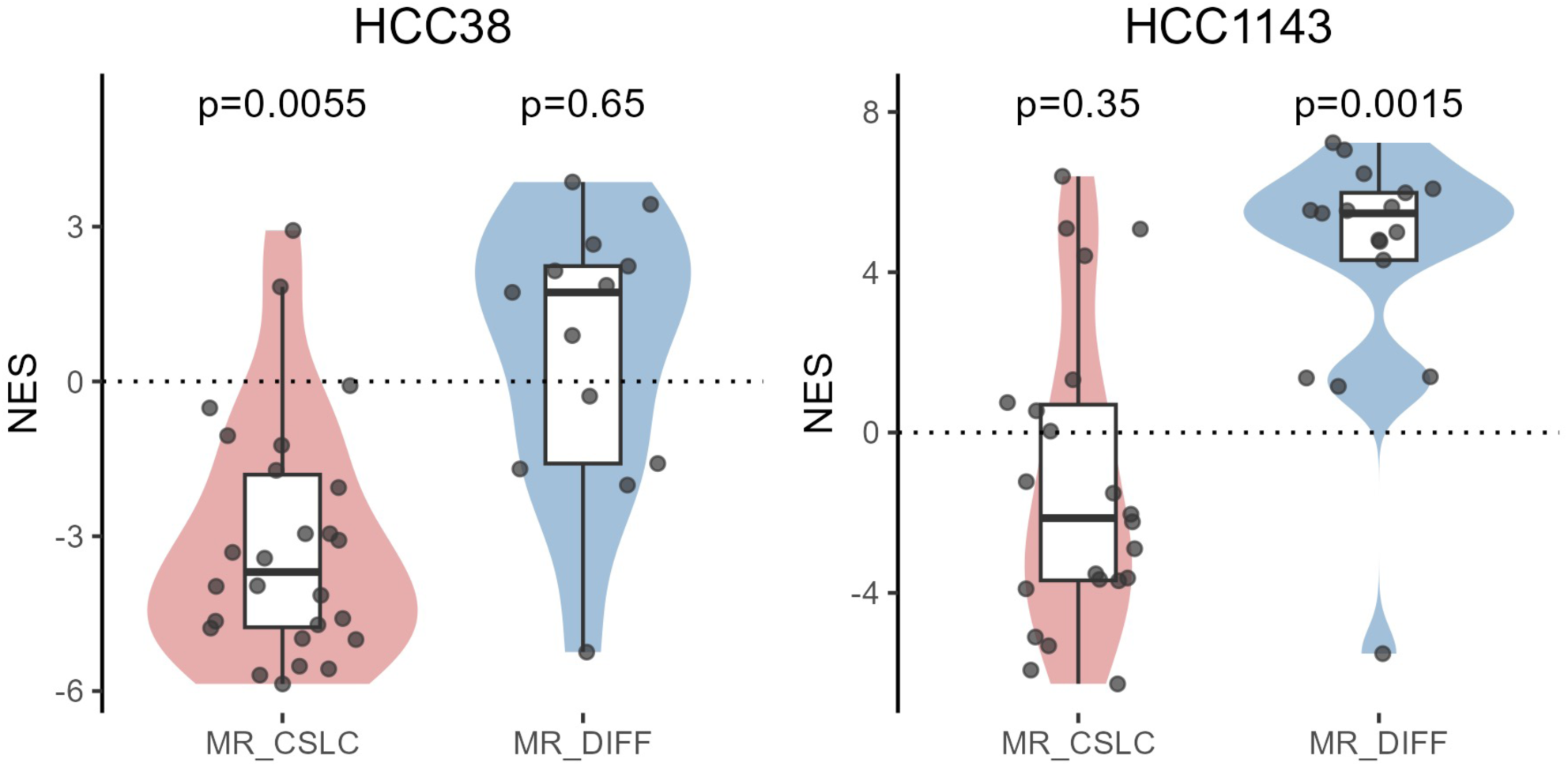
The enrichment of the gene set of stem cell process (supplementary table 2) between two groups (MR_CSLC_ and MR_DIFF_). Similar to Fig.5A, the enrichment score (NES) of stem cell process genes were significantly diminished (*p*=5.5×10^-3^) after MR_CSLC_ KO for HCC38, while NES was not significantly increased after MR_DIFF_ KO for the same cell line. On the contrary, for HCC1143, the increase of NES was more striking (*p*=1.5×10^-3^) after MR_DIFF_ KO, while the NES change was not significant for the group of MR_CSLC_ KO.

**Figure S23.**
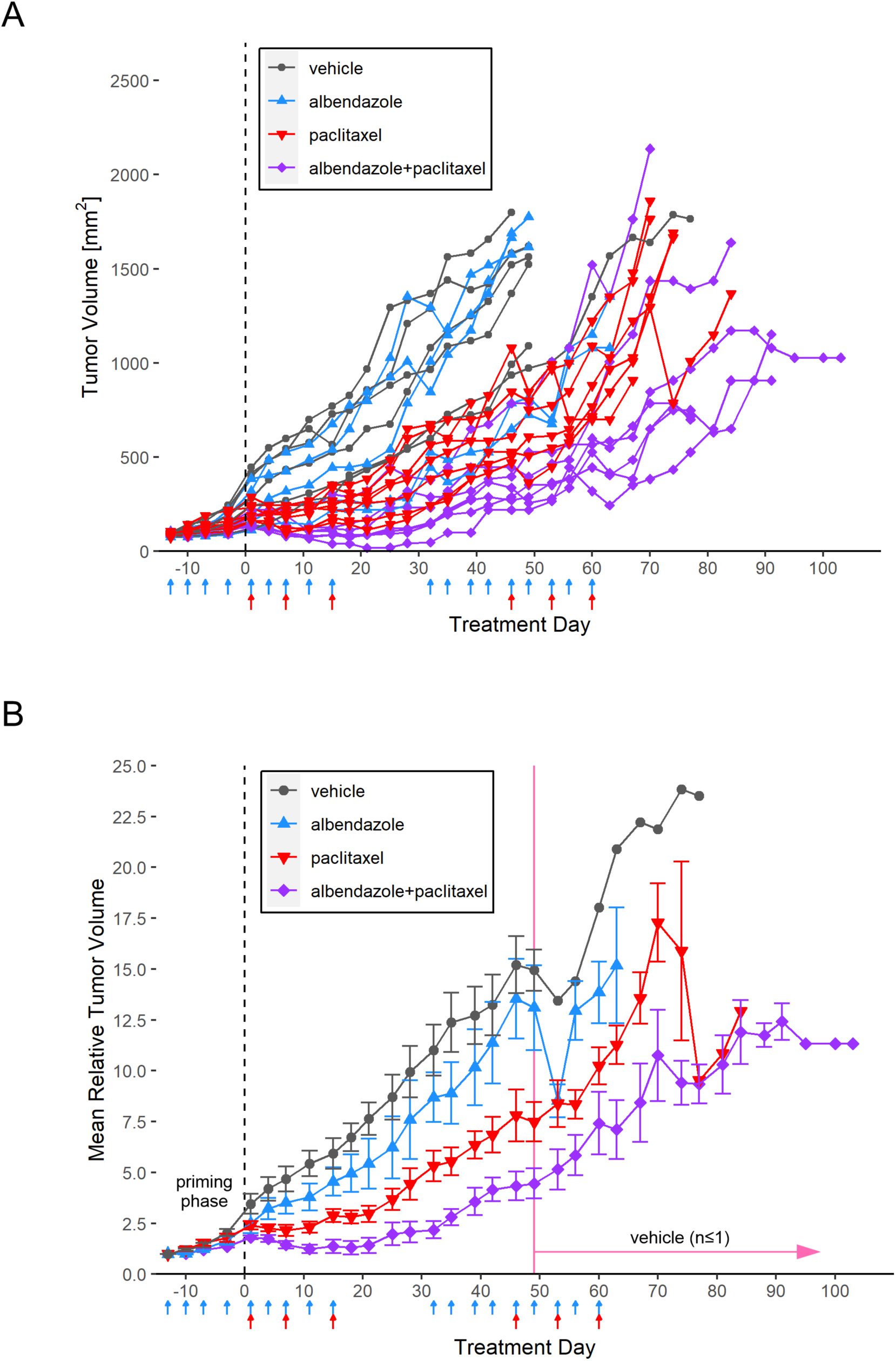
**A.** Spider plot of tumor volume measurements over time for individual mice. During a priming phase, mice were treated with albendazole 3 times weekly for two weeks before the start date of the combined drug therapy with paclitaxel, in order to sensitize the tumor cells. Mice with albendazole monotherapy were treated for the same amount of time as in the combination therapy. **B.** Mean relative tumor volumes over time. Tumor volumes are normalized by their volumes at Day-13. The error bar indicates one standard error of the mean. 5 out of 6 control mice (i.e. vehicle treatment) were euthanized before reaching Day 50.

**Figure S24.**
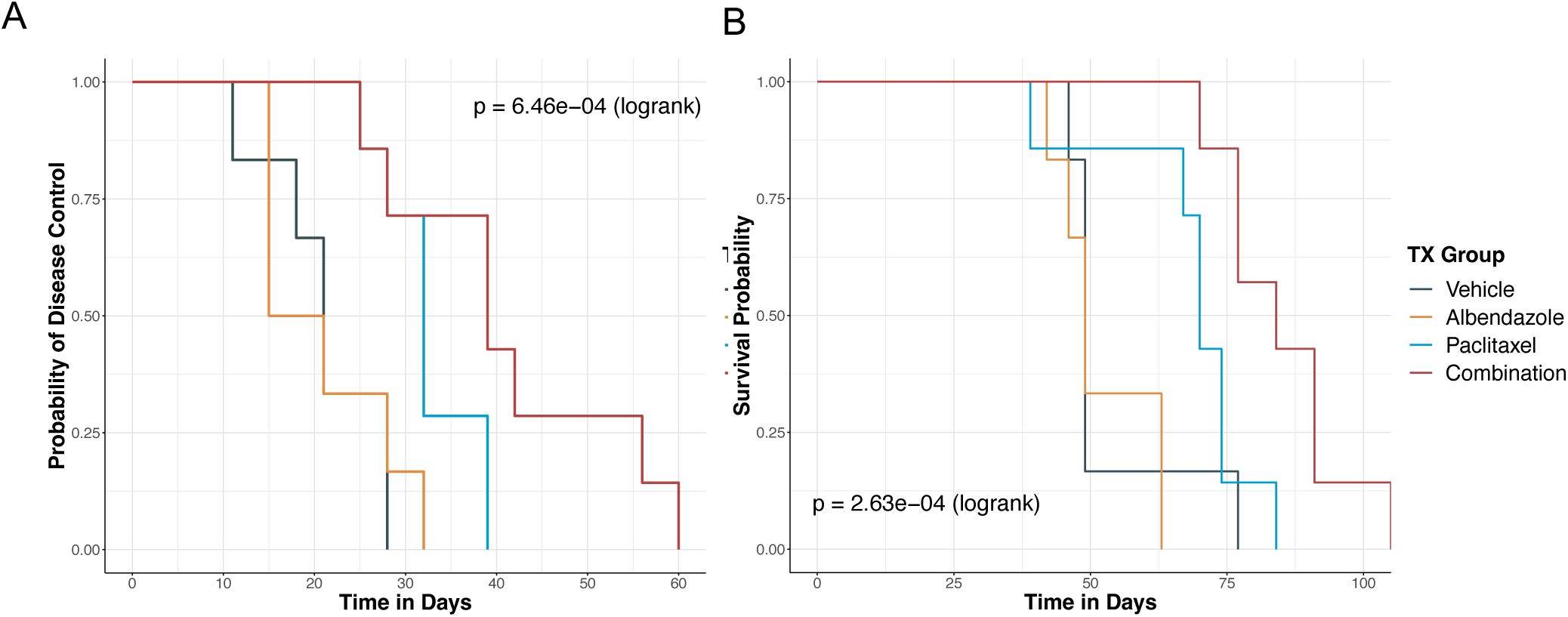
Kaplan-Meier analysis of the preclinical measurements for Disease Control (**A**) and survival (**B**) following treatment with albendazole and paclitaxel monotherapy vs. the combination.

